# Study of Motor Unit Action Potential Conduction Velocity and Firing Rate in Low Force Contractions using Empirical Mode Decomposition

**DOI:** 10.1101/2023.10.19.563177

**Authors:** Anton Dogadov, Jean-Yves Hogrel

**Affiliations:** Université Paris-Saclay, Centre National de la Recherche Scientifique, Paris-Saclay Institute of Neuroscience (NeuroPSI), Gif-sur-Yvette, France; BIOTN laboratory, Paris Est-Créteil Val de Marne University, Créteil, France; Neuromuscular Investigation Center, Institut de Myologie, GH Pitié-Salpêtrière, Paris Cedex 13, France

**Keywords:** Motor unit action potential conduction velocity, motor unit firing rate, ensemble empirical mode decomposition, surface electromyography

## Abstract

Arrays of surface electrodes may be used to record individual motor unit action potential (MUAP) trains from the skin surface. Processing of the later gives access to direct estimation of instantaneous conduction velocity and firing rate of individual MUAPs, which are informative physiological features. However, the estimation of conduction velocity and firing rate may be sensitive to additive noise, which is always present in surface EMG recordings. In this paper, we propose to use the Ensemble Empirical Mode Decomposition (EEMD) to represent individual MU instantaneous conduction velocity and firing rate time-series as a sum of components (intrinsic mode functions) and to omit the components with energy lower than the expected noise level. This approach enables to denoise the conduction velocity and firing rate time-sequencies, extracted from the surface EMG recordings, and to unmask a physiological relationship between them.

## 1. Introduction

Arrays of surface electrodes may be used to record individual motor unit (MU) action potentials (MUAP) from the skin surface [1–3]. Such recordings may provide information about individual motor unit conduction velocity (MU CV) and firing rate (MU FR), which are important physiological features. The MU CV depends on fiber properties, such as size of the fibers [4], their type [5], and external factors, such as temperature [6]. It is also known that MU CV and MU FR time-series are physiologically related [7–10], notably by the muscle velocity recovery cycle [11]. All in all, the observed values of MU CV and MU FR are nonstationary time-series, produced by physiologically related complex processes, influenced by a number of factors. However, it must be noticed that additive noise is always present in surface electromyographic (sEMG) recordings, which can deteriorate CV and, to a lesser extent, FR estimation.

Ensemble Empirical Mode Decomposition (EEMD) is a method for analyzing nonstationary data, which decomposes the signals using adaptive basis, derived from the data. Therefore, the signal components obtained from EEMD are usually expected to have physical meaning, which makes an EEMD a promising method in physiological studies. Moreover, this method does not require the time-series to have a constant time interval between two consecutive observations, which is the case in MU CV and FR time-series, in which a single observation corresponds to a single MUAP detection time, which has a nearly random nature.

In the current study, EEMD was applied to MU CV and FR time-series to

- omit the frequency components mostly influenced by additive noise;
- keep and analyze the significant remaining frequency components.

The results of this study may provide a tool for MU CV and MU FR time-series denoising and analysis.

## 2. Methods

### 2.1. Experimental protocol and setup

The signals were acquired in the Neuromuscular Physiology and Evaluation Laboratory at the Institute of Myology (Paris, France). The electrode system (Institute of Myology) was placed over the *abductor pollicis brevis* muscle, which was identified by palpation. The electrode system contained 11 electrodes (2 mm in diameter), grouped into three Laplacian arrays, or channels, a, b, c (Figure 1). A reference electrode was placed at the wrist.

**Figure 1.**
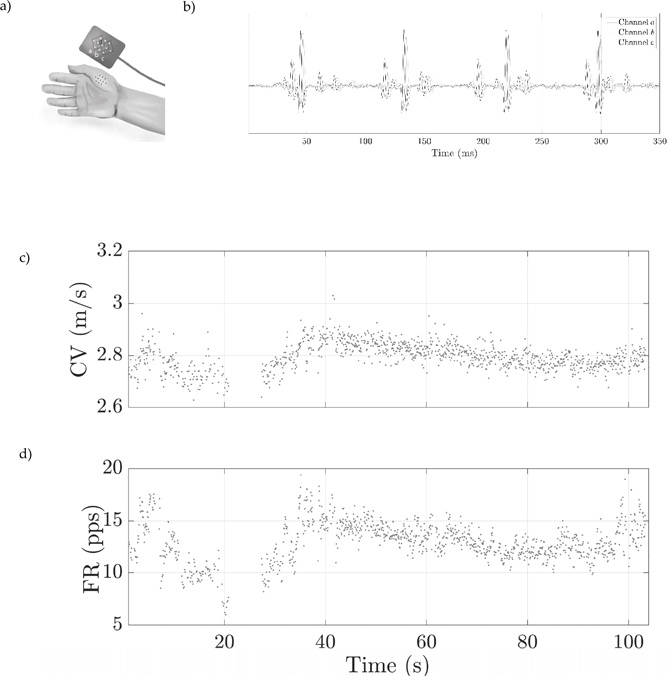
a) A view of electrode placement over the *abductor pollicis brevis* muscle. The electrodes were grouped by five into three Laplacian arrays (a, b, c). The distance between the electrodes d was 5 mm. b) Filtered EMG signal from a 3-channel Laplacian electrode; a dominant MUAP train may be observed. c) Raw estimated instantaneous conduction velocity (CV) of dominant MUAP d) Raw estimated instantaneous firing rate (FR) of dominant MUAP.

The subjects were asked to maintain a low force contraction such that only one dominating MUAP train was captured by the electrode system. The captured signals were provided to the subjects as the visual feedback. After the subjects succeeded in producing a regular MUAP train, they were subsequently asked to maintain the MUAP train during 100 s.

The analog processing of the signals consisted in band-bass filtering between 10 Hz and 1 kHz and amplification. Signals were sampled at 10 kHz using a National Instrument NI 6221 data acquisition module.

Fifteen volunteers with no prior knowledge of trauma or neuromuscular disease participated in experiments. Informed consent was obtained from all participants. The signals from ten subjects were selected for further analysis. The selection criterion was the presence of a single dominant MUAP train in the recordings, i.e., among 15 volunteers, only 10 were able to maintain a steady-state firing rate of one dominant MU.

### 2.2. MUAP detection

The MUAPs were detected in sEMG signals using continuous wavelet transform by Mexican hat wavelet. Each detected MUAP was then interpolated by a second-degree polynomial function in 7-point window (0.7 ms at 10 kHz sampling frequency) centered on the pick value of the MUAP. The MUAP detection time were calculated in each channel from the polynomial coefficients as a time coordinate of the polynomial function extremum. If several MUAP train were detected in the signal, only the MUAPs from the dominating MU were analyzed. A visual verification was performed to ensure that all detected MUAPs correspond to the same MU.

The signal-to-noise ratio (SNR) of the recorded MUAP train was calculated in each channel as ten logarithms of the ratio of signal power to the power of noise. The power of signal was calculated among all 9 ms windows, containing MUAPs of the motor unit of interest; the power of noise was calculated among all the windows, which did not contain any MUAPs (from MU of interest or any other MUs). The SNR was calculated separately in all channels, used for estimating CV and FR, and the minimal value of the two was retained.

### 2.3 .Conduction velocity and firing rate estimation

For each detected MUAP of the MU of interest, local measure of conduction velocity was calculated as the interelectrode distance *d* divided by the travel times of the *i*-th MUAP between two channels, *a* and *b* or *b* and *c* (Figure 1a). For example, the MUAP propagation velocity, calculated between channels *a* and *b* was:

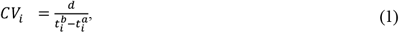

where the distance *d* was equal to 5 mm (Figure 1a).

Therefore, the instantaneous MU CV of the MU of interest may be calculated either between the first and the second channel (*a* and *b*) or between the second and the third one (*b* and *c*). Between these two possibilities, we chose that pair of channels, in which the mean CV value, calculated over all detected MUAPs of one subject was lower.

For each detected MUAP *i*, local measure of the firing rate in a given channel was calculated from the delay between the appearance time of the previous and the following MUAPs (*i*-1 and *i+1*), detected in this channel. For example, for the channel *a*, the local measure of the instantaneous MU FR was:

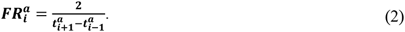

Finally, the instantaneous MU FR of the MU of interest ***FR***_***i***_ was calculated as an average of the local measure of the firing rate 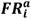 and 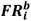 in channels *a* and *b*, if CV was calculated between *a* and *b* channels, otherwise ***FR***_***i***_ was calculated as an average of 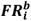 and 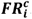. Therefore, for each detected MUAP, a CV and instantaneous FR was estimated, forming ***CV***_***i***_ and ***FR***_***i***_ time-series with a number of samples, equal to the number of detected MUAPs minus two.

### 2.4. Ensemble empirical mode decomposition

Empirical mode decomposition (EMD) is a method, proposed by Huang et al. [12], which consists in decomposition of time-series ***x*** into a sum of intrinsic mode functions (IMFs) denoted as ***c***_***m***_:

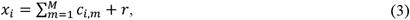

where ***r*** is a residual of EMD decomposition, and ***x***_***i***_ is a time-series, which may be ***CV***_***i***_ or ***FR***_***i***_.

The first IMFs (*m*=1) is the IMF with the highest mean frequency, and the last IMF (*m*=*M*) is the IMF with the lowest mean frequency. Huang et al. [12] considered IMFs as the adaptive basis, as it is not *a priori* known and is derived from the data. Due to this fact, a waveform of an IMF is usually expected to have a physical meaning, in contrast to Fourier transform, which basis consists of harmonic functions. One of principal drawbacks of EMD is a mixing problem. This problem is related to the fact that one IMF of EMD decomposition may consist of signals of widely different scales [13]. This problem usually appears when the signal is disturbed by noise or contains intermittencies, because the EMD is principally based on extrema distribution. Therefore, because of this problem, the results of EMD decomposition are sensitive to noise.

Ensemble empirical mode decomposition (EEMD) is a noise-assisted version of EMD [13], which is less sensitive to a mixing problem. The EEMD is an iterative procedure, which consists in adding synthetized white noise to the analyzed time-series and performing a decomposition (3) of the time-series, perturbed by the added noise [13]. The procedure is repeated for some predefined number of iterations with different white-noise signals generated for each iteration. The result of EEMD was calculated as an average of IMFs obtained at all iterations.

The time-series of CV and FR of the MUAPs were used to perform an EEMD-decomposition of the CV and FR time-series into intrinsic mode functions. EEMD was performed with 300 iterations. The standard deviation of white noise, added to CV and FR time-series was 0.2 of the CV and FR initial time-series standard deviation, based on the study of Wu and Huang [13], who concluded that the optimal number of iterations is few hundreds and the optimal deviation of the standard deviation of the added noise is 0.2 of the standard deviation of the analyzed data. The EEMD decomposition was performed using the hht package for R [14].

### 2.5. Simulation of the additive noise

Since both instantaneous CV and FR were calculated based on MUAP detection over time, the error in MUAP detection leads to an error in CV and FR time series. In turn, MUAP detection precision depends on signal-to-noise ratio (SNR) in sEMG signals.

To study the influence of raw EMG signals SNR on CV and FR, we simulated a noisy MUAP waveform by summing a single simulated MUAP waveform with randomly generated noise. A time-location of the maximum of MUAP waveform 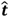 was *a priori* known from the original simulated MUAP waveform. The MUAP waveform was summed with colored Gaussian noise, created by filtering a white Gaussian noise with a low-pass FIR-filter of the order 41. The coefficients of the filter were calculated by sampling the noise spectrum observed in one-second EMG recordings in absence of muscle activity. The filter output was multiplied by a weight coefficient to represent a required noise level and was added to a MUAP waveform. For each noise level, a SNR was calculated as 10 logarithms of the ratio between the energy of the MUAP waveform and the added simulated noise. A MUAP maxima was detected in noisy signal, as explained in MUAP detection subsection, and a detection error 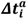 was found as a difference between the true maxima time-location of the MUAP waveform 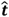 and a detected time-location of the MUAP maxima in presence of noise 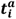:

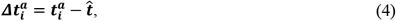

where ***i*** = **1**… ***N*** is a number of simulation.

A detection error in two channels, *a* and *b* was simulated. For each of these two channels and for each noise level, N=20,000 simulations of random noise were performed. As a result of simulation at a given noise level, simulated time-series of MUAP detection error in channels *a* and *b* (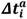, 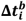 correspondingly) were obtained. In our study, the properties of both channels were considered the same, therefore 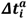 and 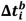 were independent and identically distributed.

The absolute error on conduction velocity estimation 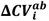, calculated from the MUAP travel time between the channels a and b, was calculated as:

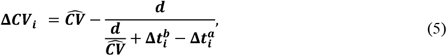

where 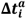 and 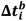 are simulated detection errors, *d* – interelectrode distance (5 mm) and 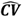 is a mean conduction velocity.

The absolute error of instantaneous firing rate estimation 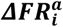, calculated from the delay between the detection of two sequential MUAPs in channel *a*, was calculated as:

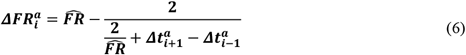

Where 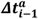, 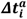 and 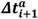 are independent and identically distributed variables, generated at steps *i*-1, *i*, and *i*+1, and 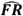 is a mean firing rate.

An error of the averaged value of the local FR in channels a and b was:

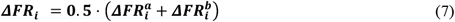

In such a way, for each SNR level from 10 to 70 dB, characterizing the quality of sEMG signals, ***ΔFR***_***i***_ and ***ΔCV***_***i***_ sequences were simulated. The standard deviation of the sequences ***σ***_Δ***CV***_ and ***σ***_Δ***FR***_ was calculated at each SNR level. An exponential relationship was proposed to link the SNR of raw sEMG signals and standard deviation of errors ***σ***_Δ***CV***_ and ***σ***_Δ***FR***_:

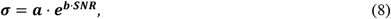

where *a* and *b* were the identified parameters and ***σ*** stands for ***σ***_Δ***CV***_ or ***σ***_Δ***FR***_.

### 2.6. Study of the information content of IMFs of the CV time-series

Due to random errors ***ΔCV***_***i***_ and ***ΔFR***_***i***_, MU CV and MU FR time-series are influenced by noise, related to MUAP detection error. We assume that this noise is white additive Gaussian noise, therefore, both estimated CV and FR time series may be represented as

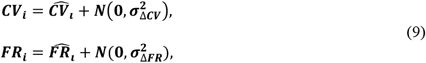

where 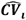 and 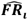 are real values of conduction velocity and firing rate for *i*-th MUAP, and ***N***(**0, *σ***) is white Gaussian noise with zero mean and standard deviation ***σ***_Δ***CV***_ and ***σ***_Δ***FR***_, for MUAP CV and FR time-series correspondingly.

Wu and Huang [15] have shown that the expectation of natural logarithm of the energy ***E***_***m***_ of m-th IMF (m ≥2) from a decomposition of a white noise time-series with unitary power is related with the expectation of a logarithm of its averaged period ***T***_***m***_ by the following equation:

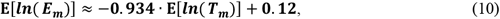

where the averaged period of m-th IMF ***T***_***m***_ is estimated as number of samples in time-series divided by a number of maxima.

Wu and Huang [15] also demonstrated that the *j*-th percentile of the distribution of the energy logarithm of *m*-th IMF of white noise time-series with N samples and mean period ***T***_***m***_ may be found as:

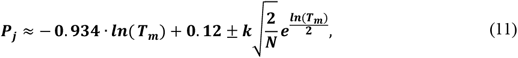

where k= 2.326 for 5^th^ (-) and 95^th^ (+) percentiles.

The equations (10) and (11) assume that the white noise time-series has unitary power. To take into account a power of noise, a term **In**(***σ***^**2**^) was added to these equations, where ***σ***^**2**^ is a variance of the white-noise. Therefore, the equation (11) for 95 ^th^ percentile becomes:

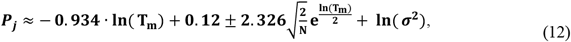

where **In**(***σ***^**2**^) is a logarithm of noise variance in MUAP CV 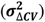 or FR 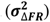 time series, which was found using simulation as explained in subsection “Simulation of the additive noise”.

To study the information content of IMFs of CV time-series, an SNR of raw sEMG signals was calculated for each subject. Then, using (9) a variance of noise in CV 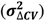 and FR 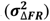 time-series was estimated for each SNR level. After EEMD decomposition of CV and FR time-series of each subject, the energy of each IMFs from CV or FR decomposition were estimated and compared with 95^th^ percentile of the distribution of energy of IMFs of white noise with a variance 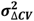 or 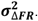. Only such IMF components were considered significant, which were located above the 95 ^th^ percentile.

## 3. Results

Bellow we present the results for a typical subject (#1). The results from the other subjects are presented in Annex 2.

### 3.1. Recorded signals

A 350 ms window of filtered sEMG signal from the subject #1 is shown in Figure 1b. A dominant MUAP train may be seen from the signals. Instantaneous CV and FR of the dominant MUAP were estimated, forming CV (Figure 1c) and FR (Figure 1d) time series. For the subject #1, the minimal SNR was equal to 27.2 dB among the channels, in which CV and FR were calculated.

### 3.2. EEMD

EEMD was applied on estimated MU CV and MU FR time-series to separate the time-series into IMFs. Normalized IMFs of MU CV (blue) and MU FR (black) time-series decomposition is shown in Figure 2a, and the residuals are shown in Figure 2b.

**Figure 2.**
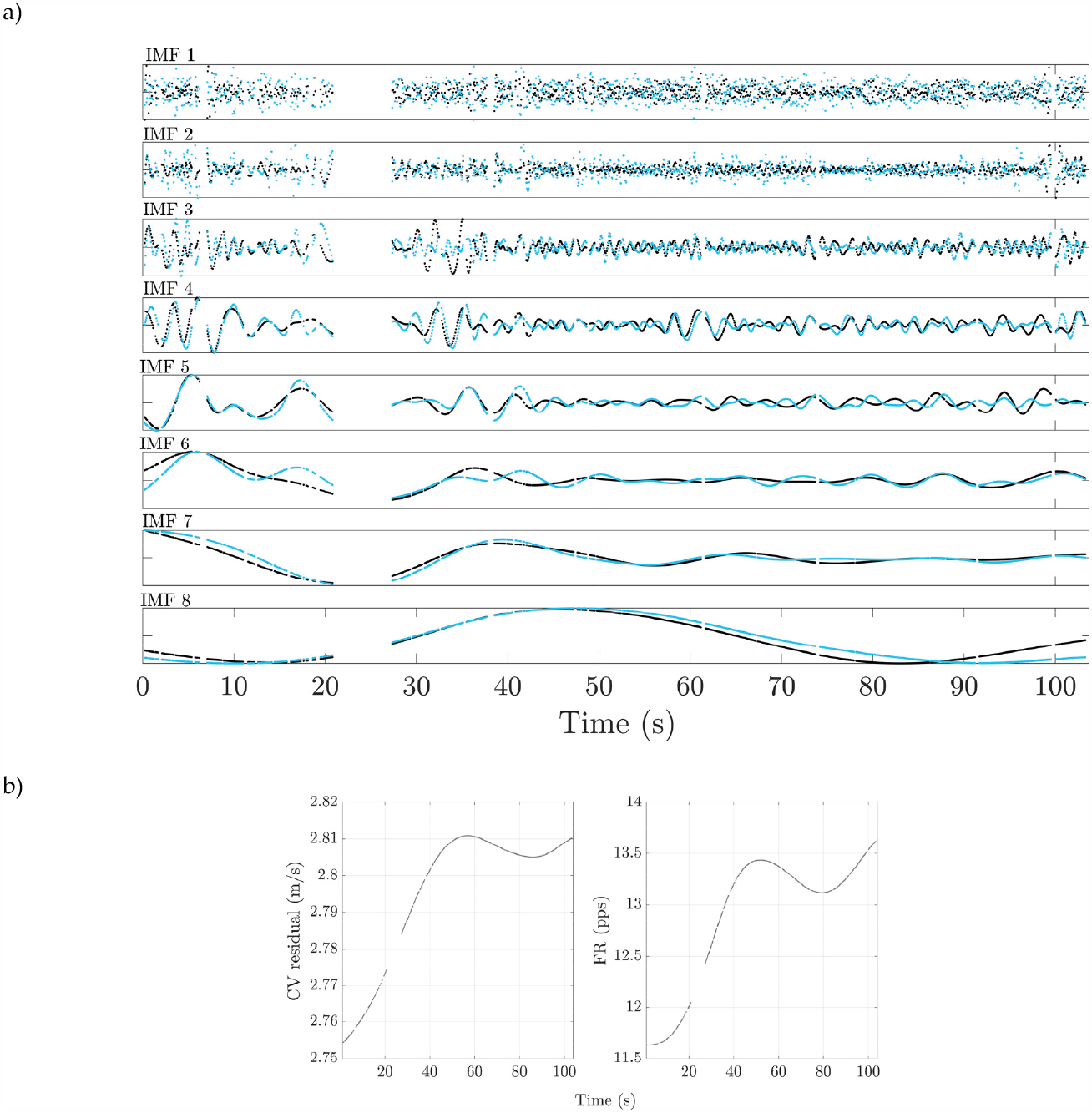
Normalized IMFs from CV (blue) and FR (black) time-series decomposition and residuals (b) (subject #1).

The mean frequency of IMF components decreases from the top to the bottom in the Figure 2a. A similarity may be noticed between corresponding low-frequency IMF components of CV and FR time-series decomposition, as well as between the residuals.

### 3.3. Simulation of the additive noise

Figure 3 shows the influence of the raw sEMG signal SNR on the standard deviation of the error in MU CV and MU FR, obtained from simulated data and the mean CV and FR values equal to 3 m/s and 10 pps correspondingly.

**Figure 3.**
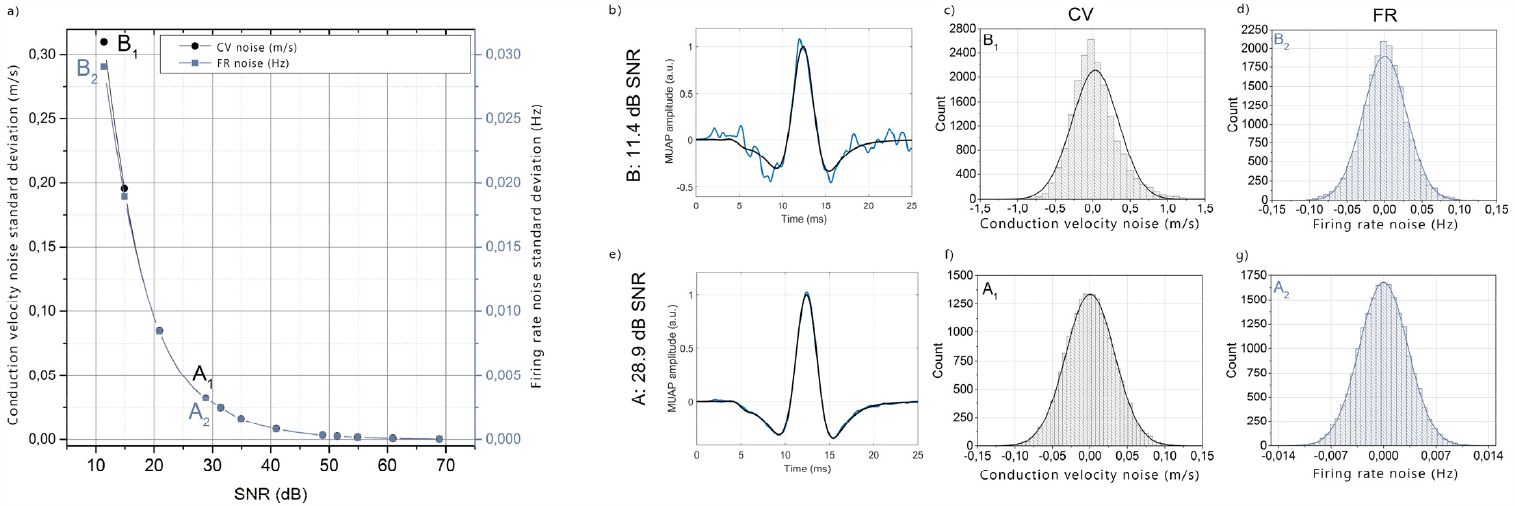
The influence of the signal-to-noise ratio in raw EMG signals on standard deviation of the error in CV and FR time series (a). The estimation was performed for mean CV and FR values equal to 3 m/s and 10 pps. Points A1 and A2 correspond to 28.9 dB SNR and points B1 and B2 correspond to 11.4 dB SNR. Panels b and e show the initial simulated MUAP waveform (black) and the MUAP waveform with added noise (blue) at 11.4 dB (b) and 28.9 dB (e) Histograms of the standard deviation of the error in CV time-series are shown for 11.4 dB (c) 28.9 dB (f). Histograms of the standard deviation of the error in FR time-series are shown for 11.4 dB (d) 28.9 dB (g).

Figure 3b and e show a MUAP with additive white Gaussian noise (blue) at different signal-to-noise ratio (Figure 3b at 11.4 dB, Figure 3e at 28.9 dB). The histograms of the disribution of error in CV time-series, extracted from sEMG signal with SNR of 11.4 dB and 28.9 dB are shown in Figure 3c and f correspondingly The histograms of the disribution of error in FR time-series, extracted from sEMG with SNR of 11.4 dB and 28.9 dB are shown in Figure 3d and g correspondingly. The pannel a show that the standard deviation of error in CV time-series (left axis) and FR time-series (right axis) increases when the SNR decreases. The standard deviation of error in CV time-series was 0.03 m/s and 0.31m/s for SNR of 28.9 dB and 11.4 dB correspondingly (points A1 and B1). For the same SNR values, the standard deviation of error in FR time-series was 0.003 Hz and 0.029 Hz (points A2 and B2). The disnormality of simulated data also increased for both CV and FR error with the noise power: for two referred SNR values, the kurtosis of CV estimation error was 3.10 and 6.31, and the kurtosis of the FR estimation error was 3.04 and 3.66. Moreover, the distribution of the CV estimation error corresponding to 11.4 dB was skewed and a “tail” of overestimated CV values appeared. For two referred SNR levels, the skewness of the CV estimation error was 0.08 and 1.01. To study the whiteness of simulated data, a Box-Ljung test was applied [16]. The null-hypotheses about the whiteness was accepted for both estimated CV and FR error (p=0.13 and p=0.15 correspondingly). The relationships between the sEMG SNR and standard dviation of error in CV and FR time-series were approximated by exponential functions, shown by solid lines in Figure 3a and defined by the equation (**9**). The identified function parameters where *a* = 1.379 and *b* = -0.131 for the noise in CV time-series; *a* = 0.121 and *b* = -0.125 for the error in FR time-series. The *R*^2^ value was 0.99 in both cases. For the subject #1 (sEMG SNR 27.2 dB) the standard deviation of error in CV and FR time-series, calculated with (8), was 0.04 m/s and 0.004 pps correspondingly. However, the error of CV and FR estimation in presence of additive noise in EMG recordings depends correspondingly on the mean values of CV and FR (5-7). This dependence is particulary significant for error of CV estimation. Annex 1 shows the values of parameters for different values of averaged CV.

### 3.4. Study of the information content of IMFs of the CV time-series

Figure 4 shows enrergy and mean period of IMFs of CV and FR time-series. Each IMF component, from 2 to 8 is shown by a blue cross. The energy of the first IMF is not shown, as (10) holds only for m≥2. The abscissa shows a natural logarithm of energy of each IMF component, the ordinate axis shows the natural logarithm of a mean period of IMF components, calculated as a total number of MUAPs divided by number of local maxima in this IMF. As the mean period is expressed in number of MUAPs per period and not in seconds, the second ordinate axis is added to give a period duration in seconds. The values were obtained by multipling the period in MUAPs/cycle, divided by the averaged firing rate.

**Figure 4.**
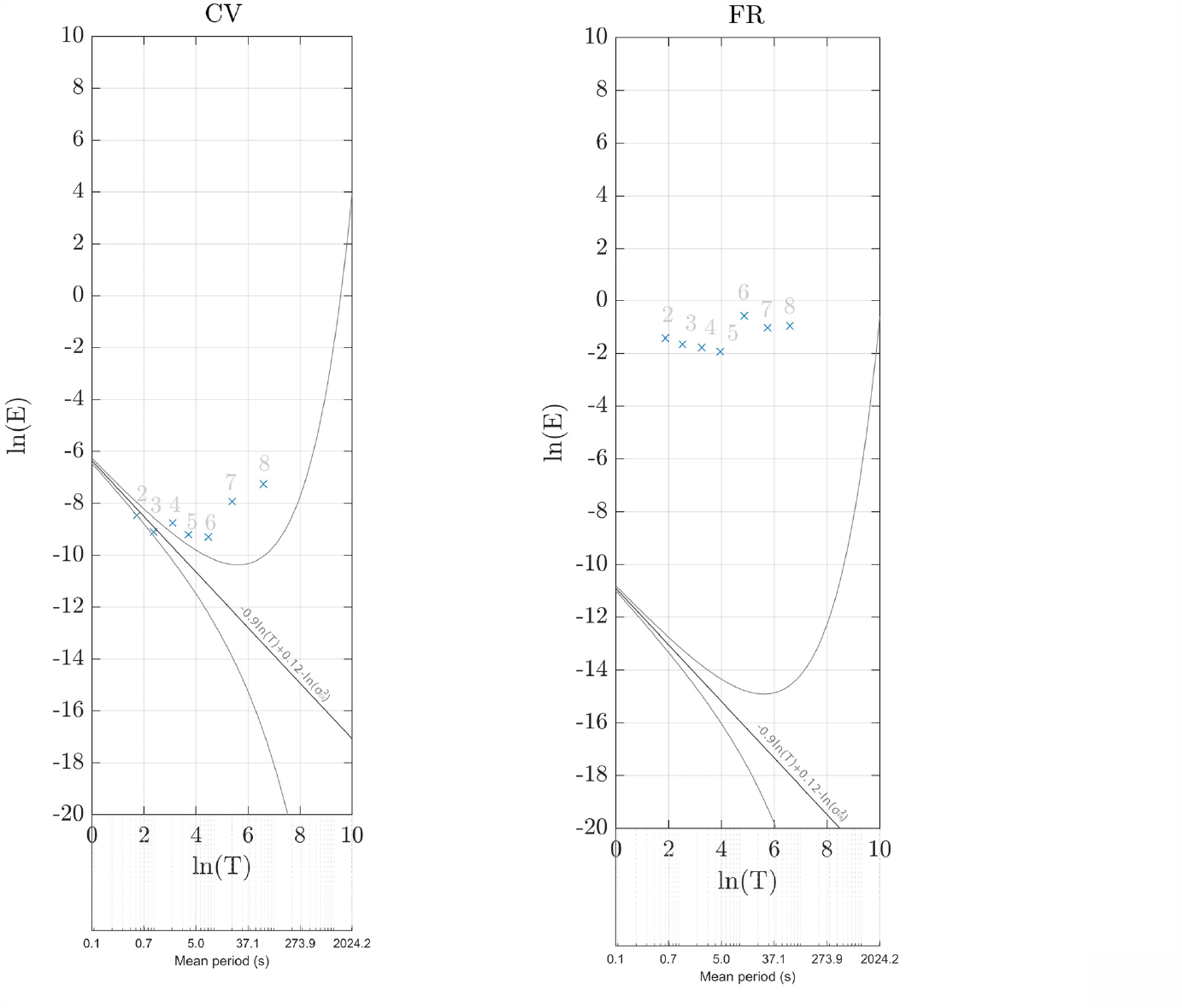
Spectrum of IMF components of CV and FR time-series decomposition. Logarithms of energy and mean-period of each component from 2 to 8 are shown by blue crosses. A mean, 5^th^ and 95^th^ quantile of noise distribution are identified on the same plot by black lines.

The same figures show the expected mean, 5^th^ and 95^th^ percentile of the distribtion of IMFs of the decomposition of white Gaussian noise time-series with the same number of samples, which was in the CV ans FR MUAP time-series and with the standard deviation of 0.04 m/s and 0.004 pps correspondingly.

It may be seen from Figure 4a that the 2^nd^ and th 3^rd^ IMFs components of CV time-series decomposition may be higly influenced by noise, related to additive noise in sEMG recordings. Indeed, these components are located below the 95^th^ percentile of the noise distribution.

## 4. Discussion

This study has shown that the EEMD might be a powerful method for analyzing noisy time-series of single motor unit CV and FR. An additive noise in surface EMG recordings is an issue, deteriorating CV estimations. An important noise level in EMG recordings may mask physiological information in estimated CV time-series.

In the current study, we proposed a relationship to link the error in CV and FR time-series with the signal-to-noise ratio in sEMG signals, from which these time-series were extracted. It was shown that the CV estimation error distribution, introduced by additive noise in sEMG recording, is close to white Gaussian noise. Therefore, CV and FR errors are distributed among all frequencies. The method proposed by Huang may be used to quantify the noise distribution among the IMFs components of CV and FR time-series decomposition. The IMF components of CV and FR time-series decomposition may be compared with the noise distribution 95^th^ percentile to identify the significant components. Figure 5a shows raw MU CV time-series, and Figure 5b shows with the same time series, from which the IMF components IMF_m_, m=1,2,3, influenced by additive noise, were removed. Omitting the noisy components, unmasks the relationship between CV and FR time-series, which may be hidden by noise. Figure 5c shows time-dependent cross-correlation between raw MU CV and MU FR time-series and Figure 5d shows time-dependent cross-correlation between mid-frequency components (sum of IMFm, m=4,5) of MU CV and MU FR time-series decomposition. The high frequency components (m=1,2,3) were omitted as these components in CV time-series decomposition can be strongly influenced by noise. The low frequency components were omitted as their period is long (more than 100 samples) compared to analyzed window size. Time-dependent cross-correlation plots were built using window size t_d_ = 101 samples and the maximum lead–lag window τ = 20 MUAPs.

**Figure 5.**
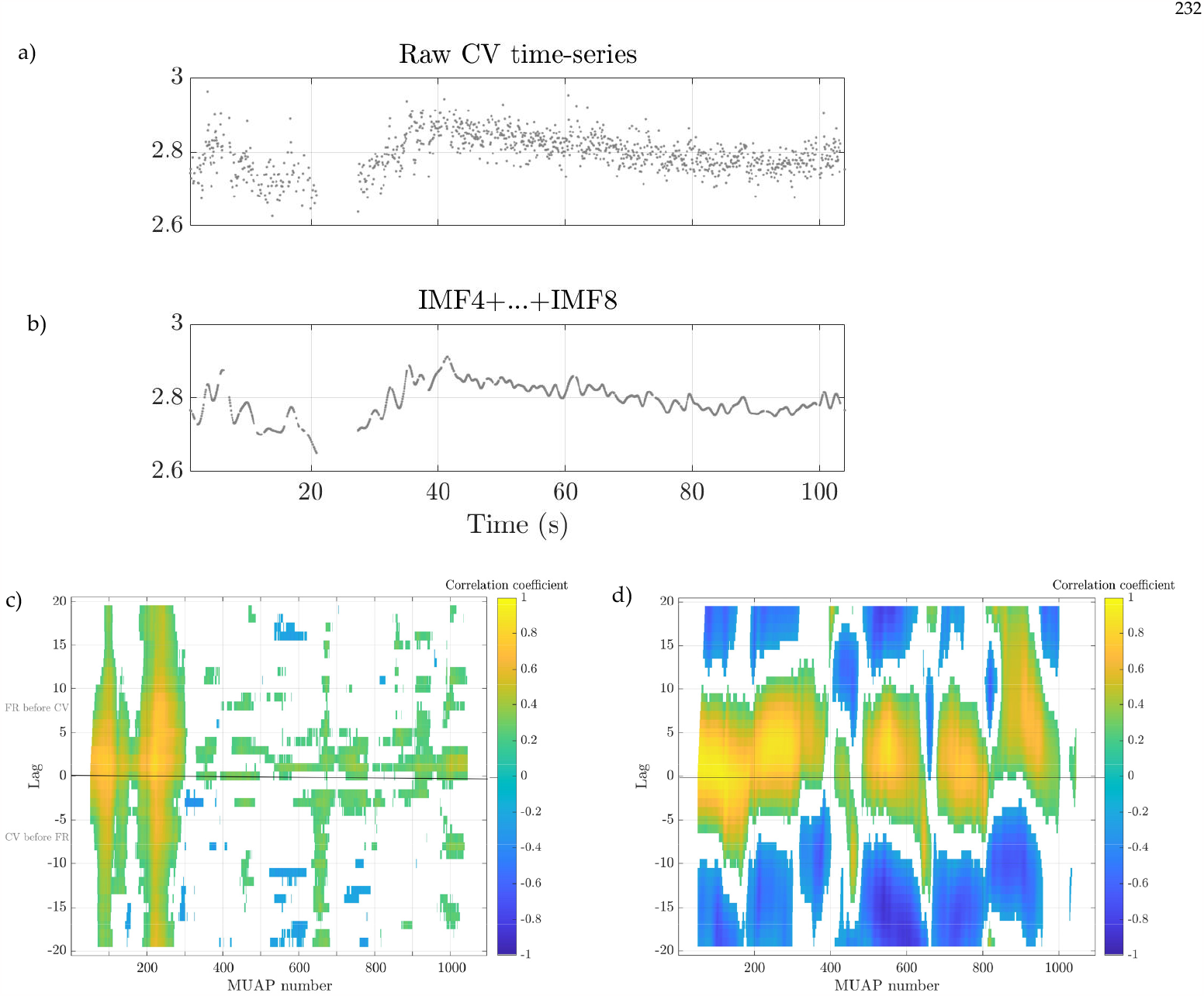
a) Raw MU CV time-series from subject #1. (b) Mid- and low-frequency components IMF4…IMF8 of EED decomposition of MU CV time-series from subject #1. c) Time-dependent cross-correlation plots between raw MU CV and MU FR time-series. d) Time-dependent cross-correlation plots between mid-frequency components of CV and FR decomposition (IMF4 and IMF5). In panels c and d, the correlation coefficient value is shown by color; white color corresponds to the time-windows, for which the correlation coefficient was not significant (p>0.05)

Each point of the Figure 5c and Figure 5d represent the correlation coefficient between time windows from CV and FR time series, where the axis of the figures determine location of the window in time-series and the lag between them. The color shows the correlation-coefficient value, and the white color corresponds to the time-windows, for which the correlation coefficient was not significant (p<0.05) The cross-correlation plot between raw CV and FR time-series (a) shows high correlation at the beginning of the time-series. However, the correlation stays insignificant most of the time, which might be explained by noise. Computing cross-correlation plot only between the mid-frequency components of CV and FR EEMD decomposition makes the cross-correlation between CV and FR visible during the whole MUAP sequence. It may be seen that CM time-series lags behind the FR time-series. The mean lag between mid-frequency components of CV and FR time-series was about 2 MUAPs, which was calculated as the location of cross-correlation maxima, averaged along all time-windows (abscissa axis).

### 4.1. Limitations and perspectives

In the current study, the proposed method was applied on low-threshold MUs of subjects, performing low-force activations of the *abductor pollicis brevis* muscle. The method may be combined with MUAP decomposition algorithm to study multiple MUAP trains from muscle activations at a higher force level.

The proposed method may be also used to also study sustained fatiguing muscle activity. In such a case, an adaptive temperature control system may be worthwhile to be implemented, as in [4]. In the current study, as the measurement was performed during 100 s only, no temperature control system was used.

Finally, in the current study, it was shown that EEMD decomposition of MU CV and MU FR time-series help revealing the relationship between them. As a further work, EEMD decomposition may be used to study the influence of multiple factors on CV.

## Supporting information

Annex 1. Error of CV estimation as a function of SNR level in raw EMG recording and the mean value of MU CV

Annex 2. Analysis of CV and FR time-series

