## Supplementary material for "Study of Motor Unit Action Potential Conduction Velocity and Firing Rate in Low Force Contractions using Empirical Mode Decomposition": Annex 1. Error of CV estimation as a function of SNR level in raw EMG recording and the mean value of MU CV

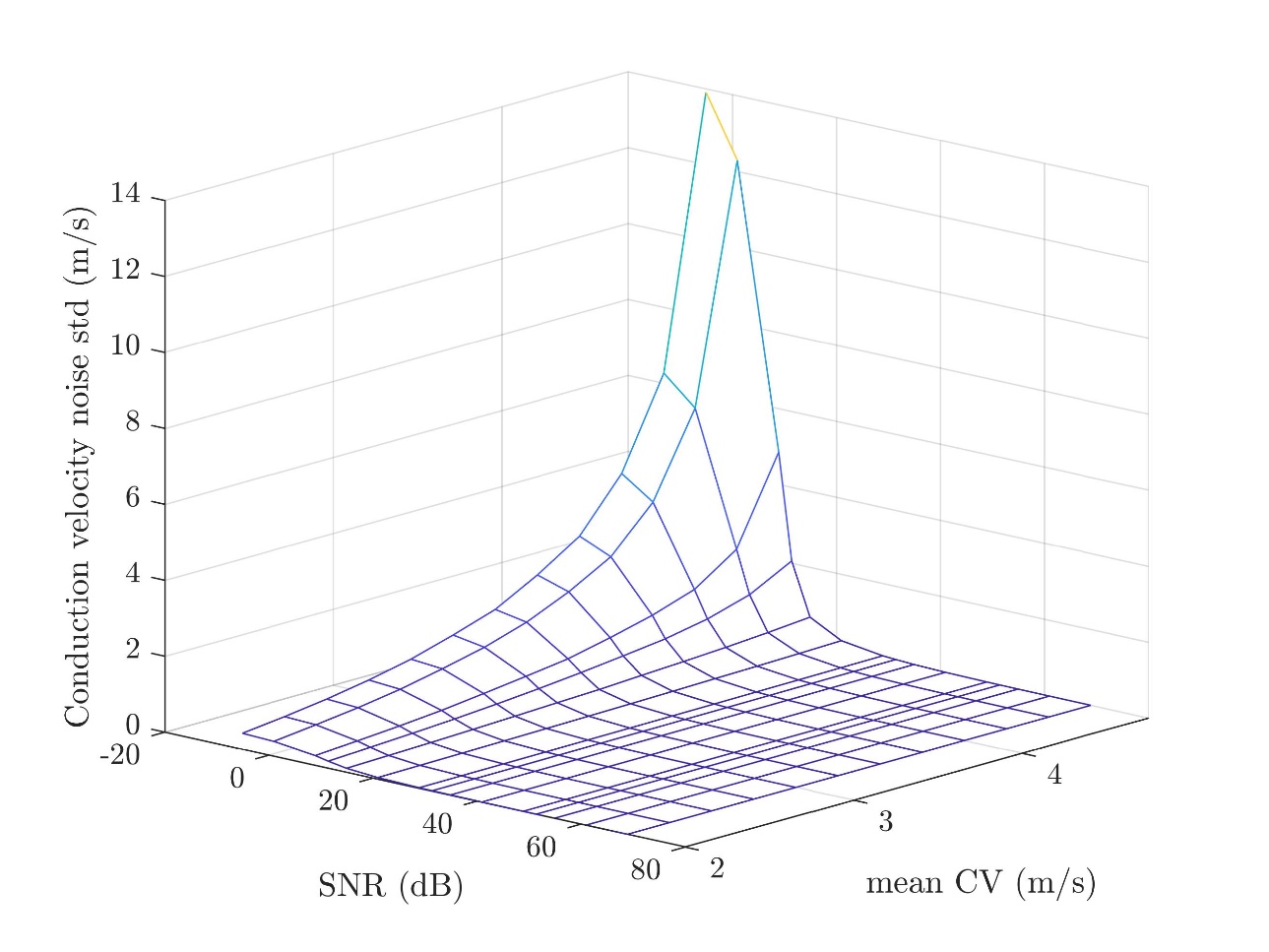

An exponential relationship the SNR of raw sEMG signals and standard deviation of errors $\boldsymbol{\sigma}_{\boldsymbol{\Delta CV}}$:

$$\boldsymbol{\sigma}_{\boldsymbol{\Delta CV}}\mathbf{=}\boldsymbol{a}\boldsymbol{\cdot}\boldsymbol{e}^{\boldsymbol{b}\boldsymbol{\cdot}\boldsymbol{SNR}}\boldsymbol{,}$$

The value of parameters *a* and *b* depends on mean CV:

| Mean CV (m/s) | *a* | *b* | *R*^2^ |
| --- | --- | --- | --- |
| 2,00 | 1,013 | -0,1492 | 0,9983 |
| 2,25 | 1,347 | -0,1516 | 0,9982 |
| 2,50 | 1,725 | -0,1541 | 0,9975 |
| 2,75 | 2,151 | -0,1552 | 0,9977 |
| 3,00 | 2,756 | -0,1592 | 0,9973 |
| 3,25 | 3,551 | -0,1644 | 0,9969 |
| 3,50 | 4,756 | -0,1739 | 0,9953 |
| 3,75 | 5,951 | -0,1777 | 0,9957 |
| 4,00 | 7,397 | -0,1826 | 0,995 |
| 4,25 | 11,17 | -0,2006 | 0,9937 |
| 4,5 | 19,99 | -0,2294 | 0,9936 |
| 4,75 | 51,3 | -0,2826 | 0,9921 |
