## Supplementary material for "Study of Motor Unit Action Potential Conduction Velocity and Firing Rate in Low Force Contractions using Empirical Mode Decomposition": Annex 2. Analysis of CV and FR time-series

Subject 1-4, 8, 10-15 could maintain a single dominant MUAP train during the recording. The results of these subjects are presented below.

Subject 1

Raw CV and FR time series:

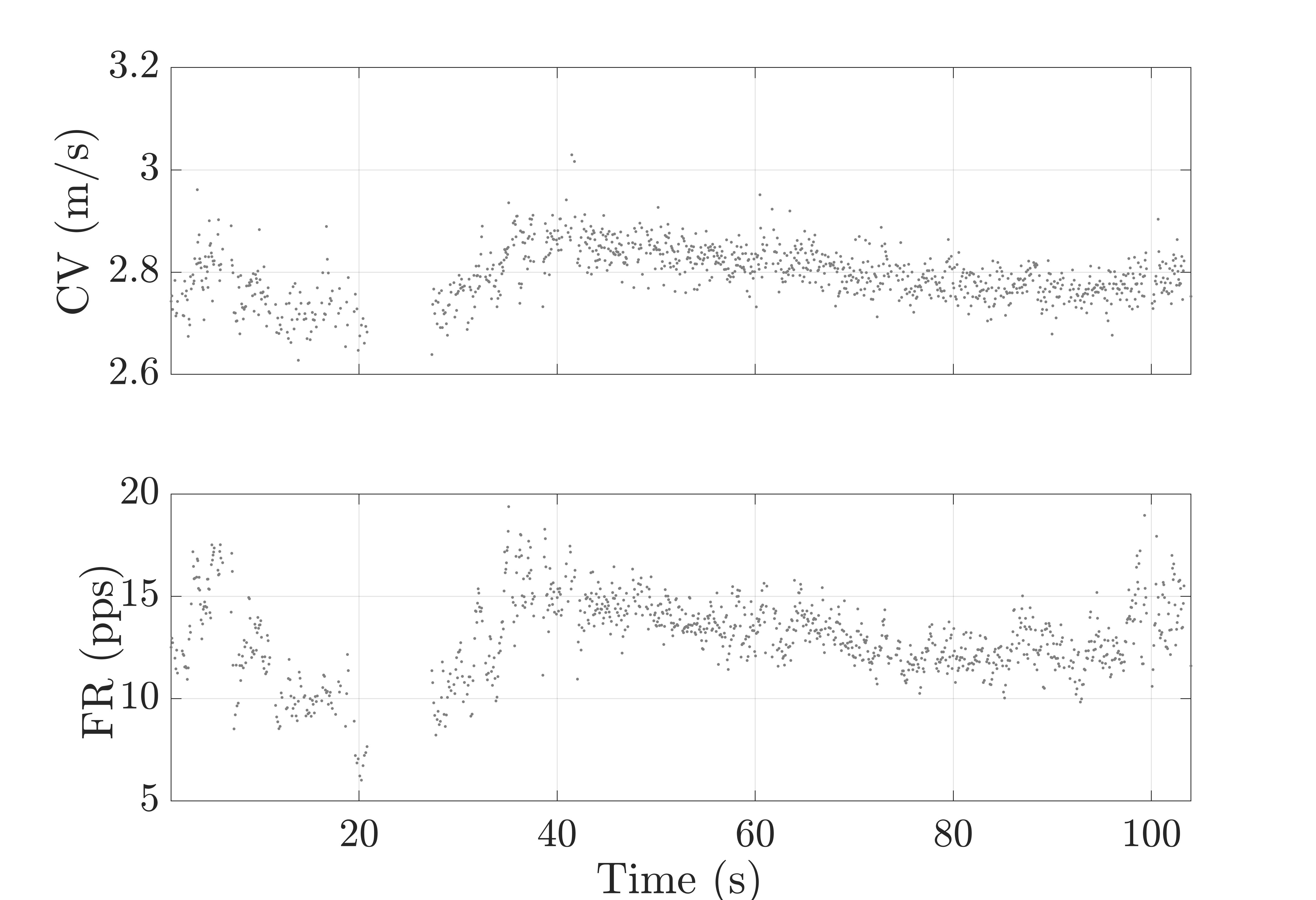

IMFs and residuals of EEMD decomposition of CV and FR time-series:

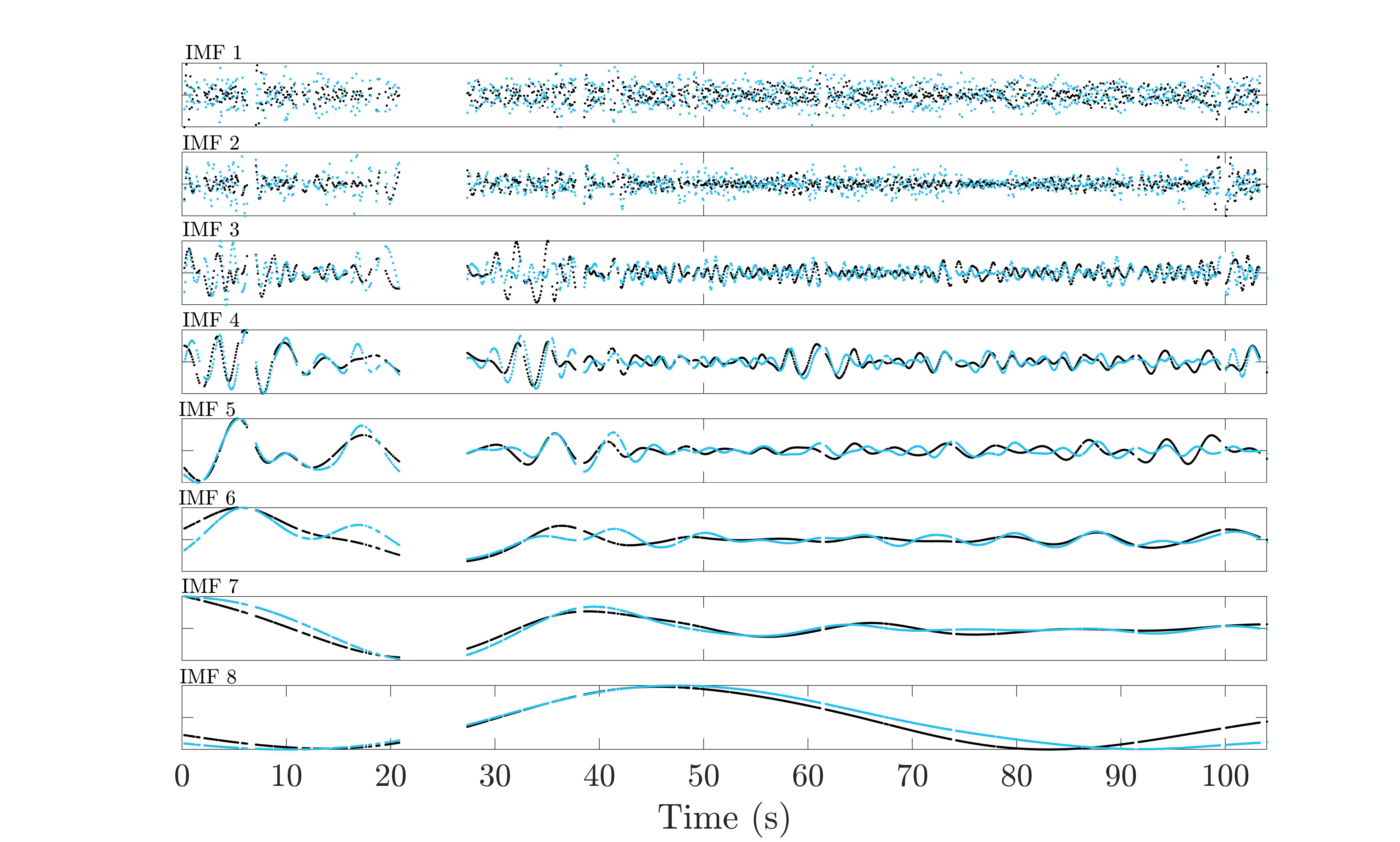

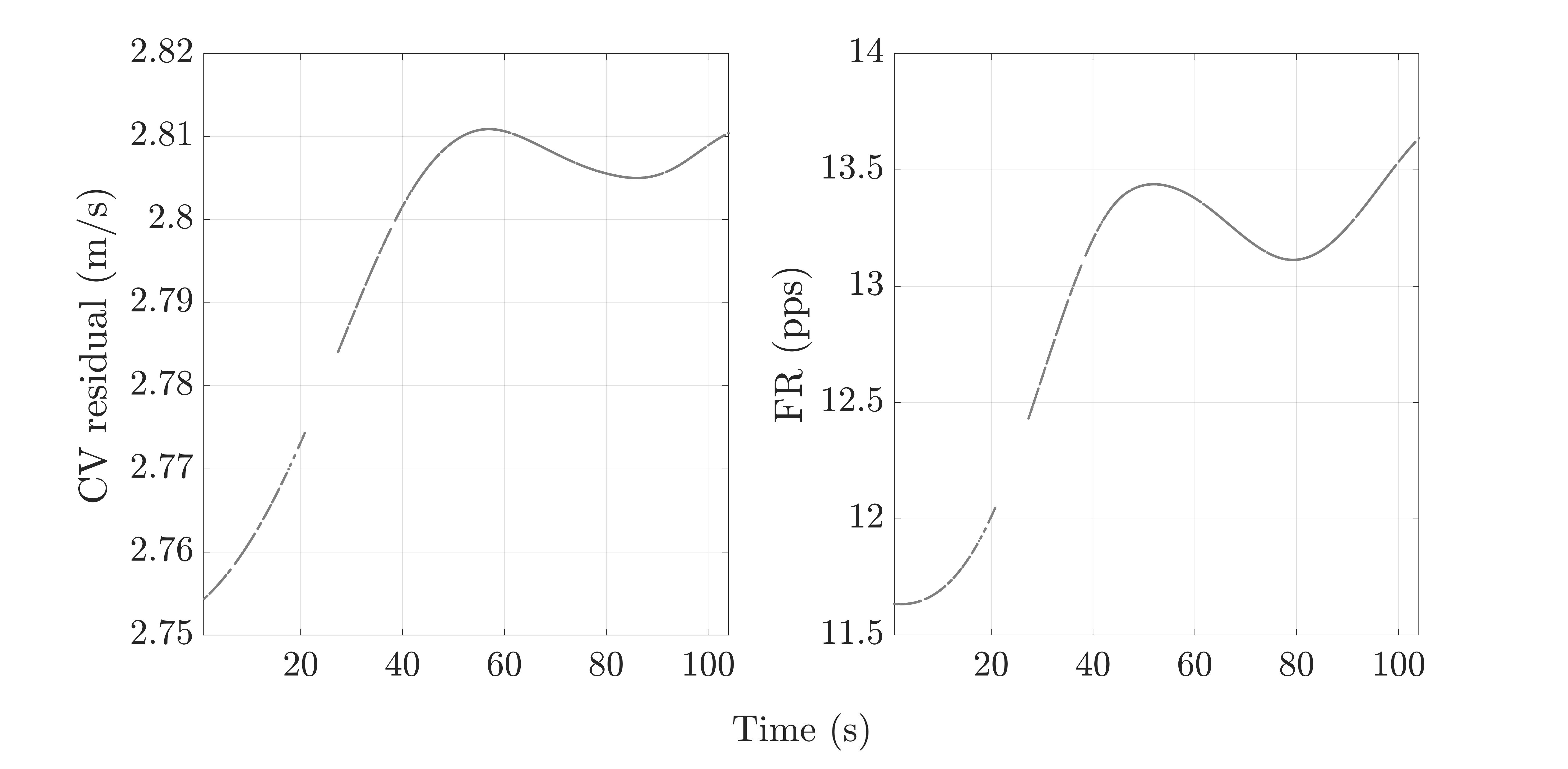

Significance of IMFs components:

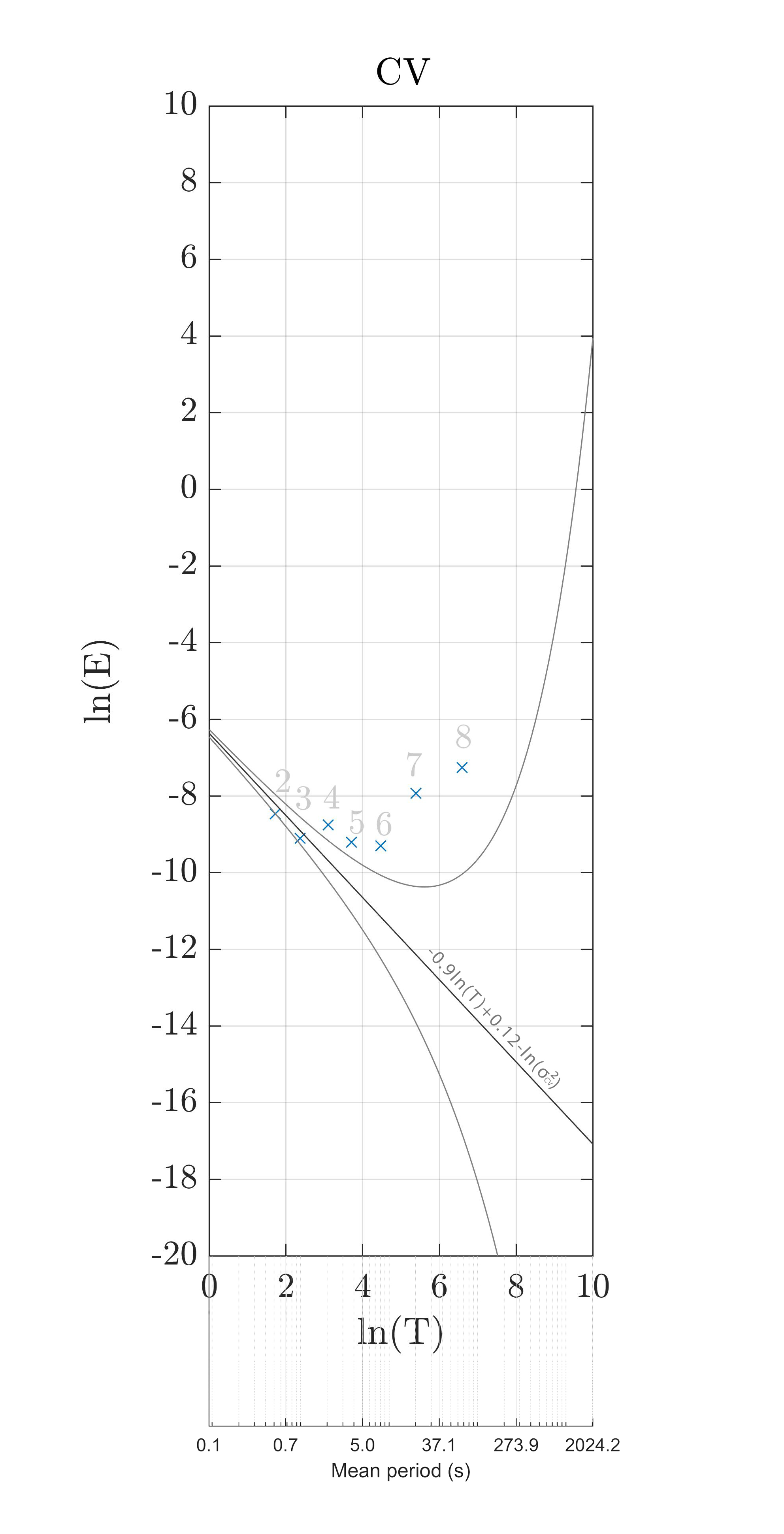

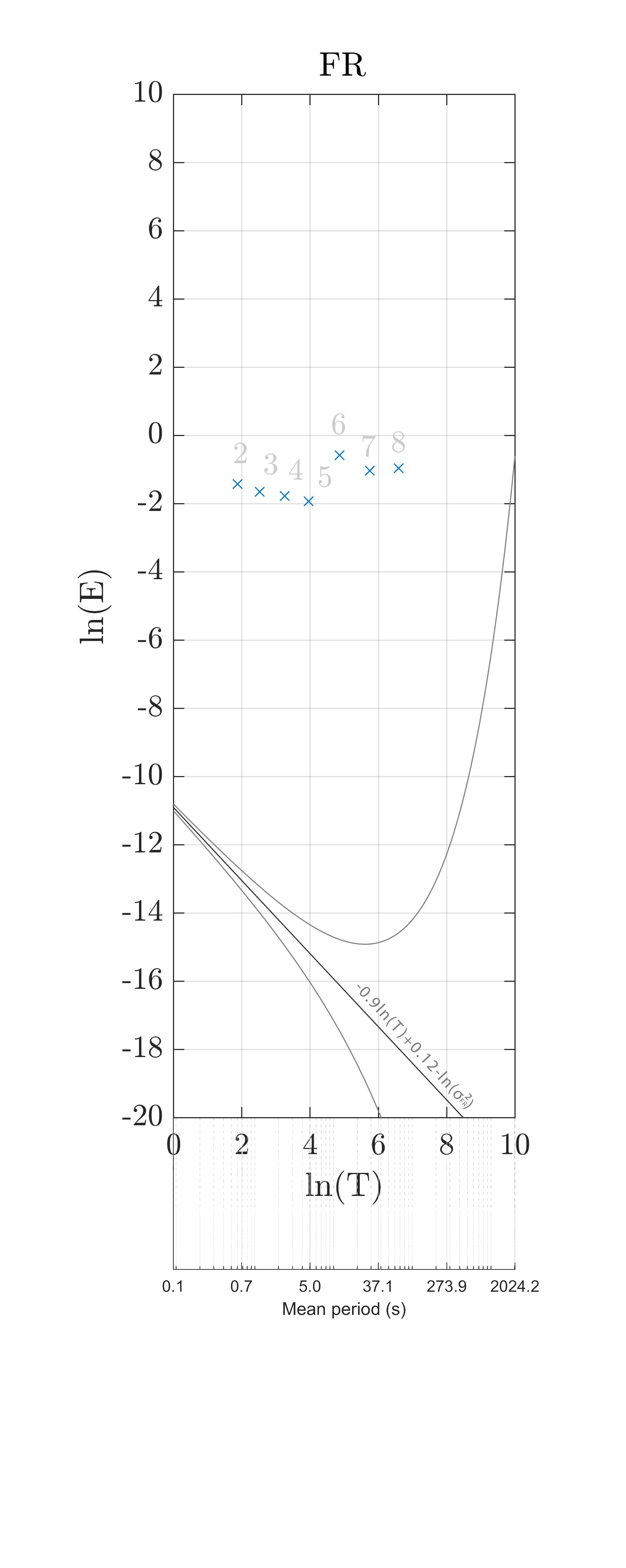

Raw CV time series and CV time series with removed high-frequency components:

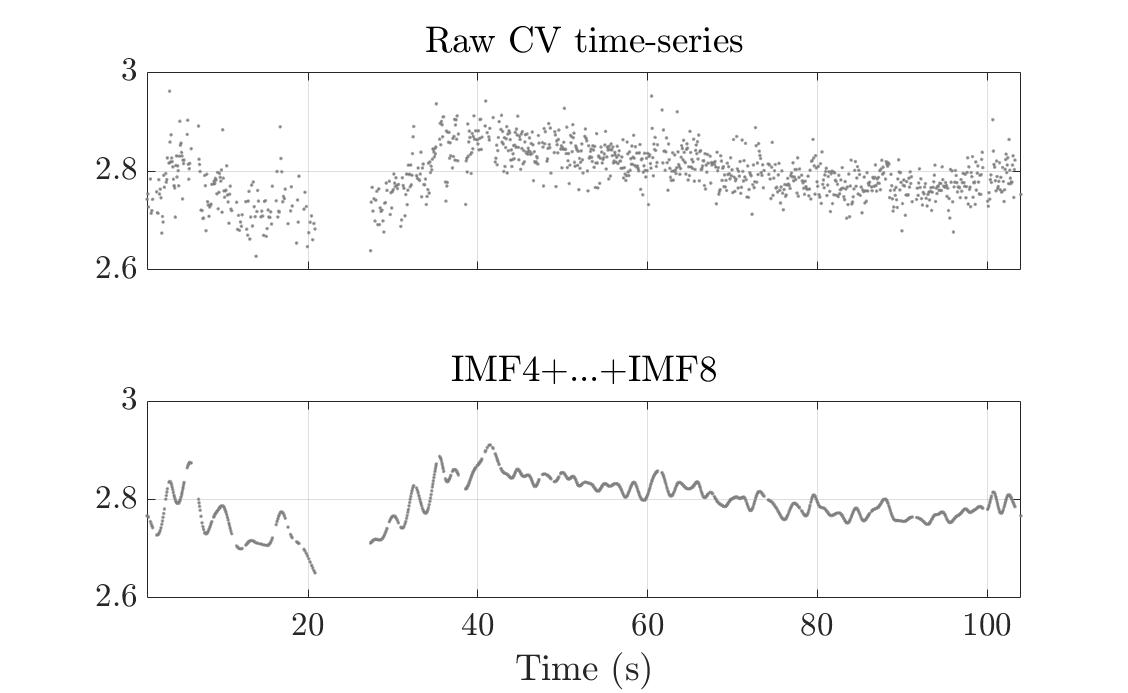

Time-dependent cross-correlation plots between raw MU CV and MU FR time-series:

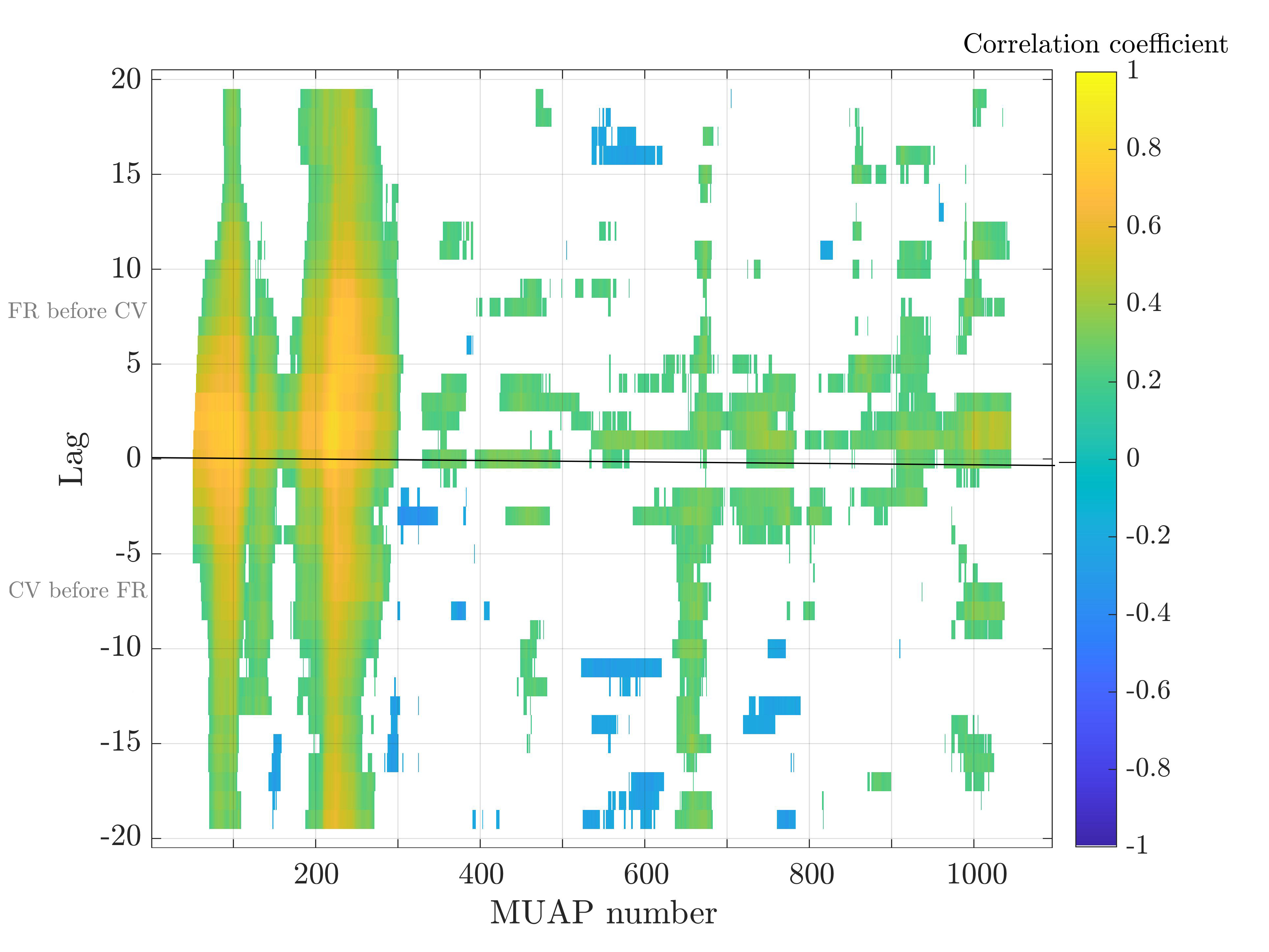

Time-dependent cross-correlation plots between mid-frequency components of MU CV and MU FR time-series:

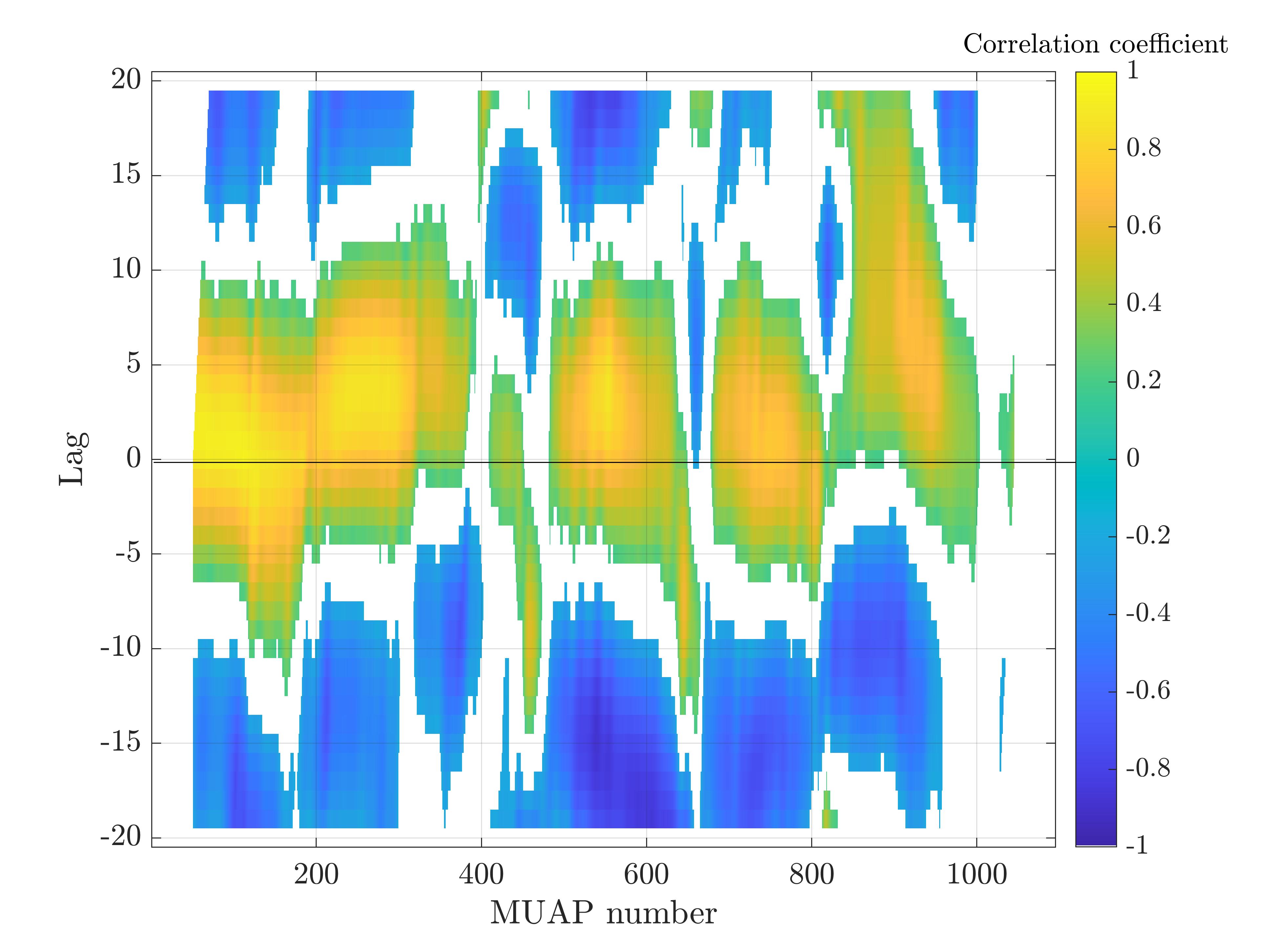

Subject 2

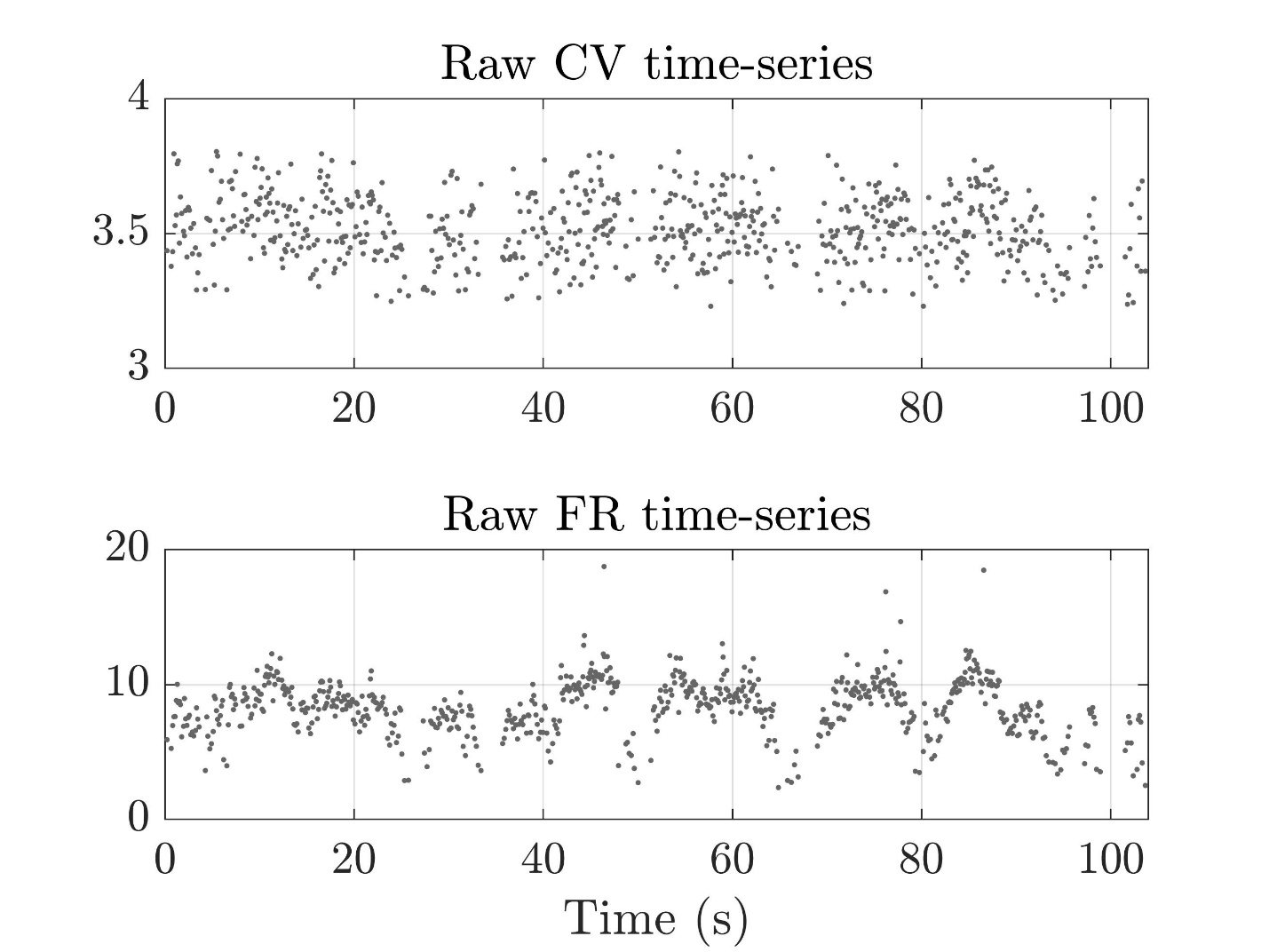

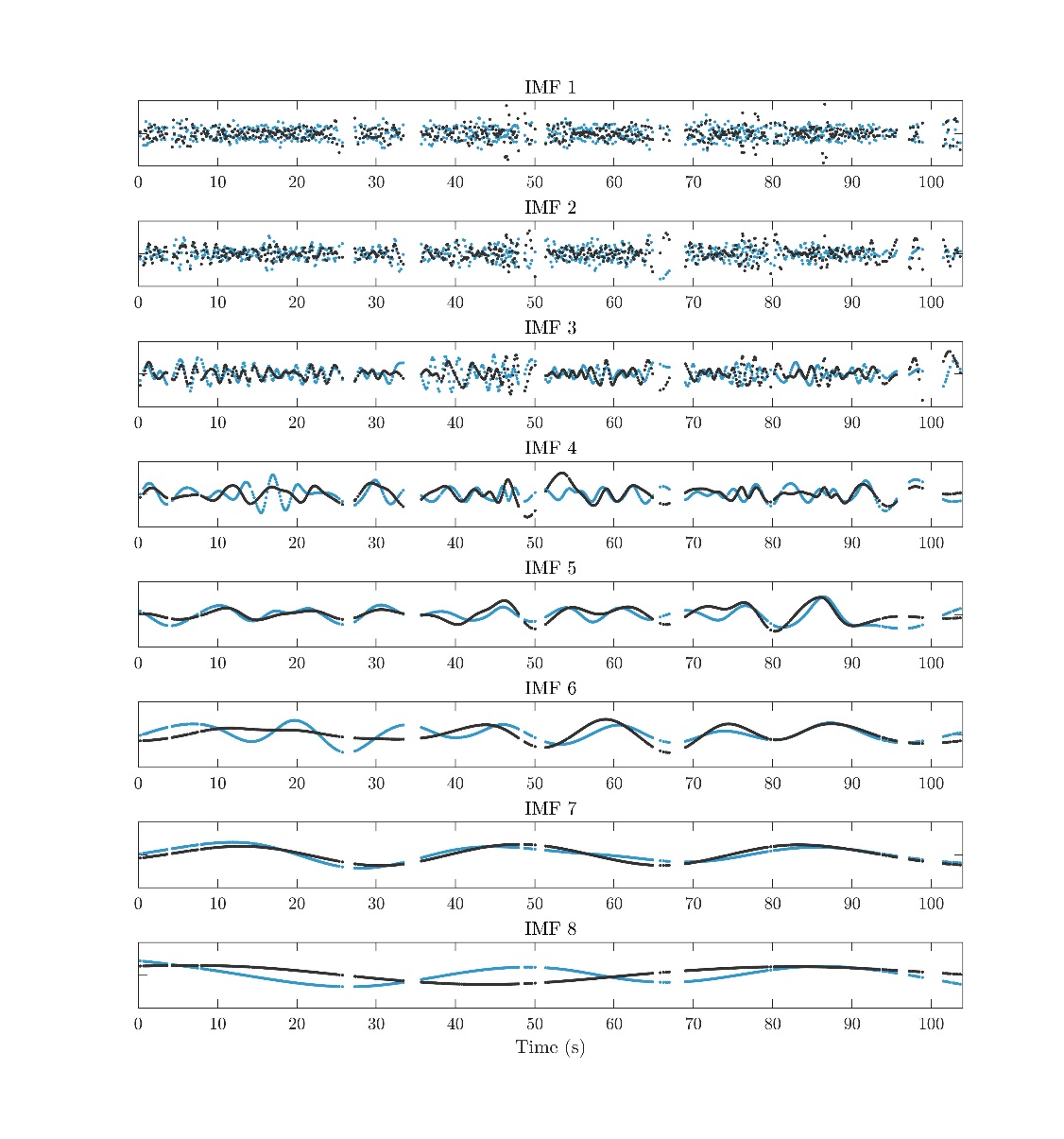
\

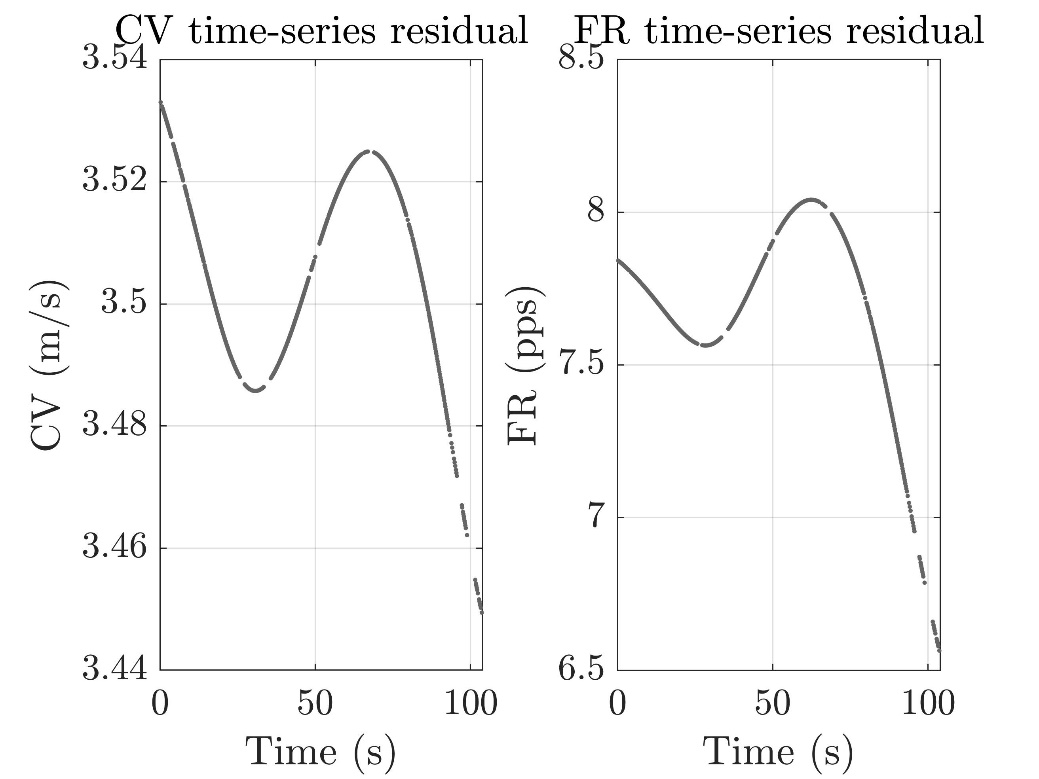

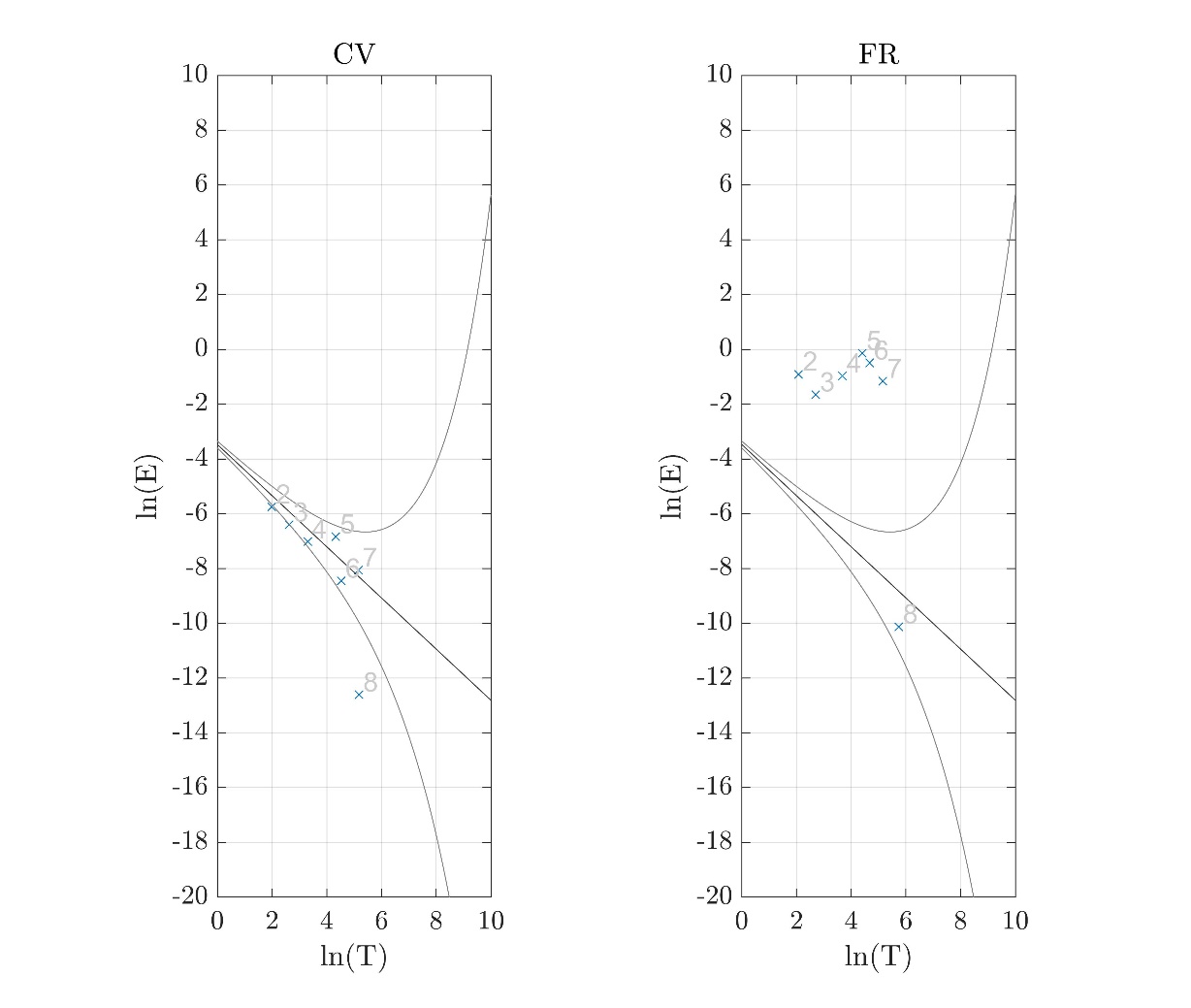

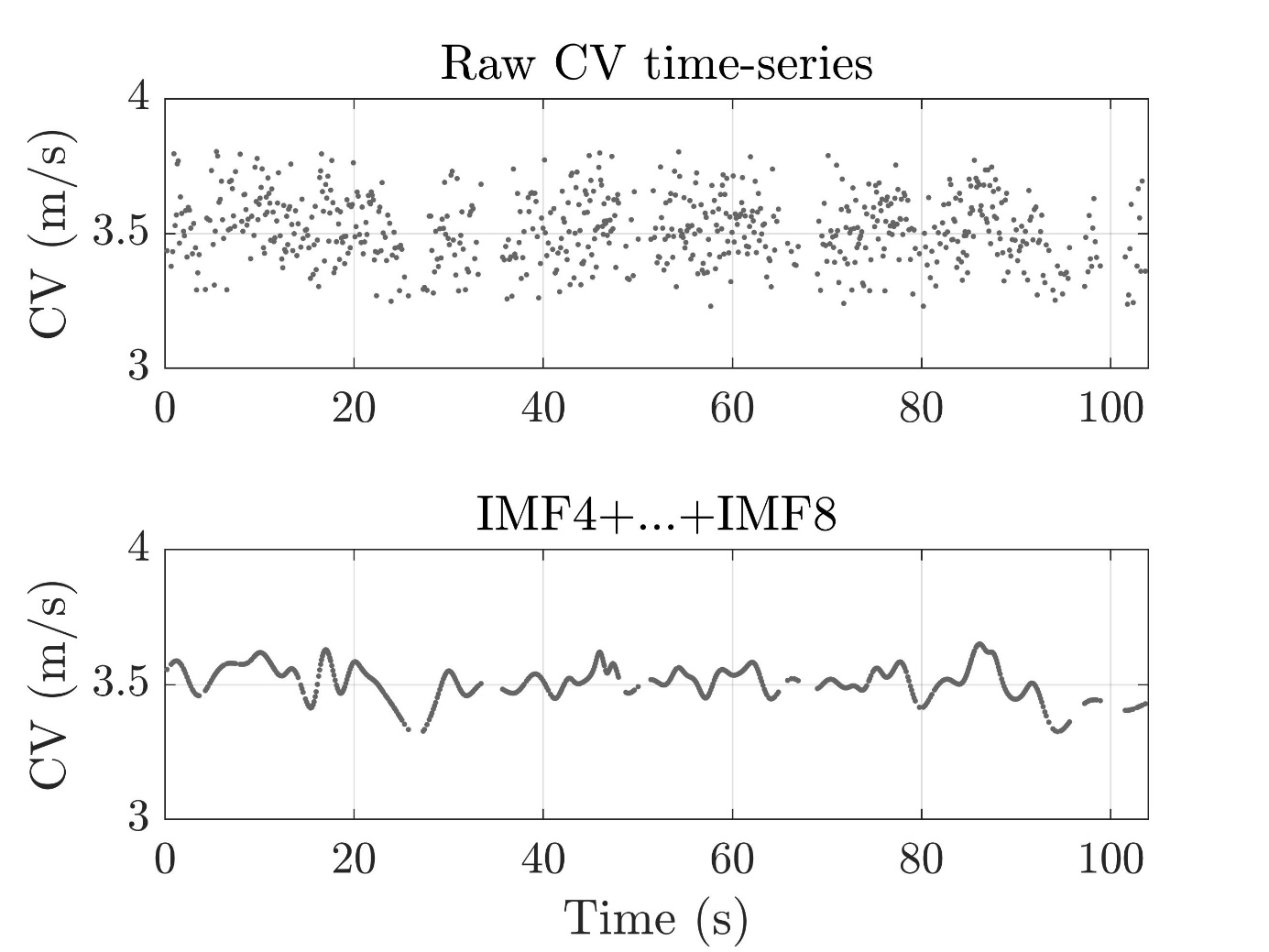

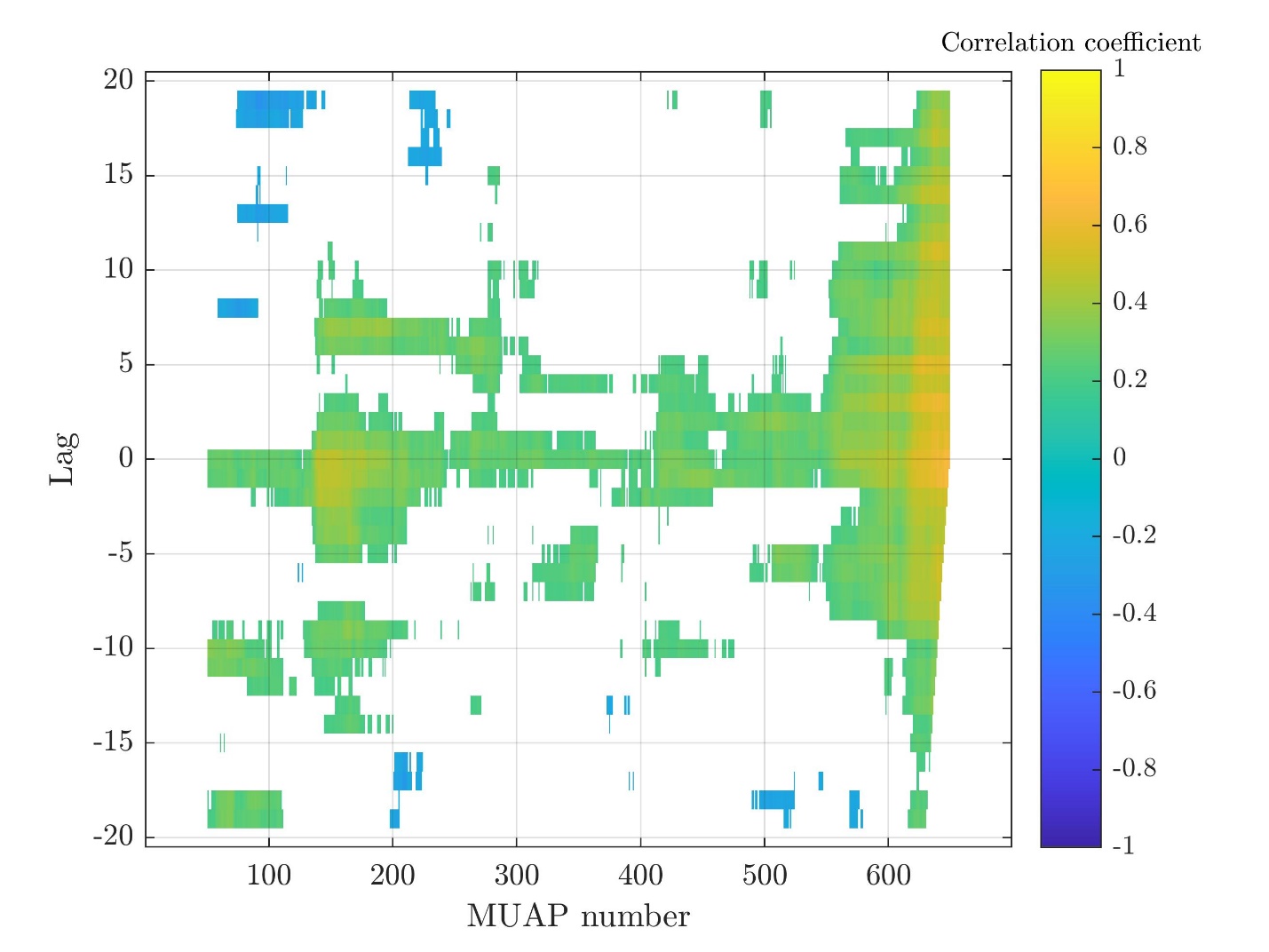

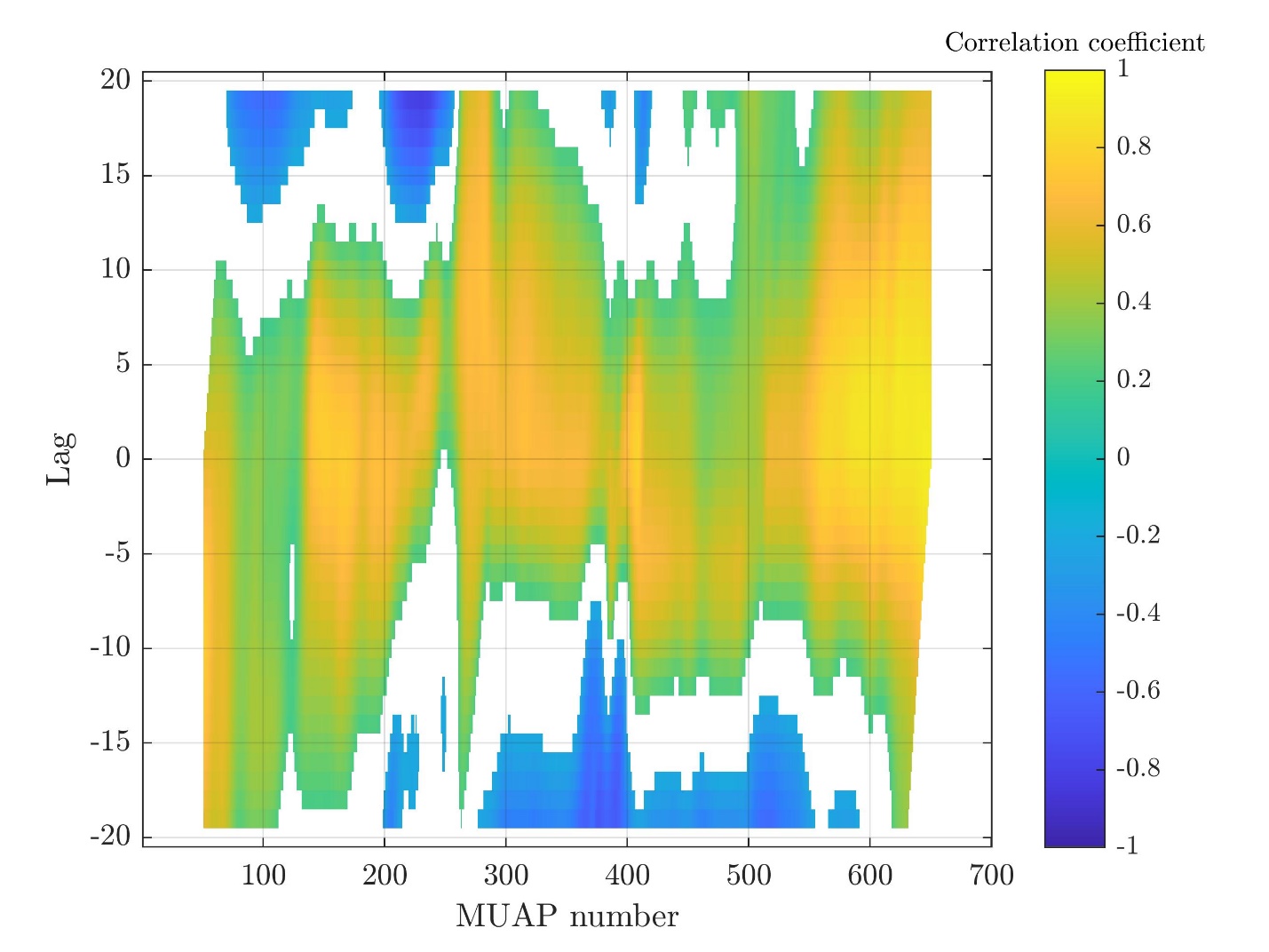

Subject 3

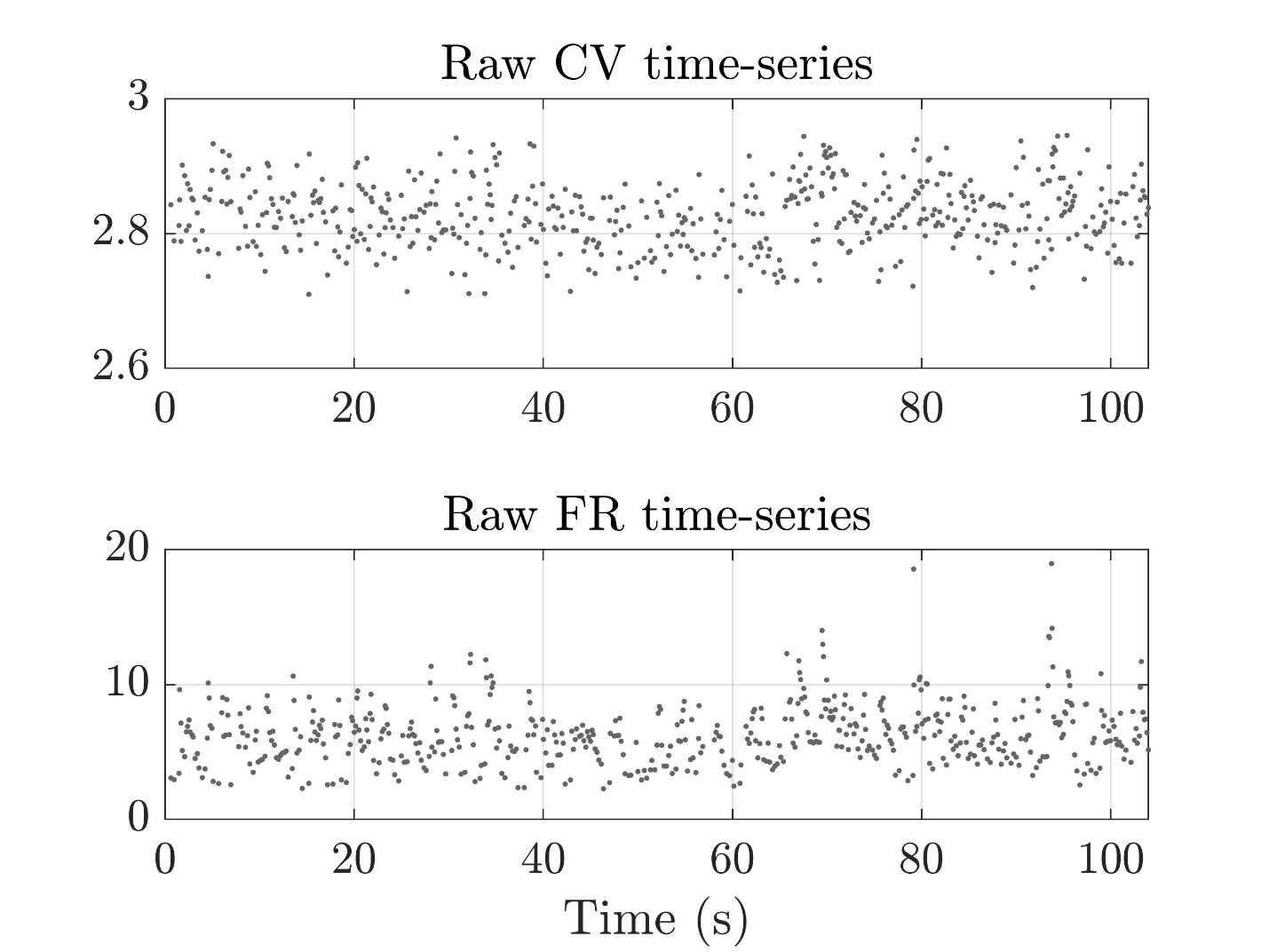

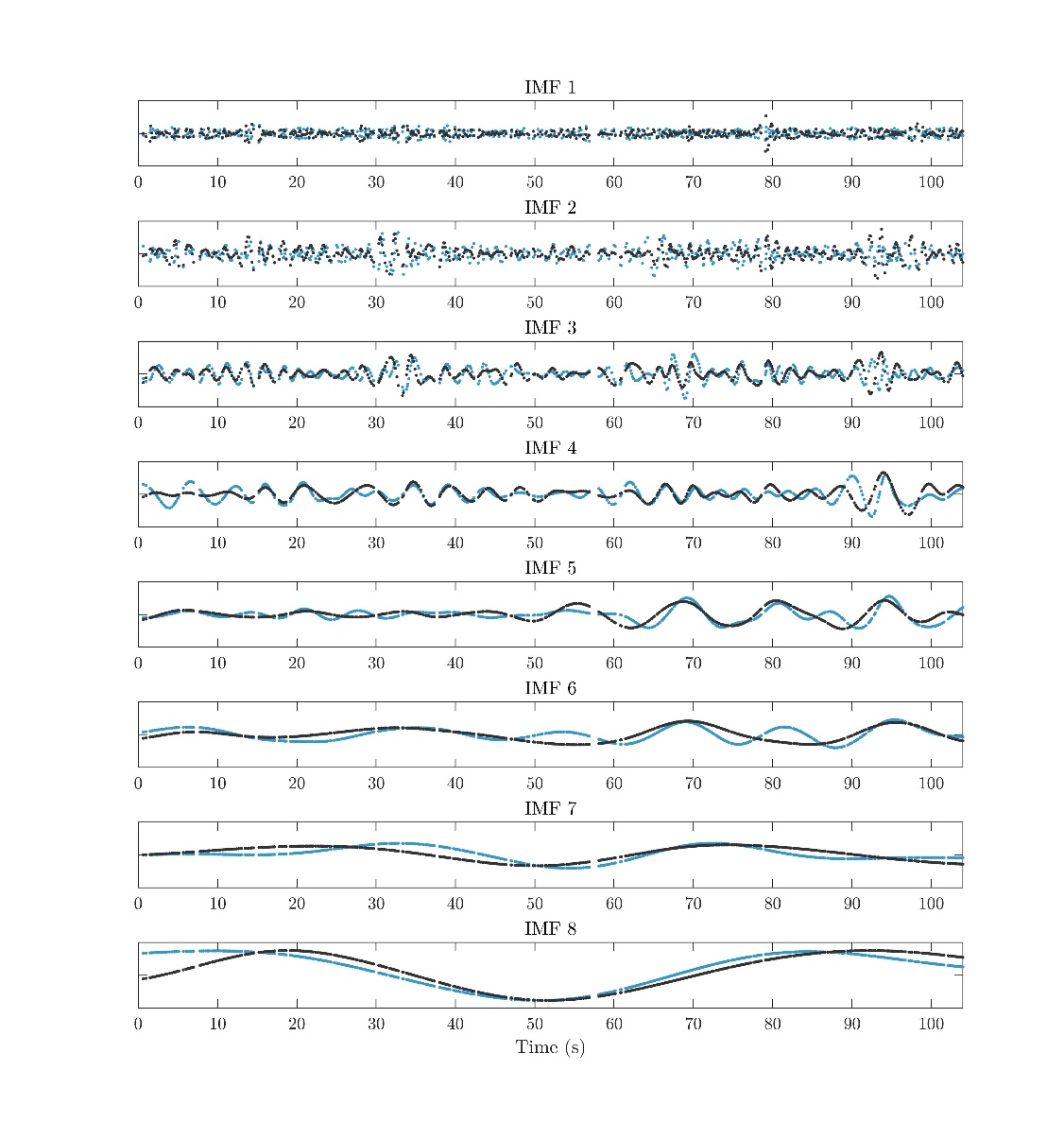

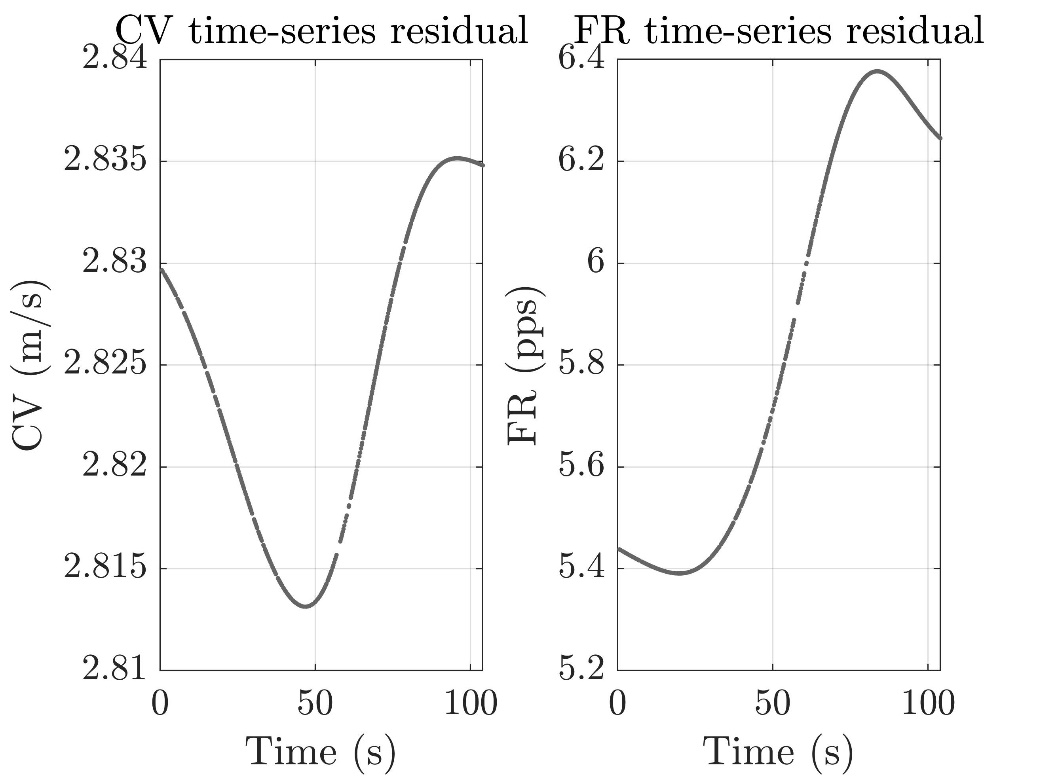

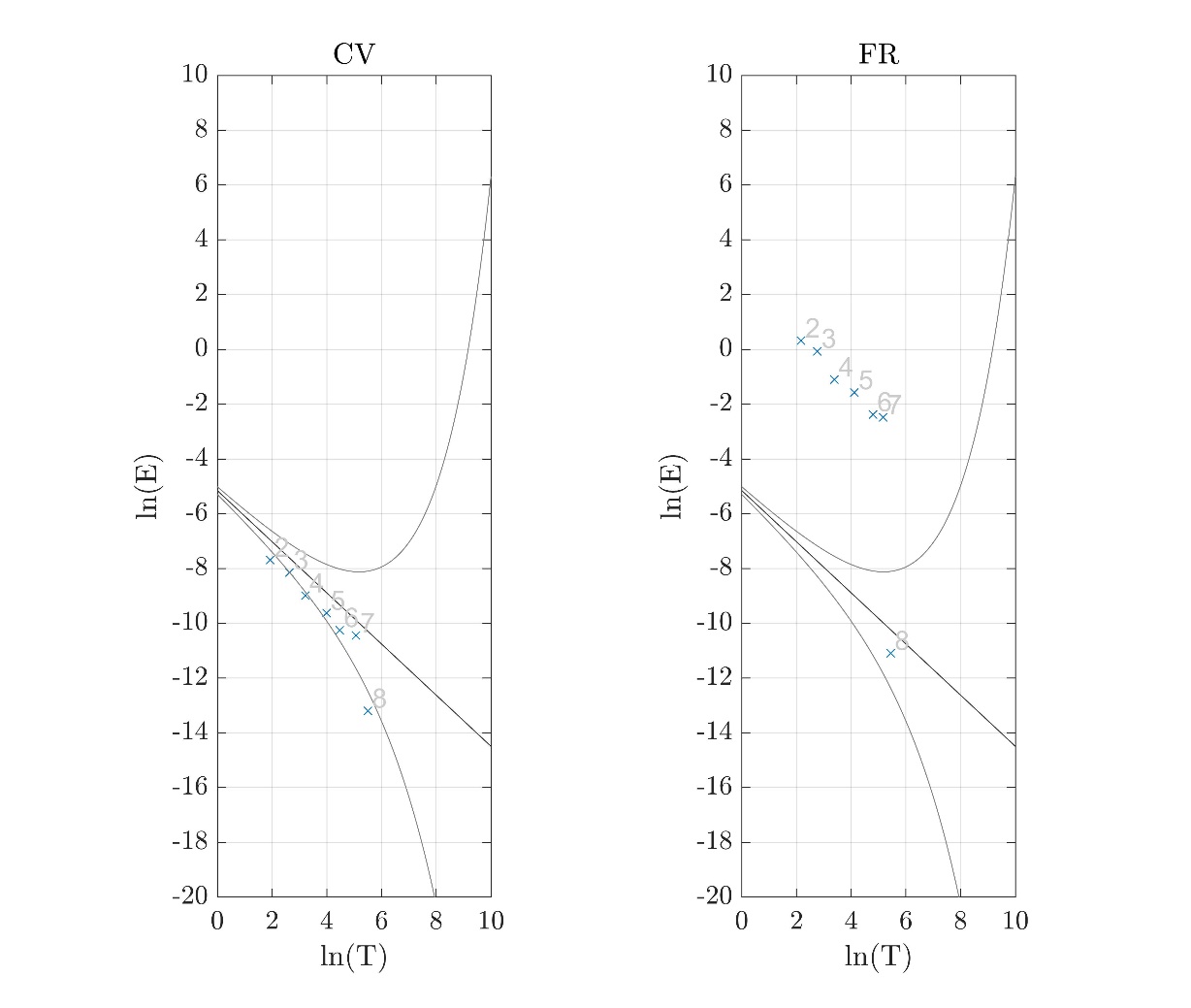

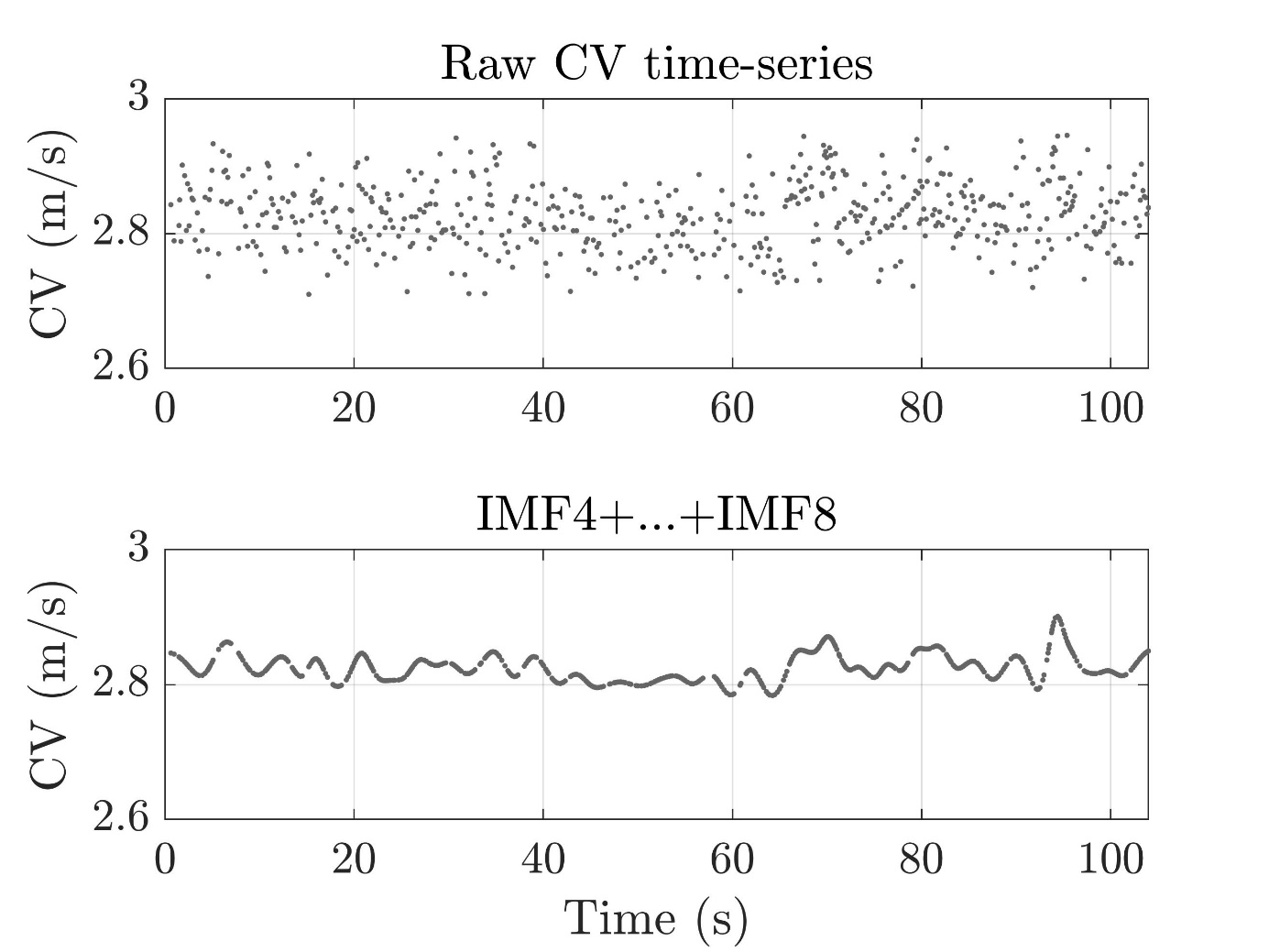

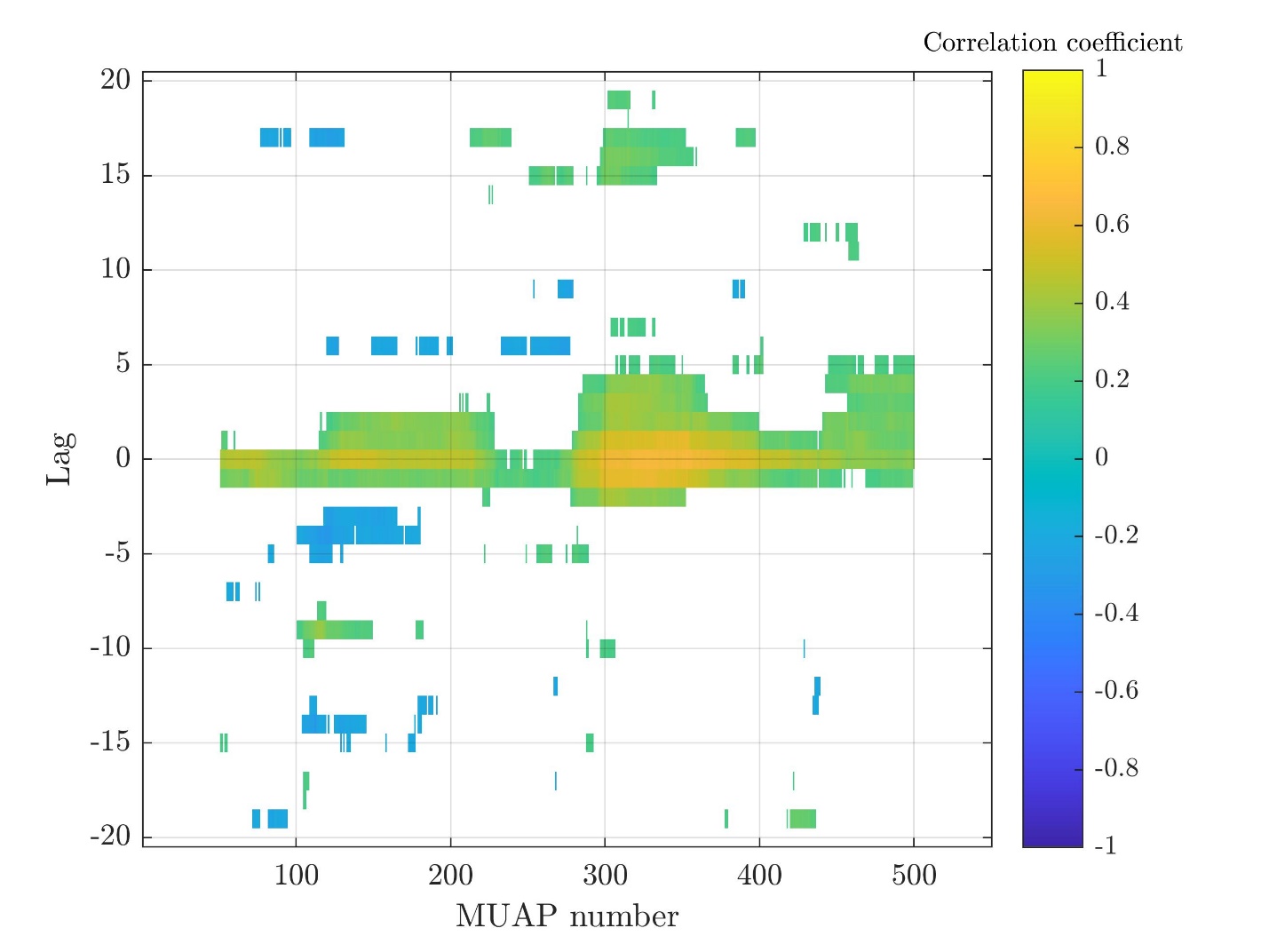

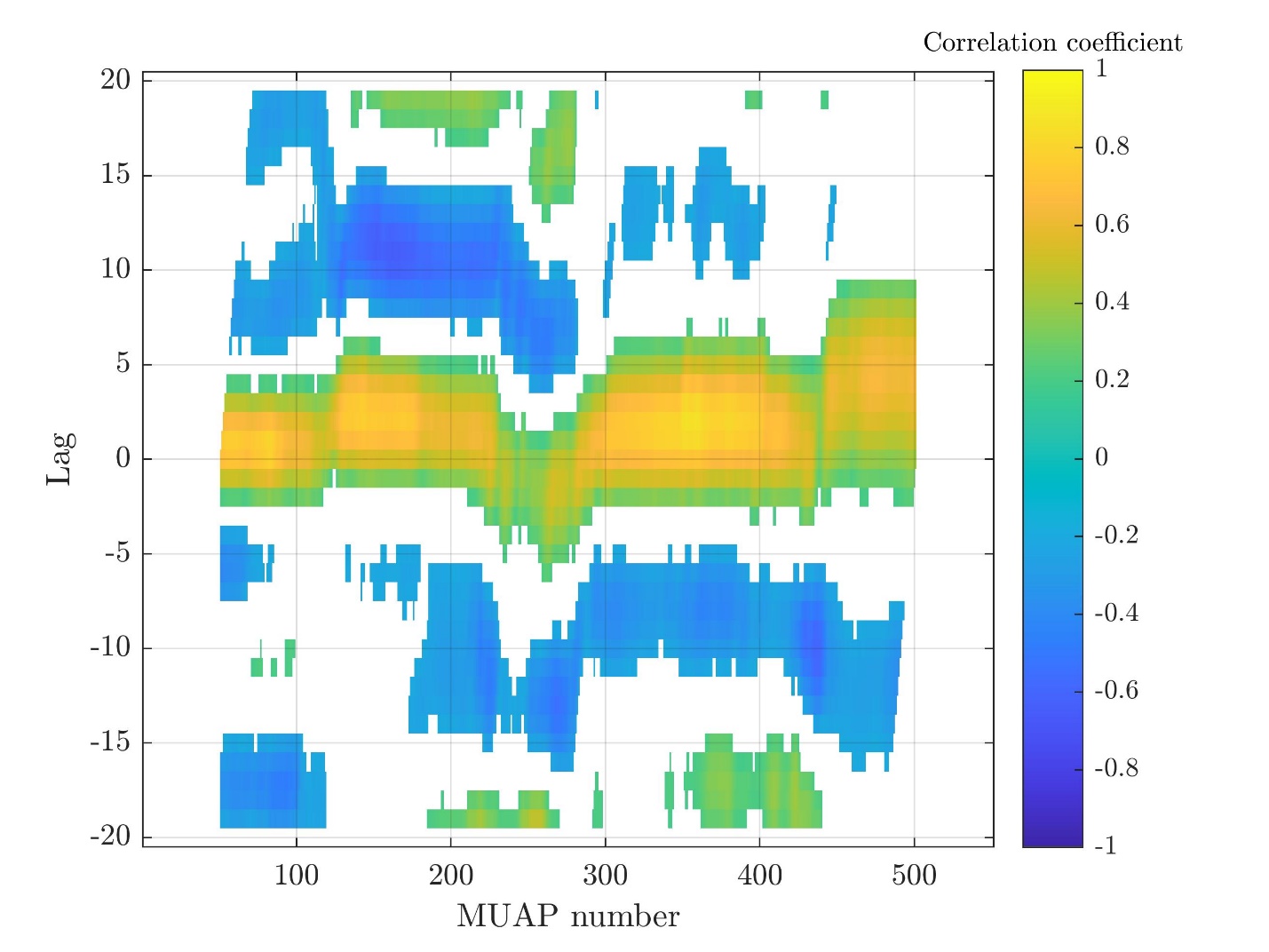

Subject 4

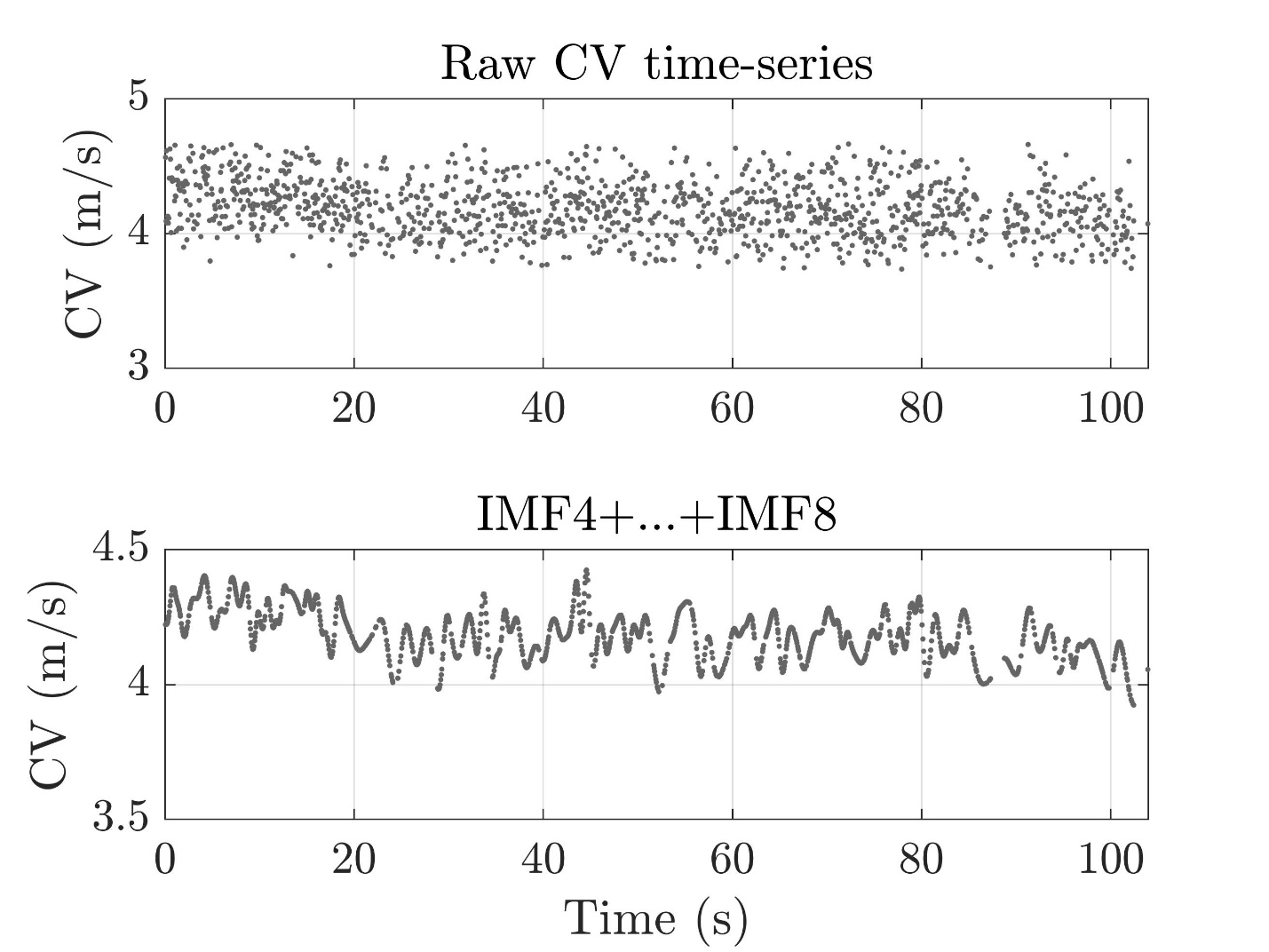

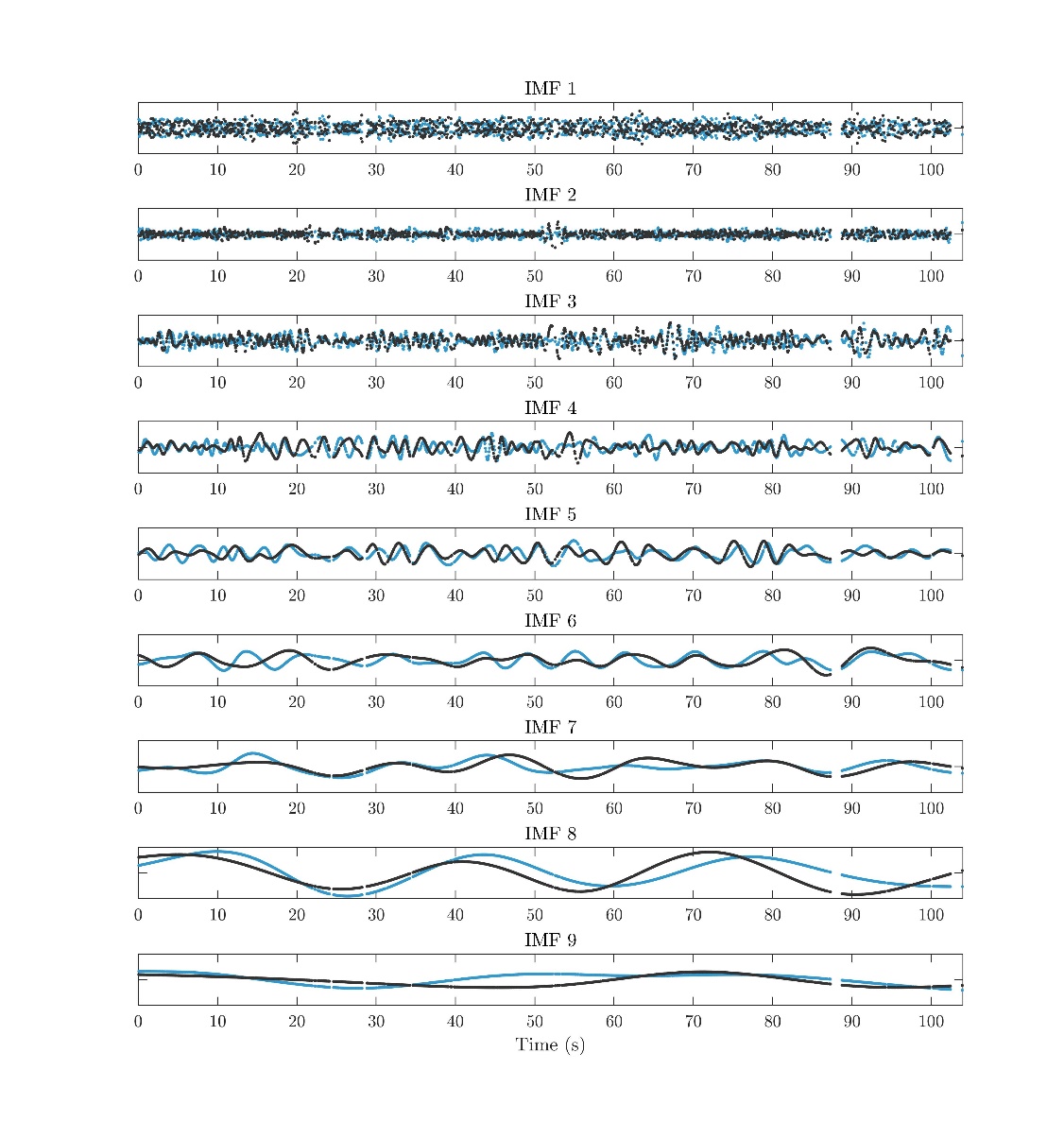

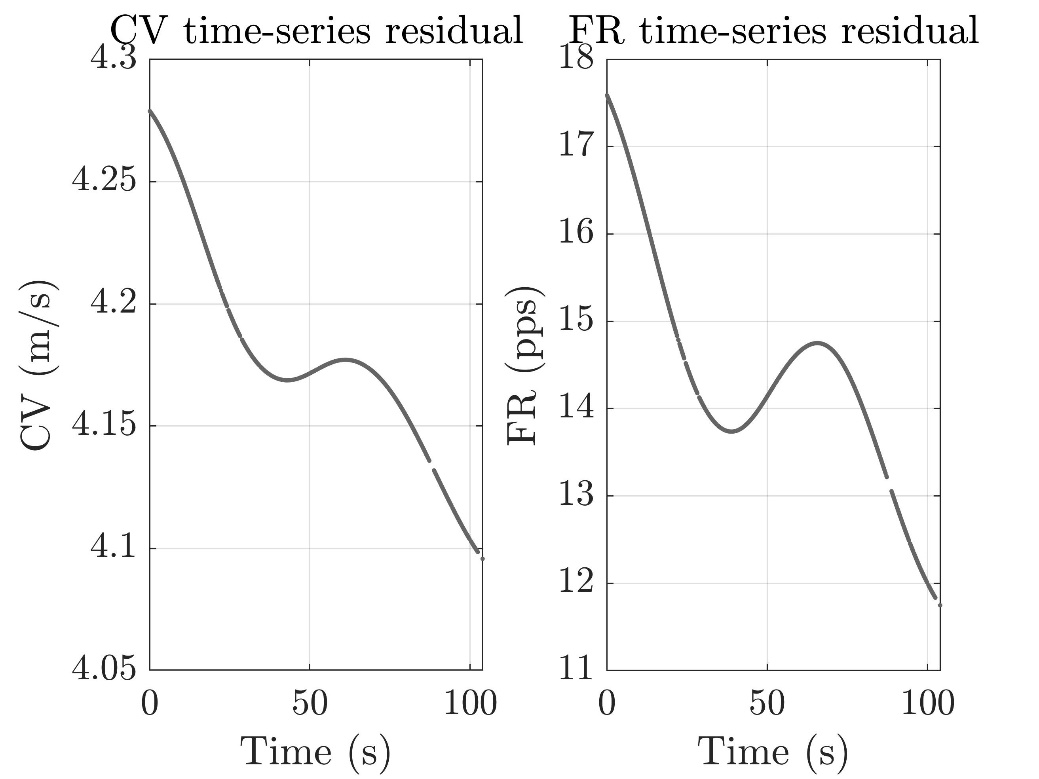

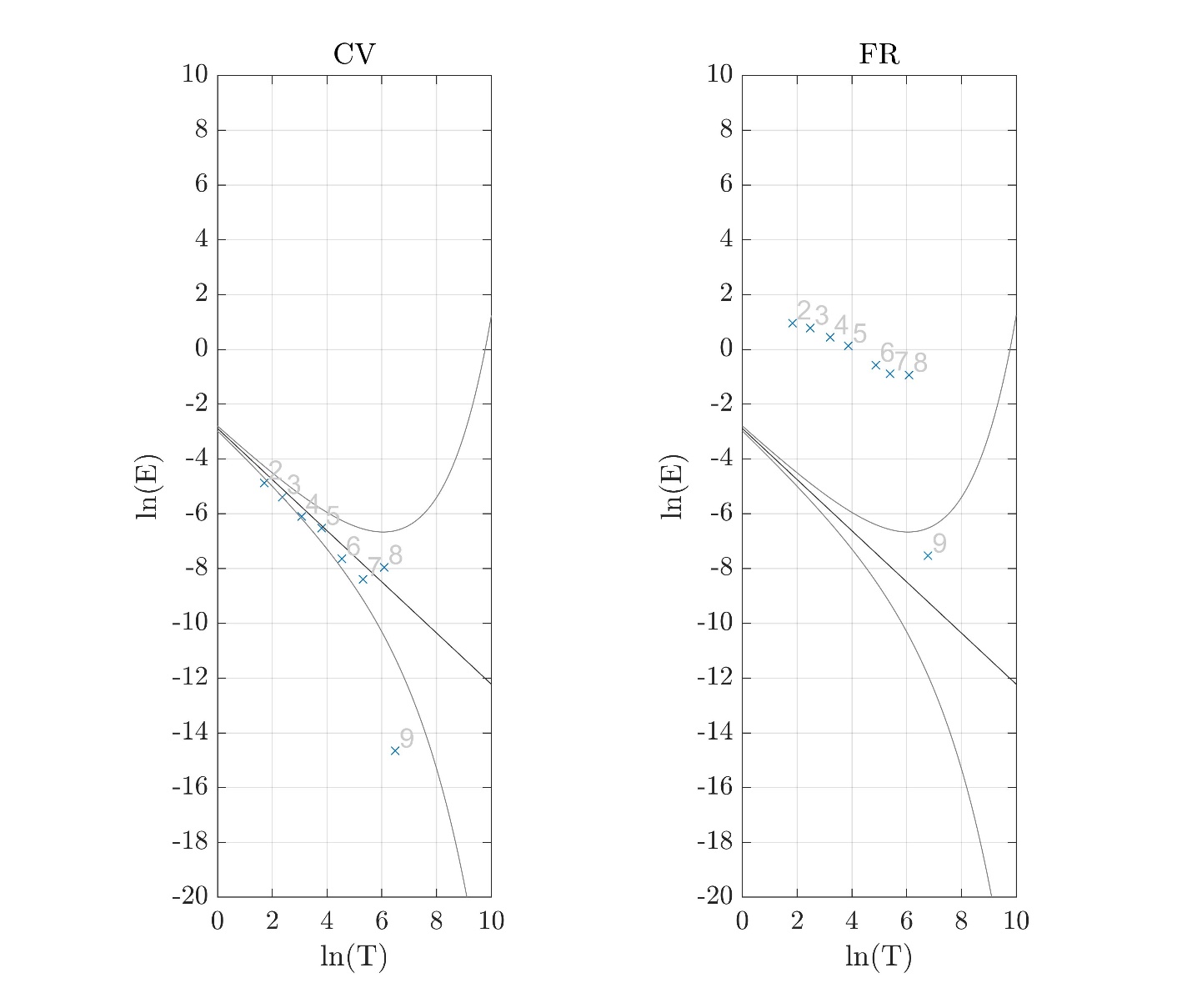

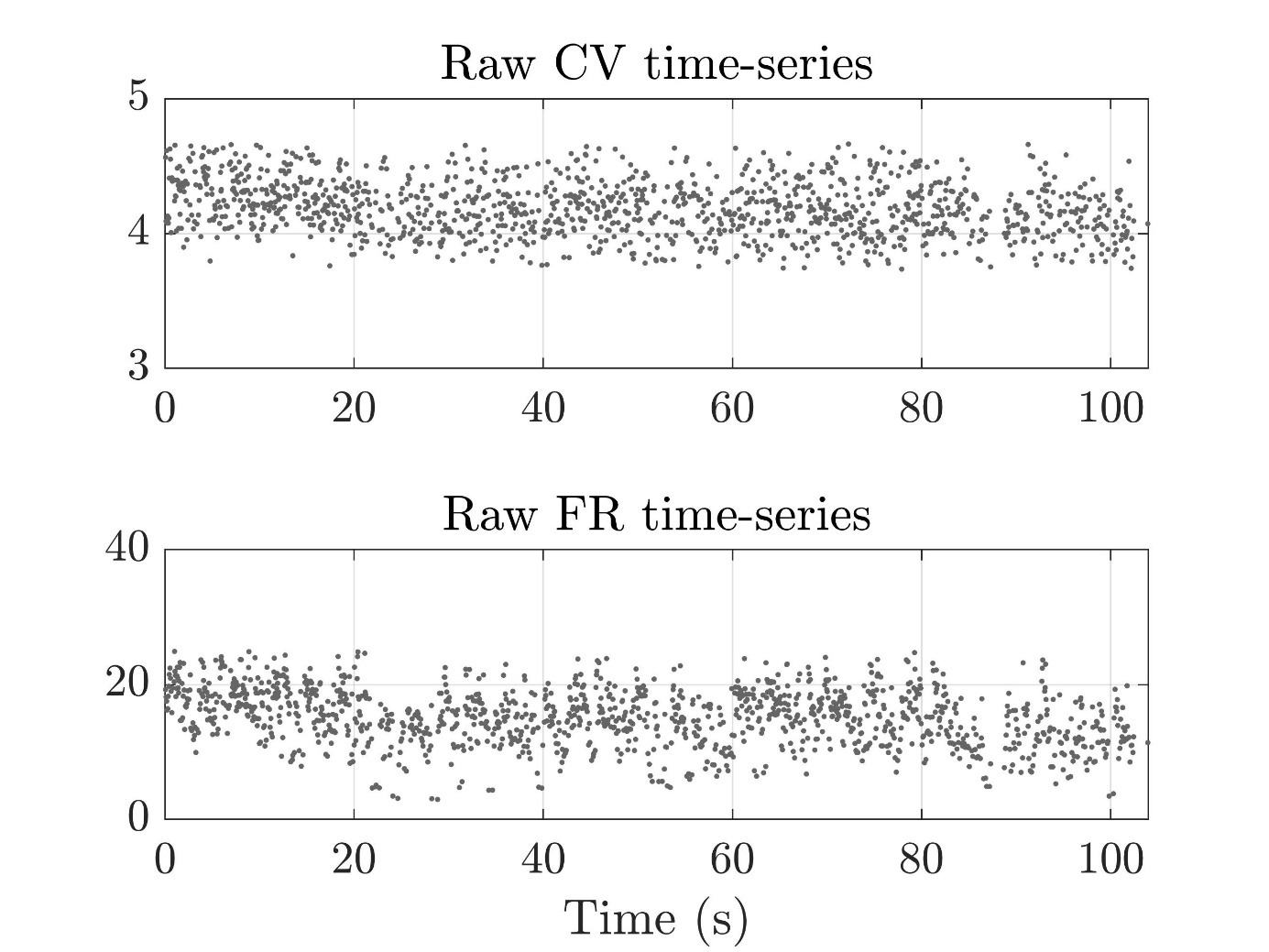

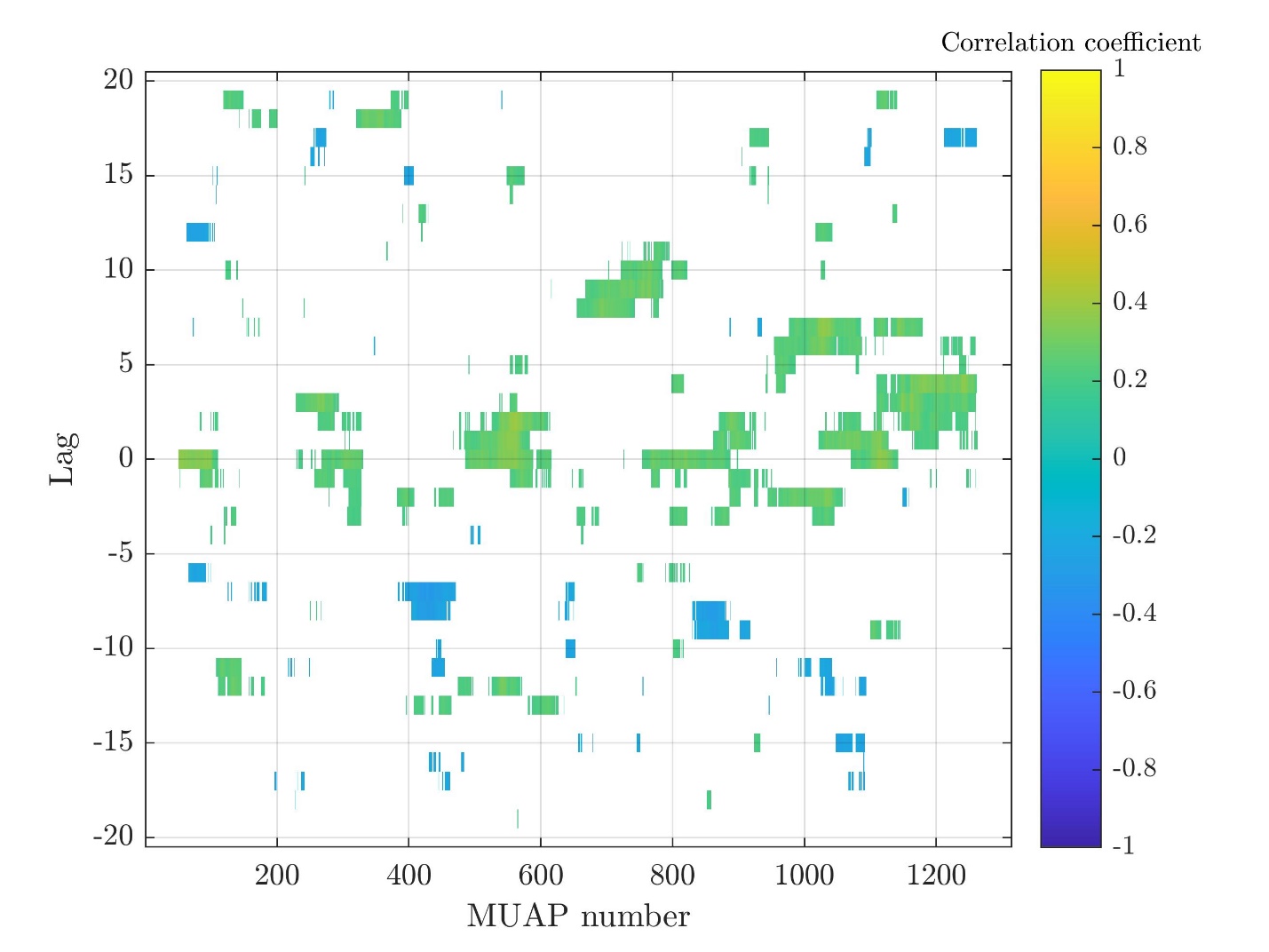

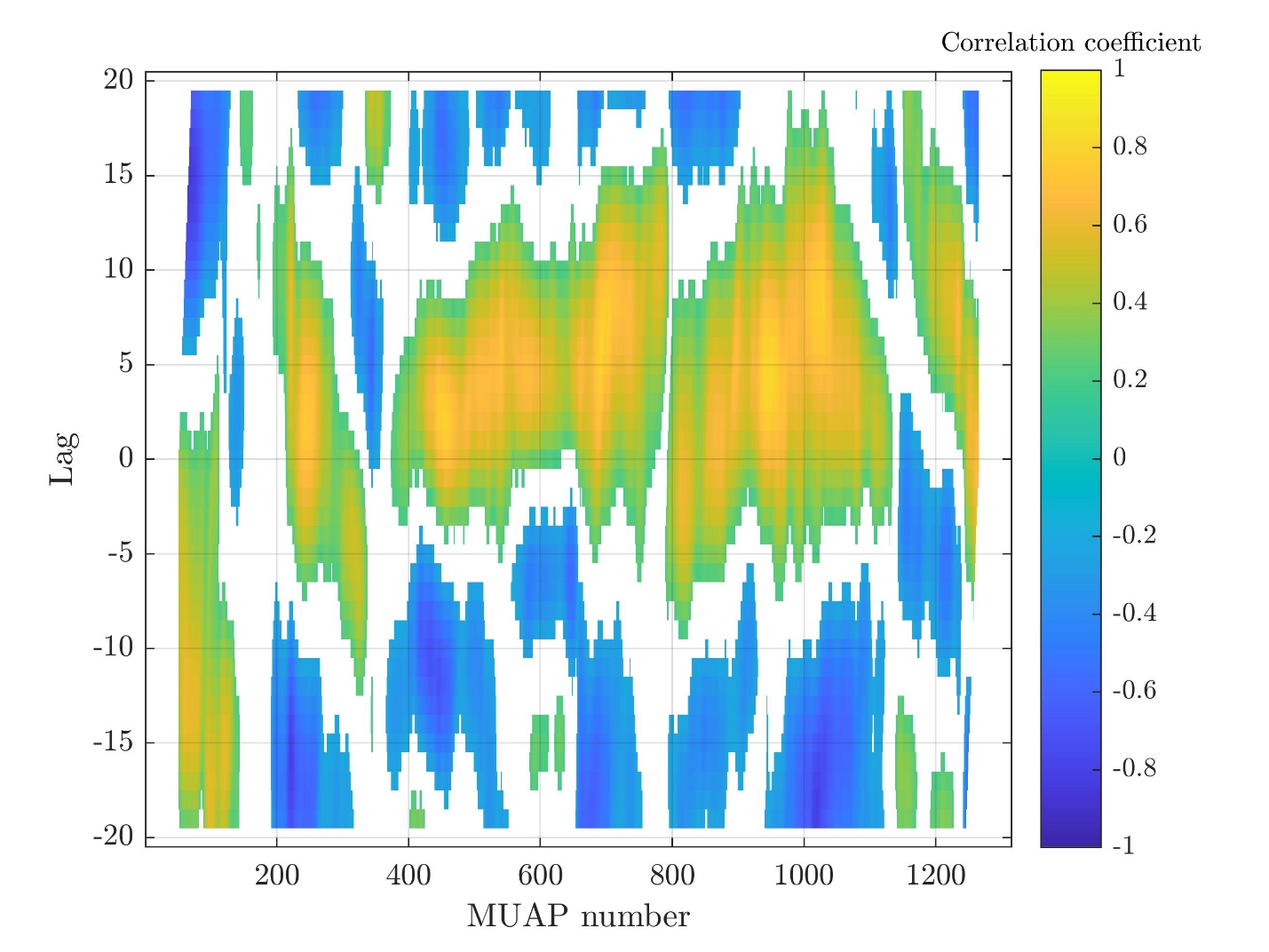

Subject 8

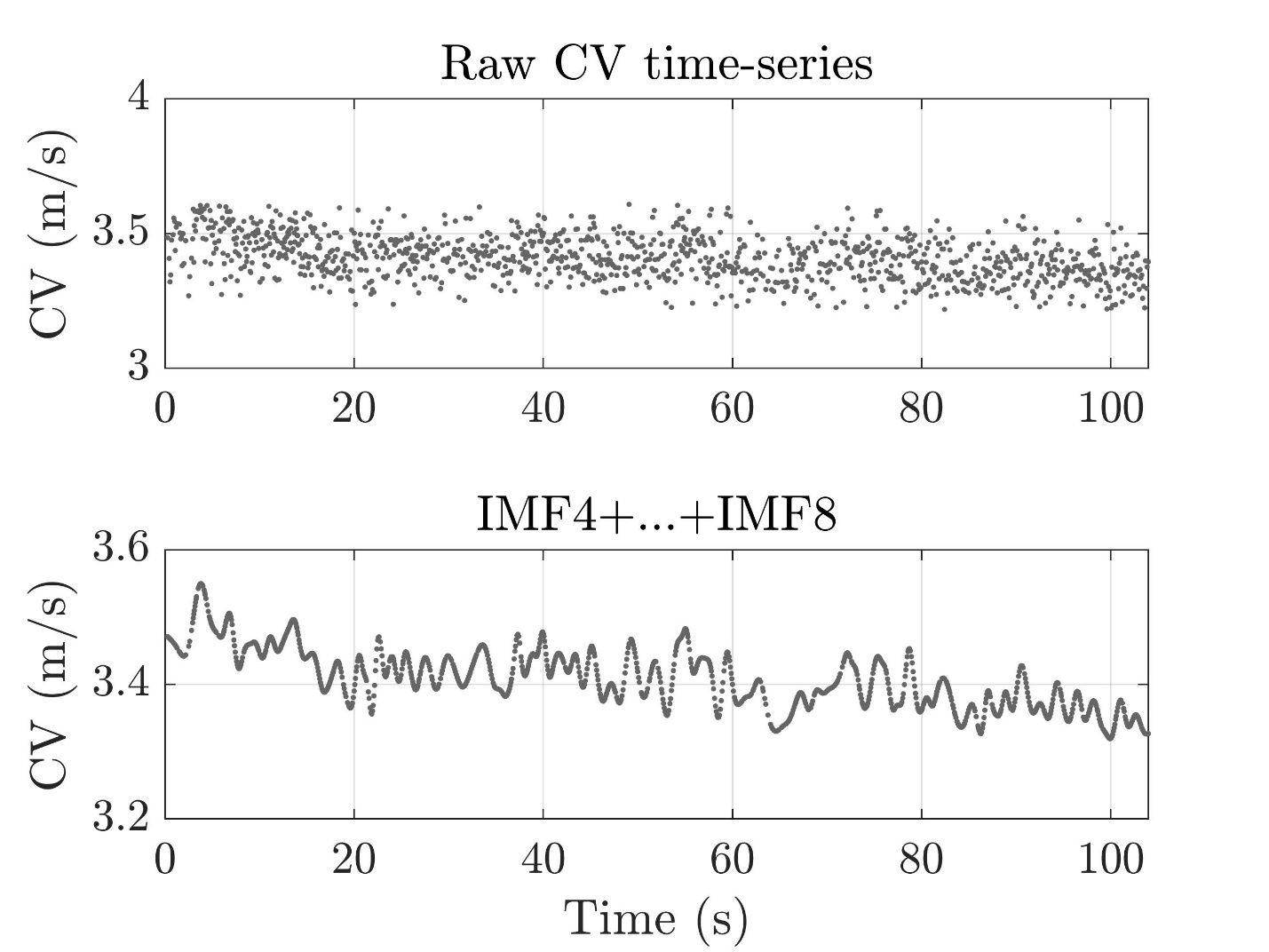

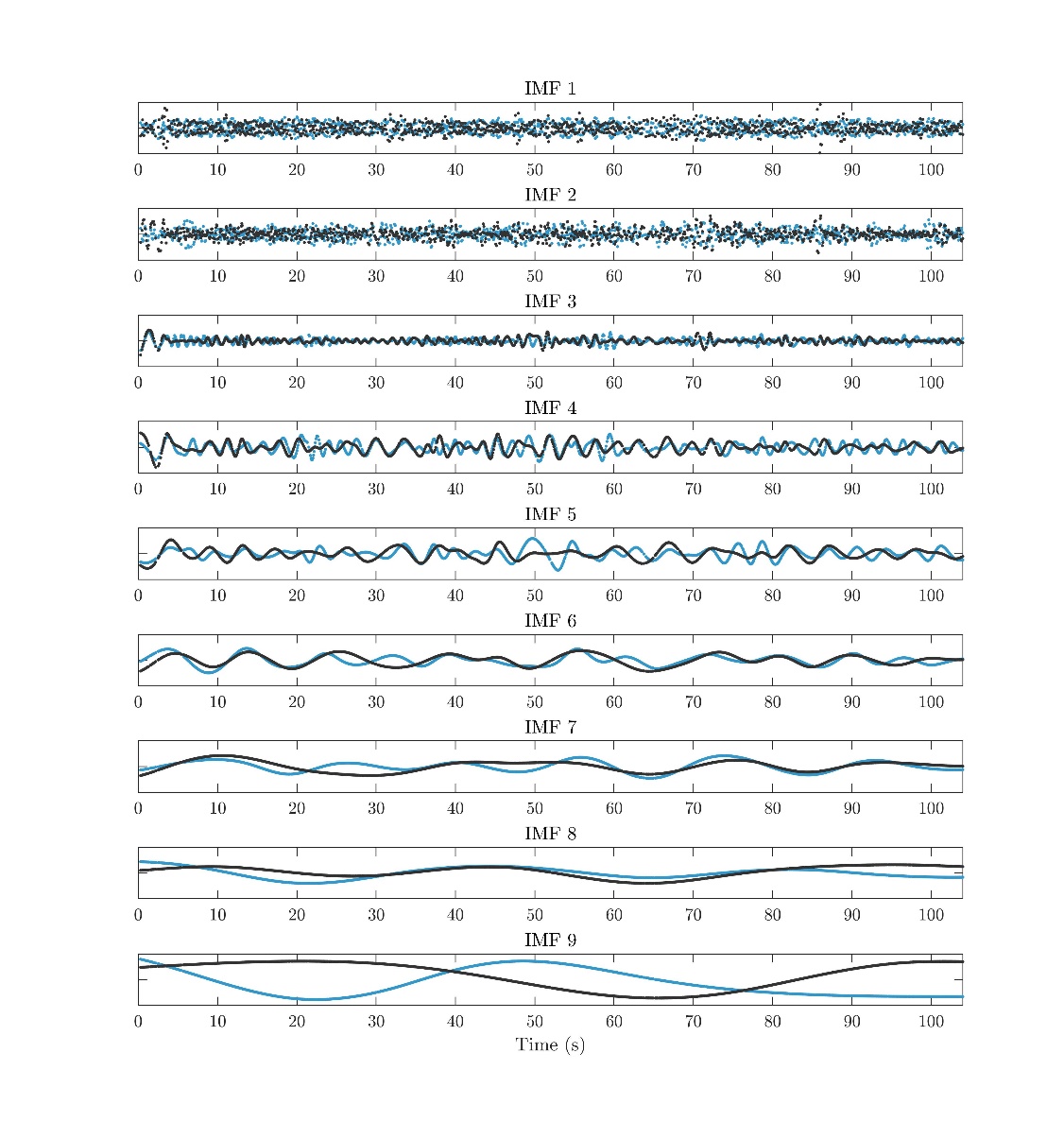

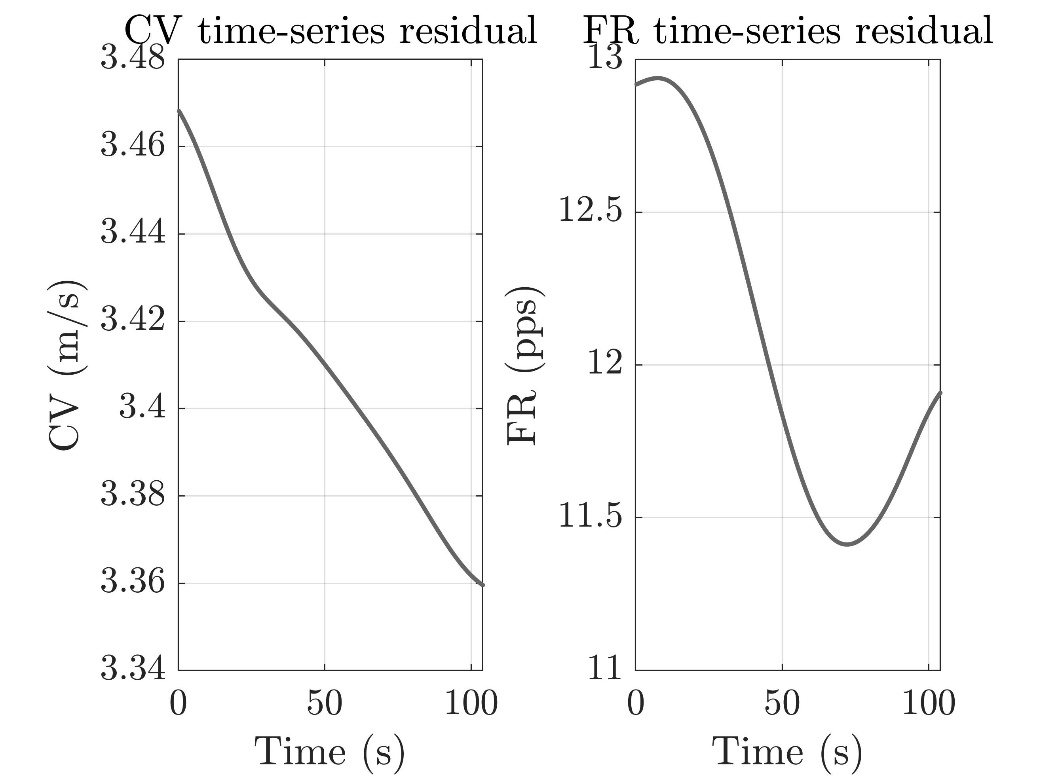

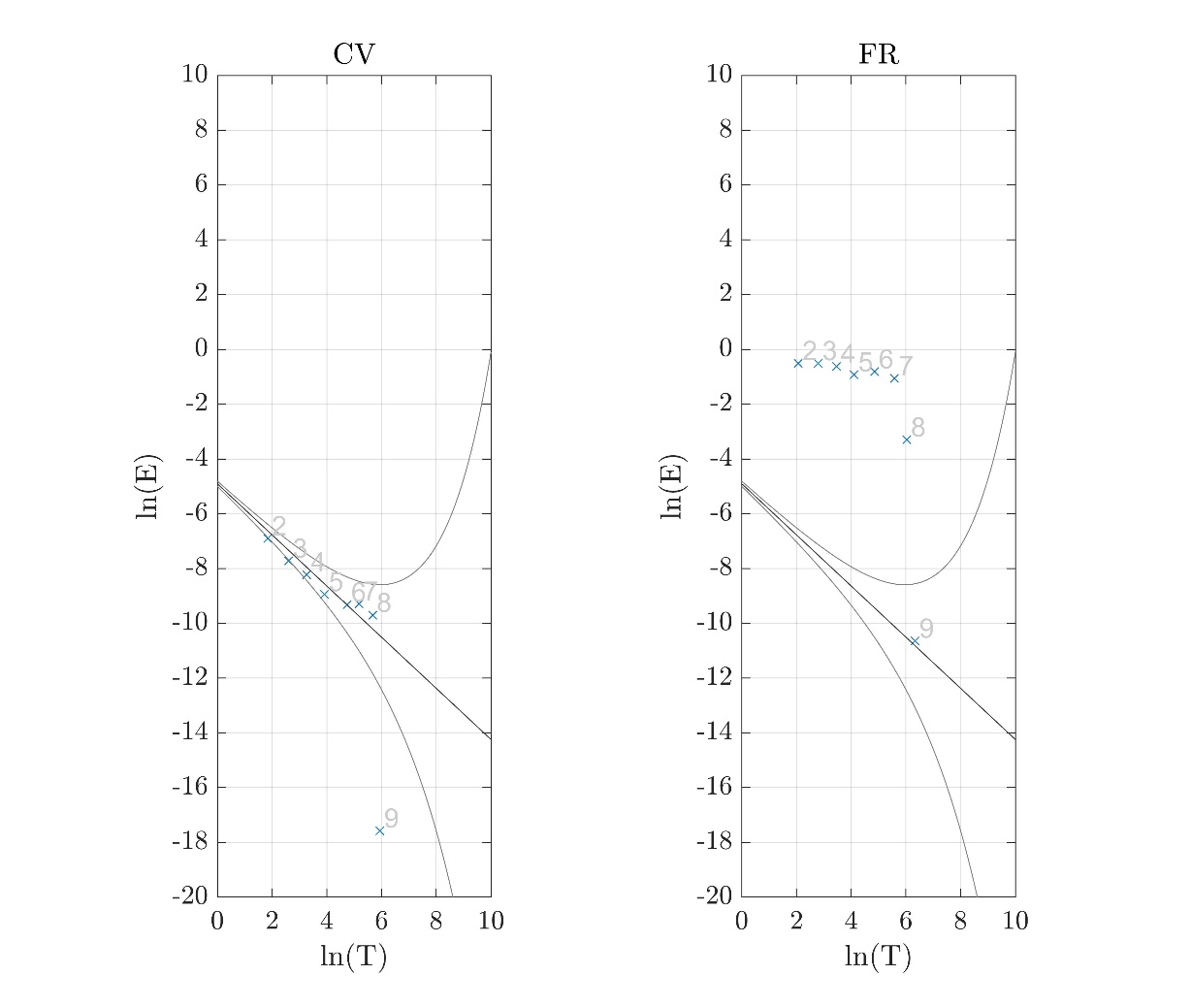

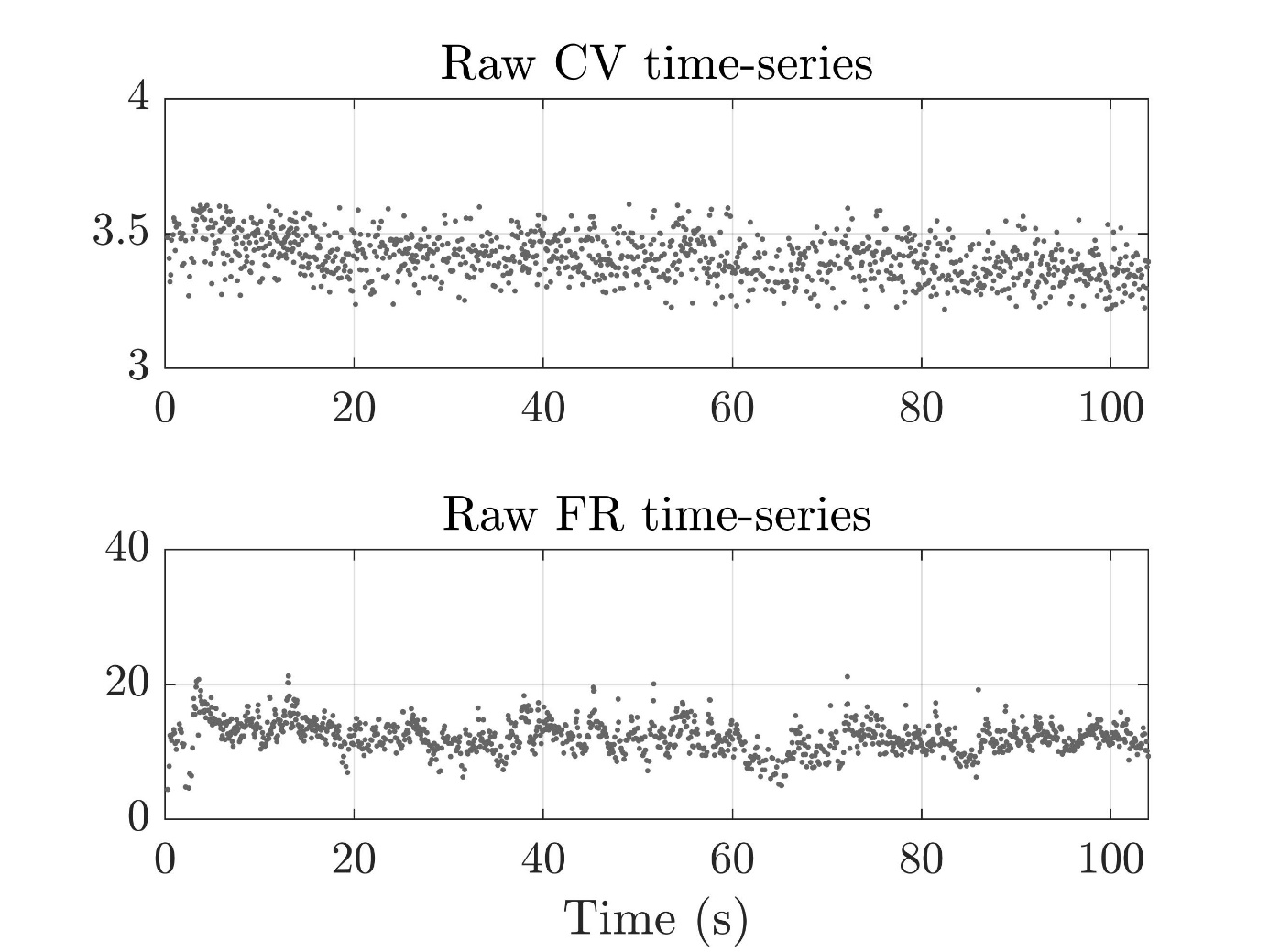

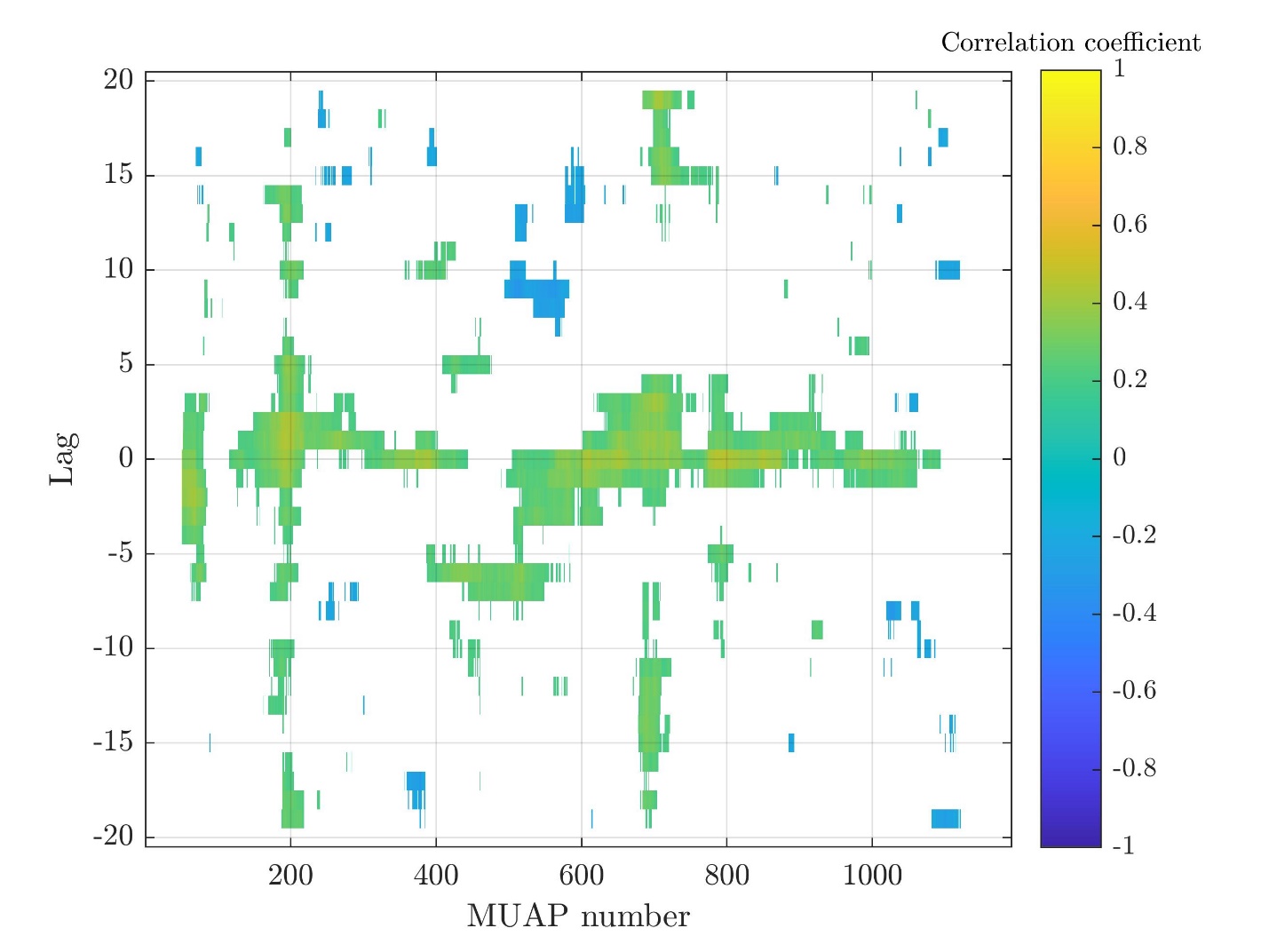

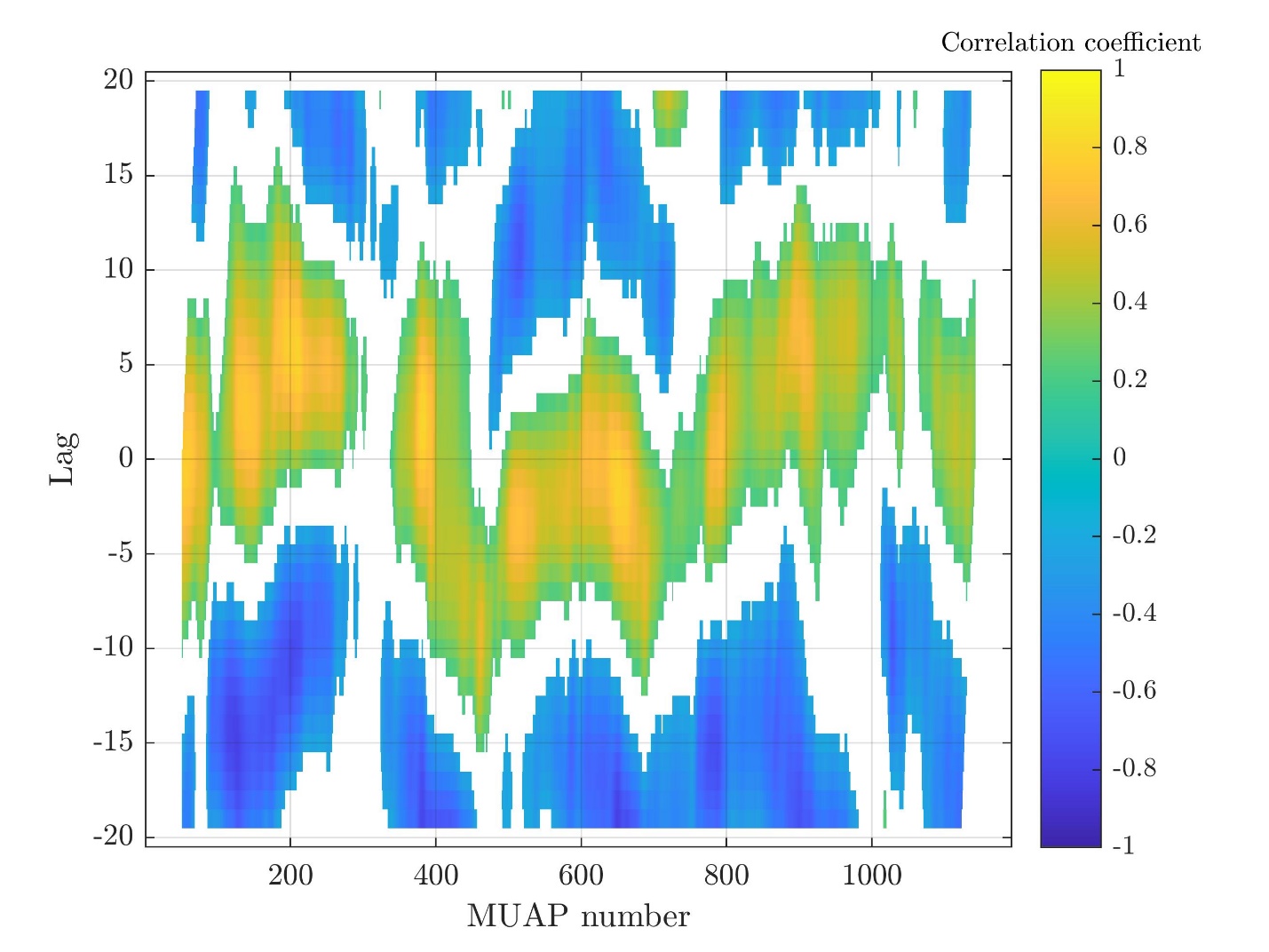

Subject 10

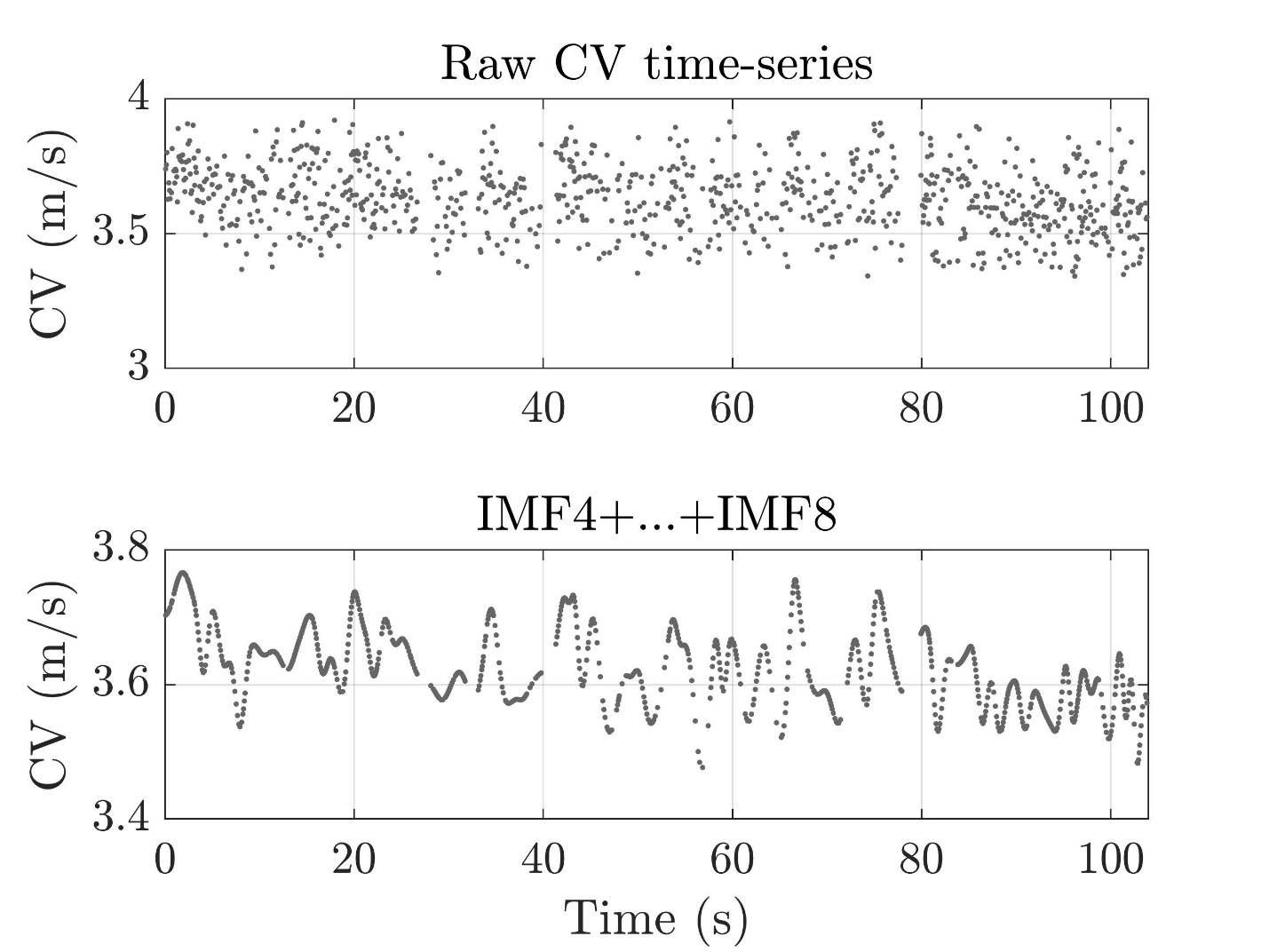

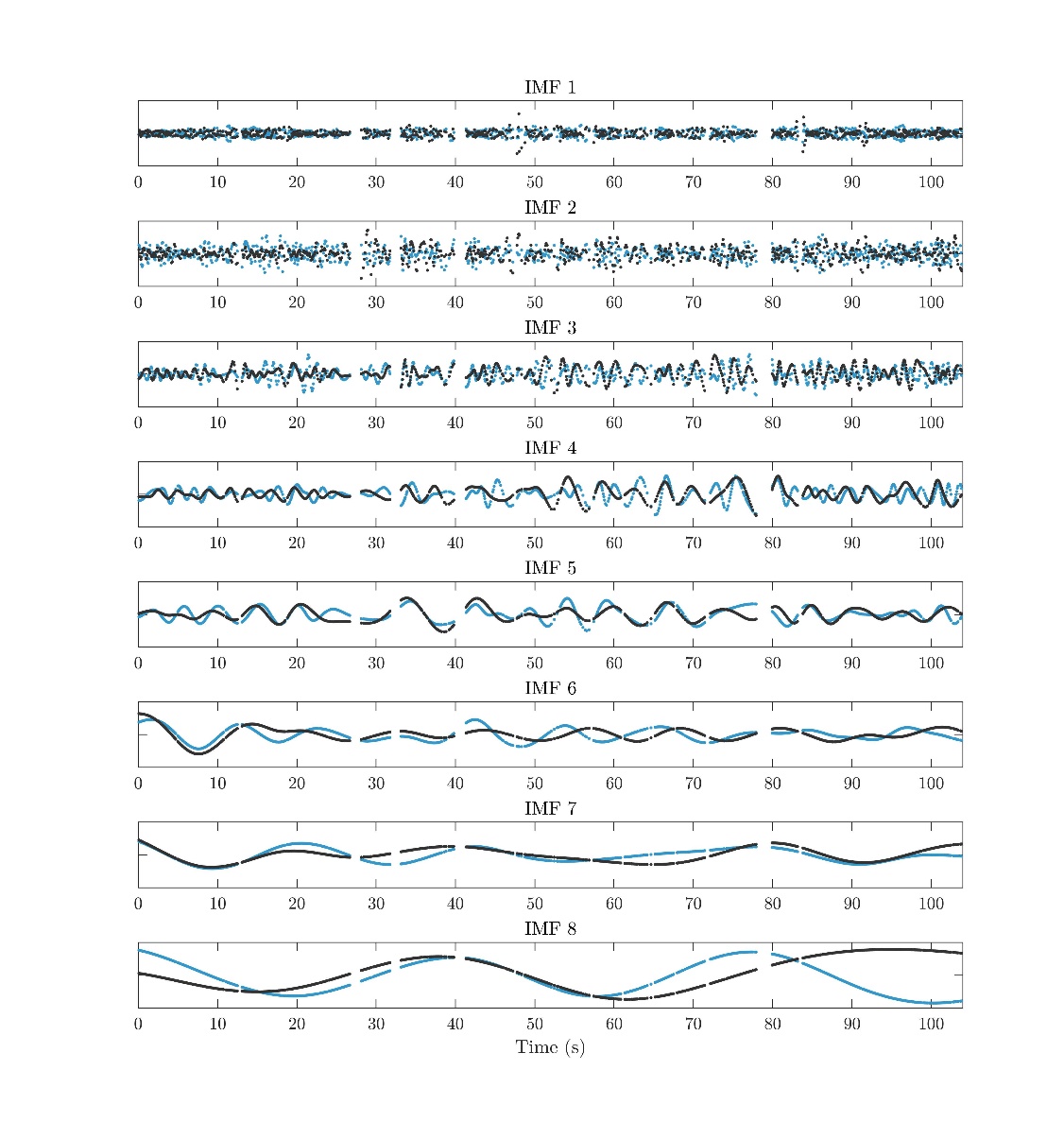

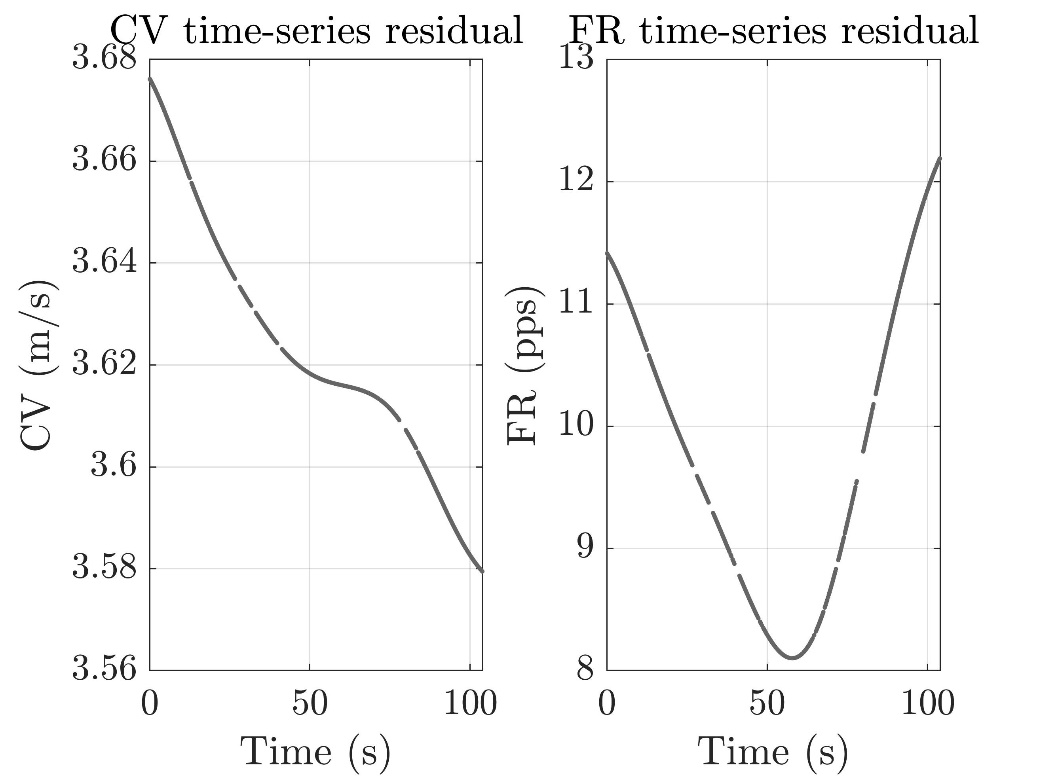

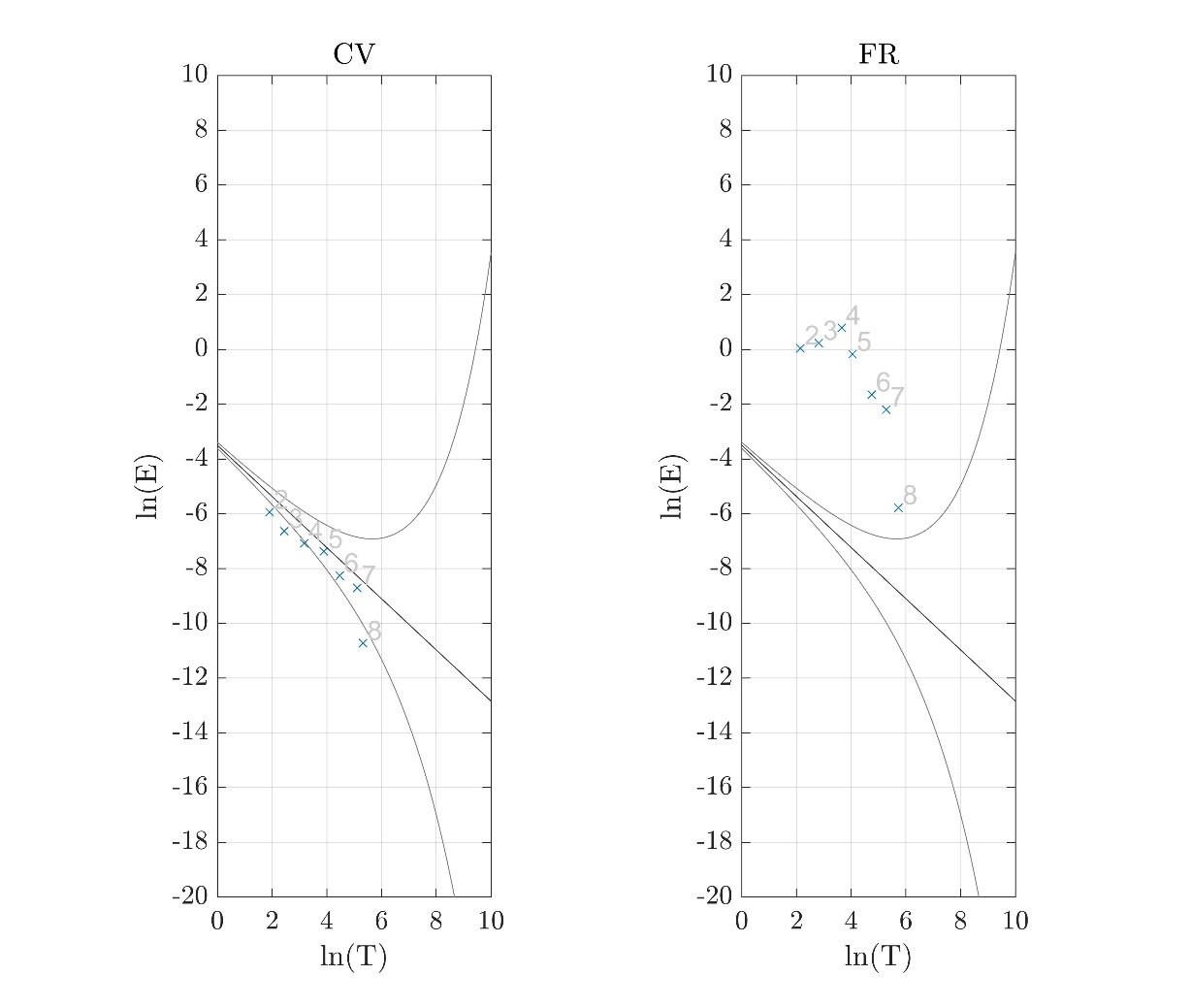

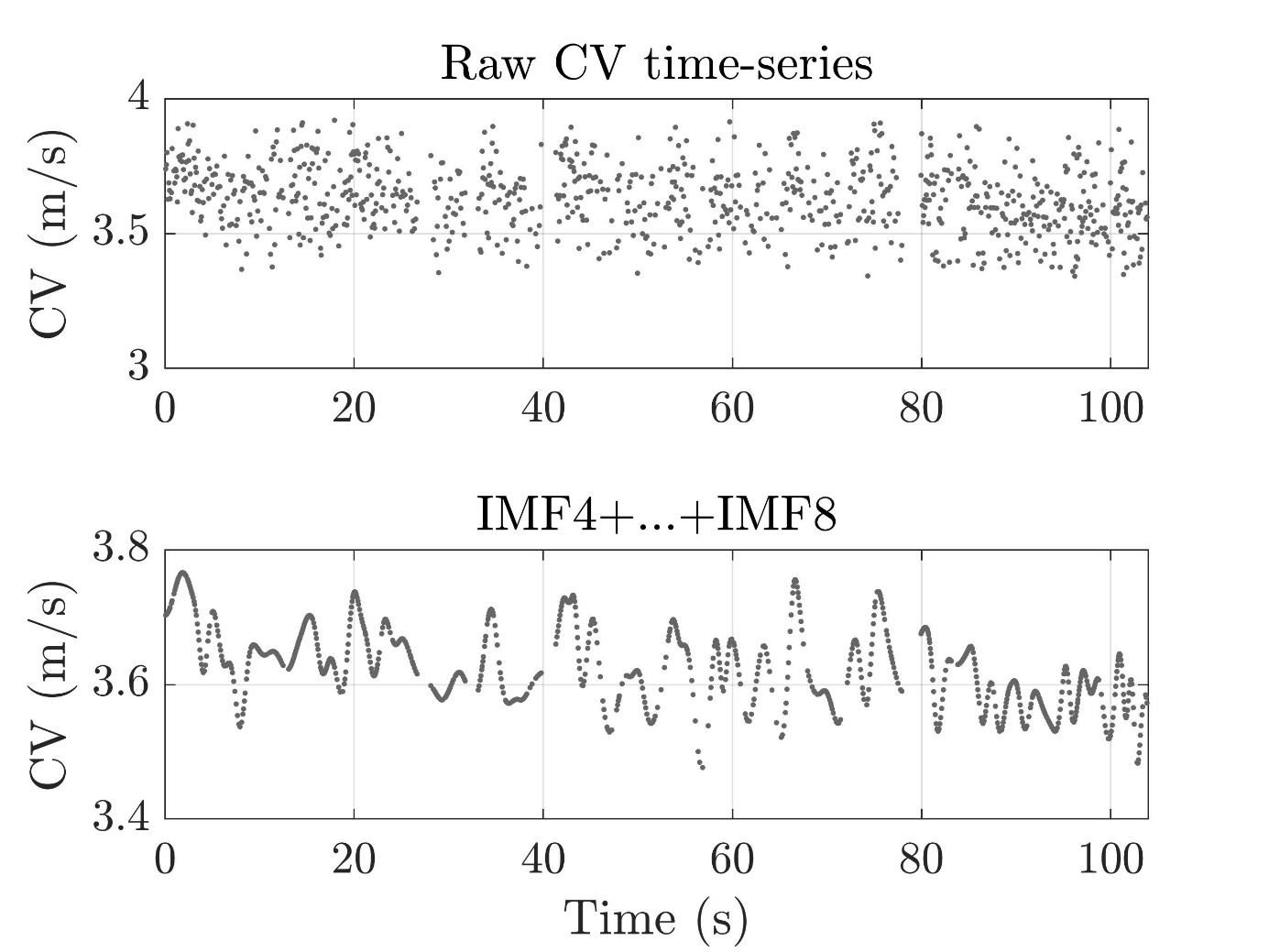

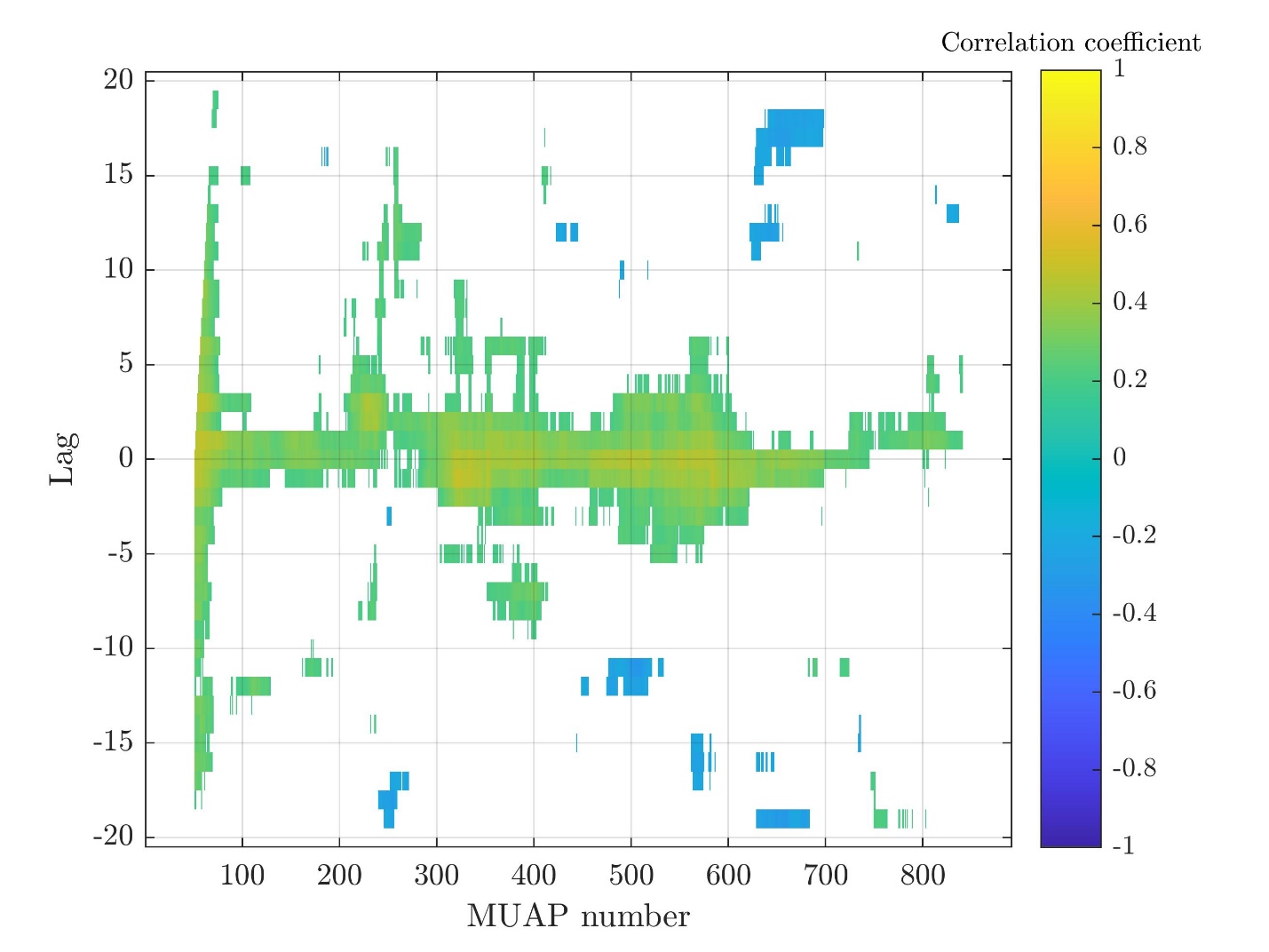

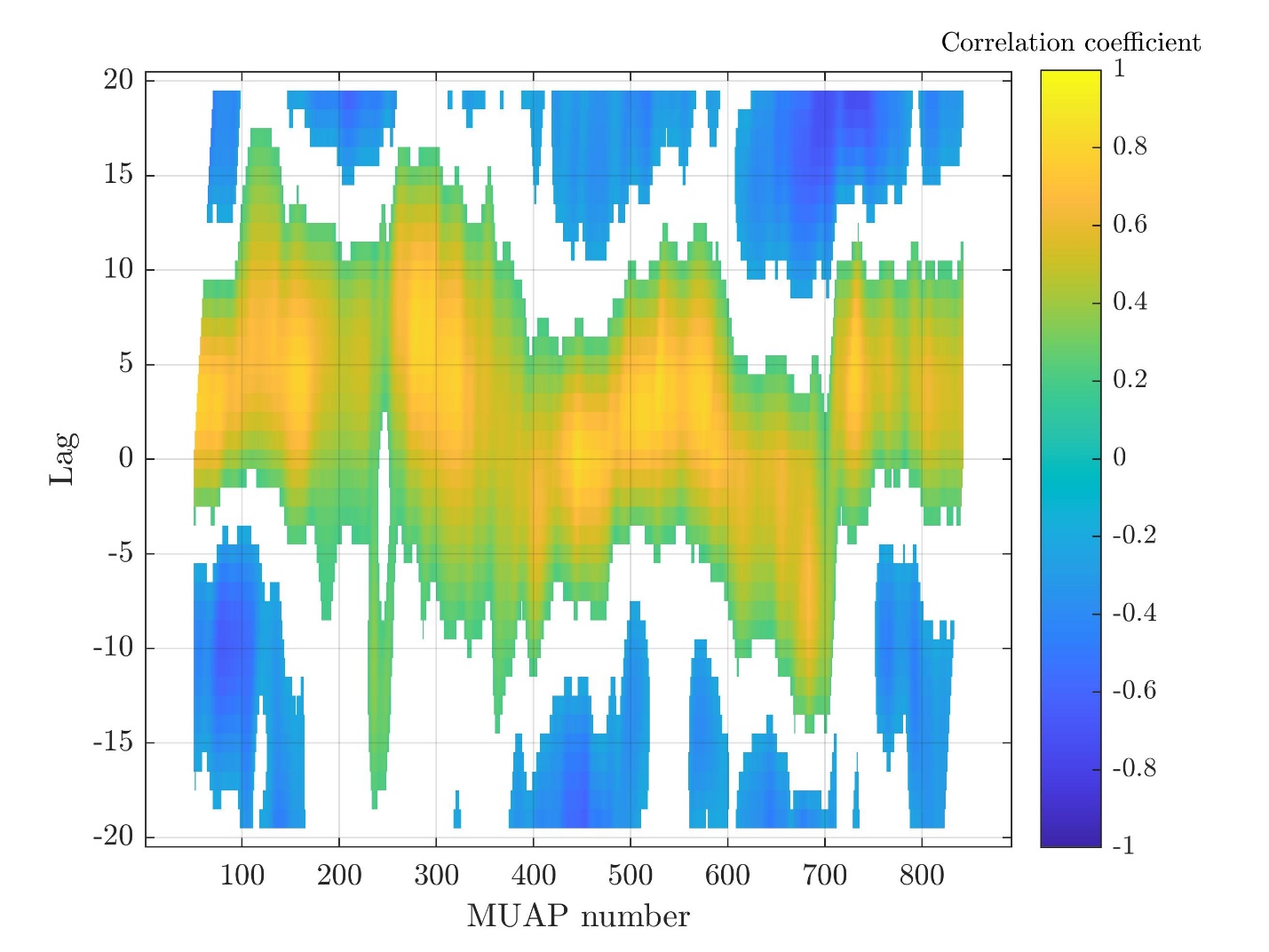

Subject 11

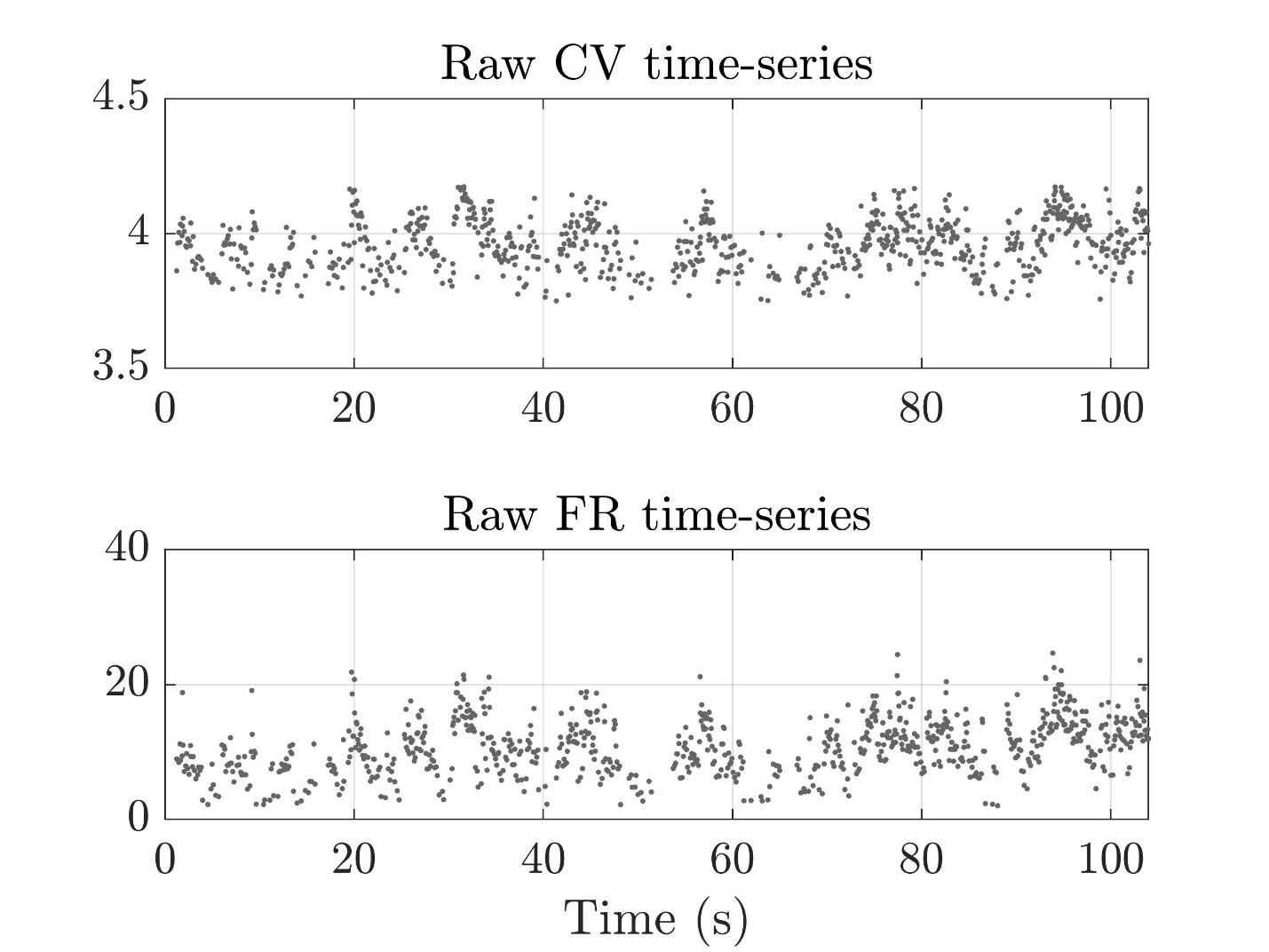

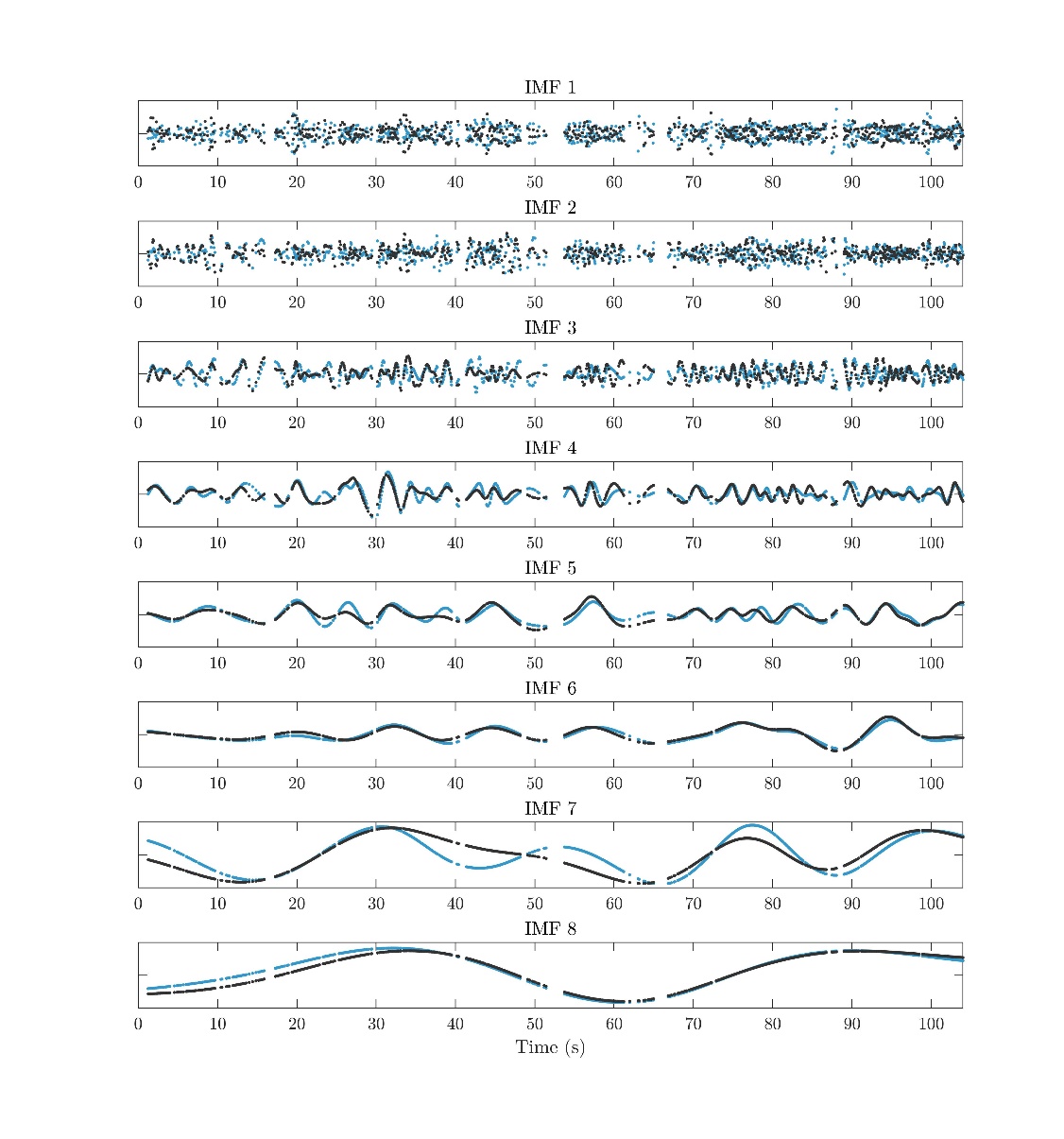

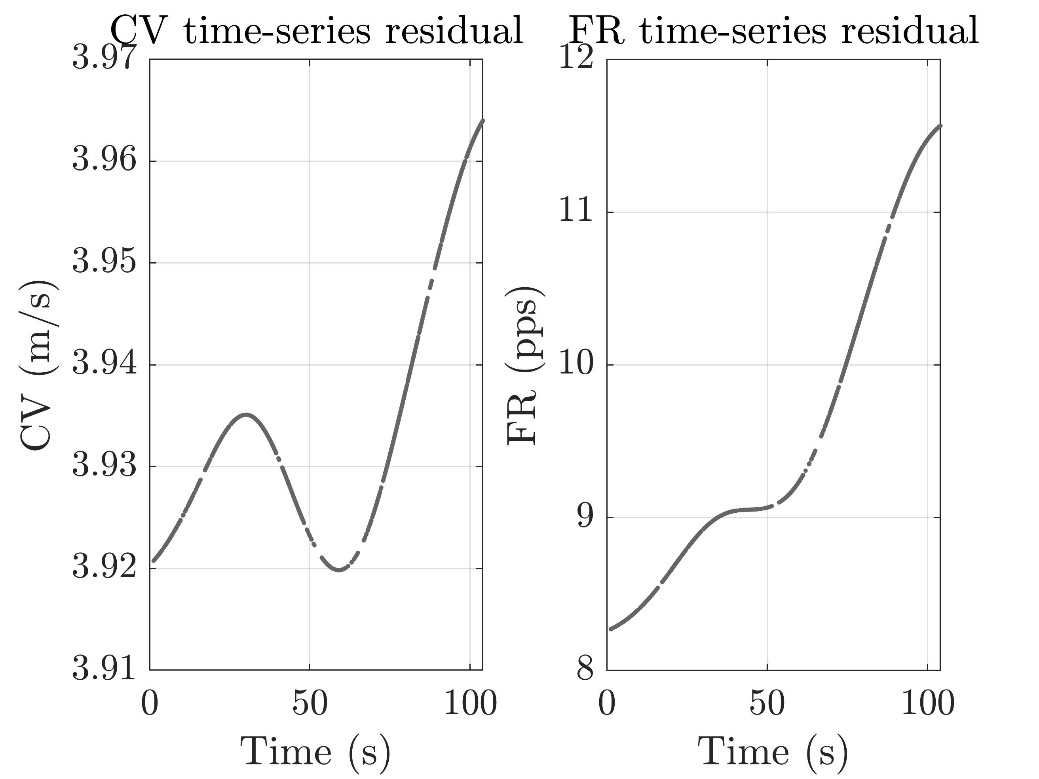

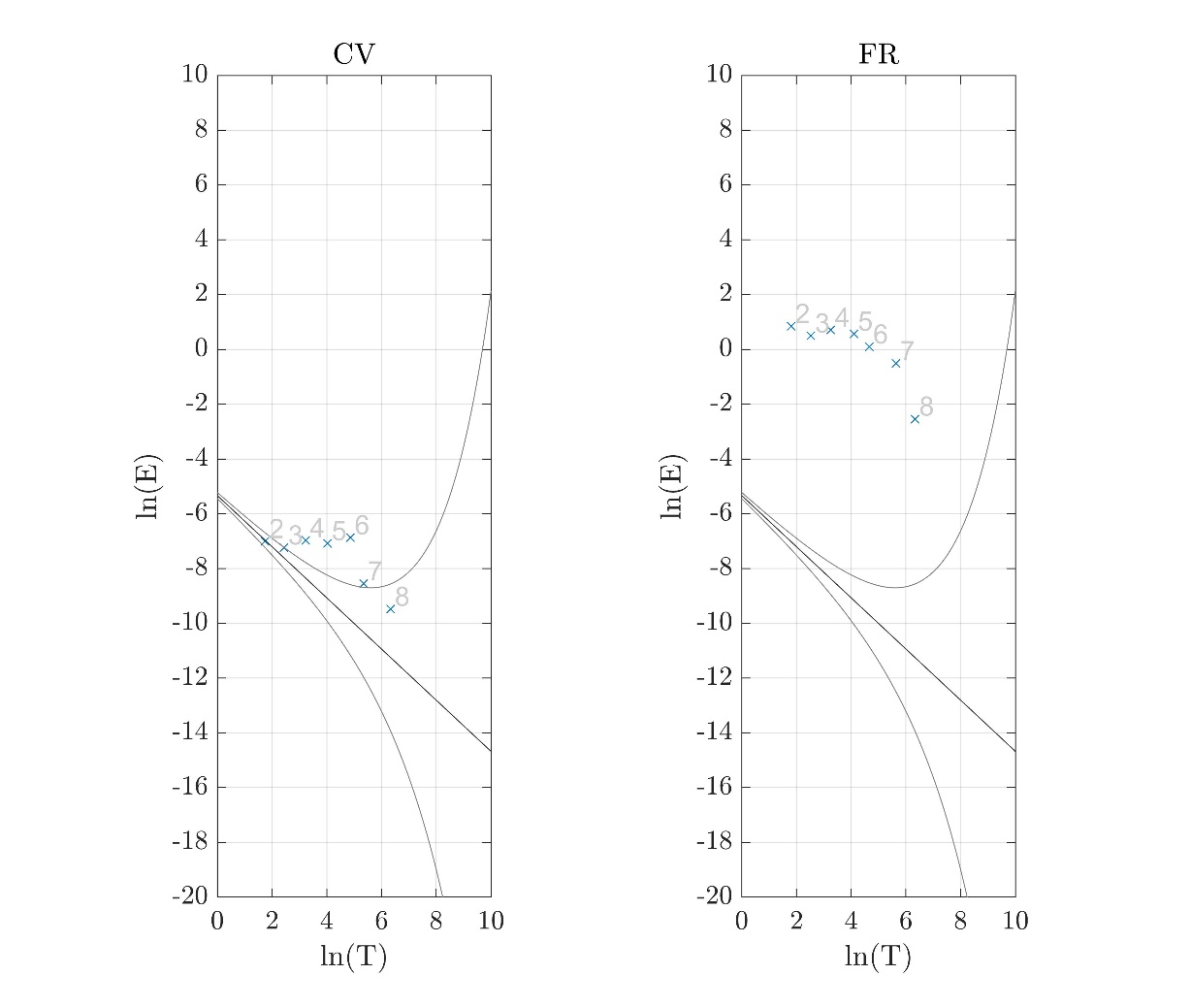

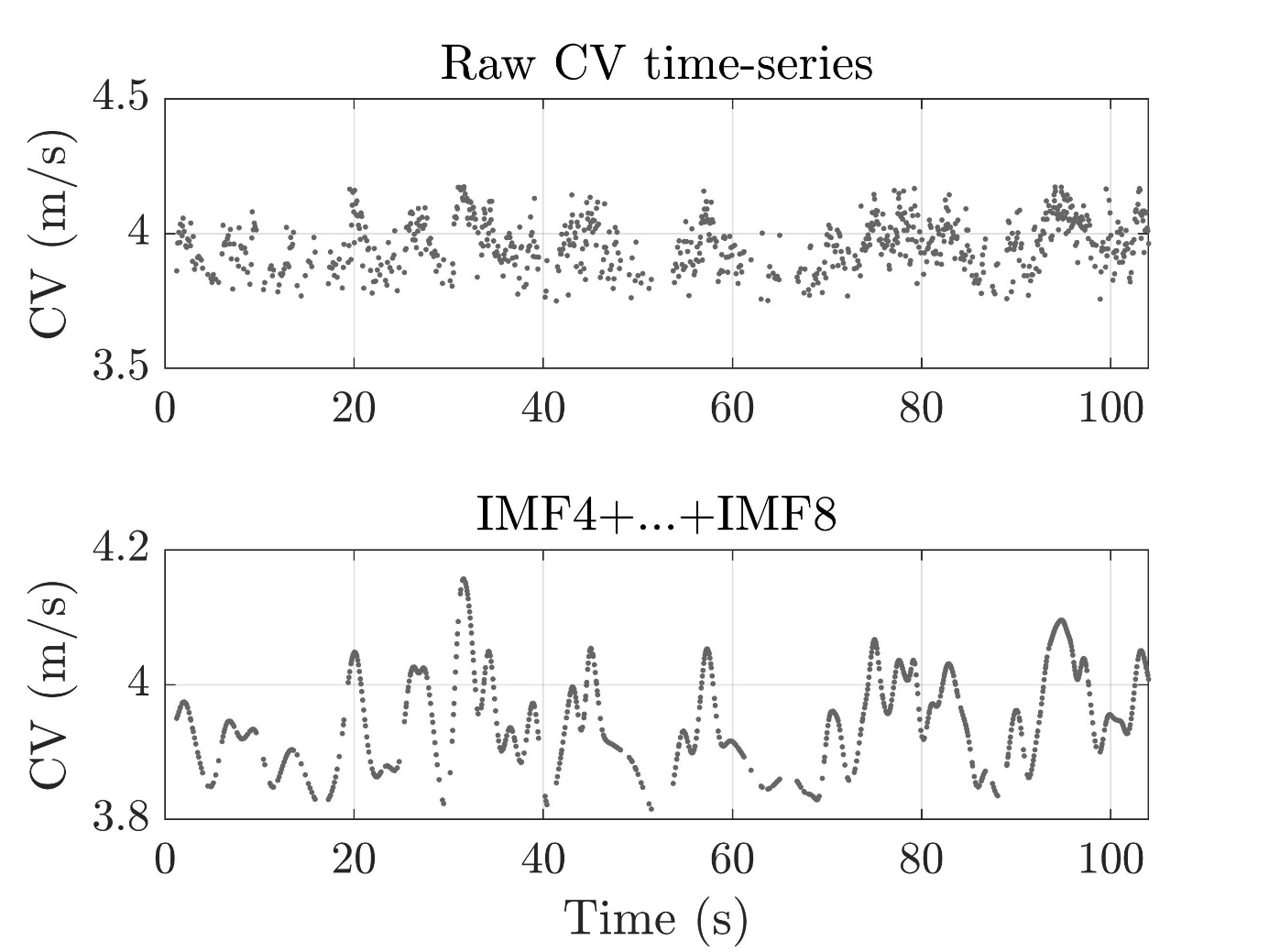

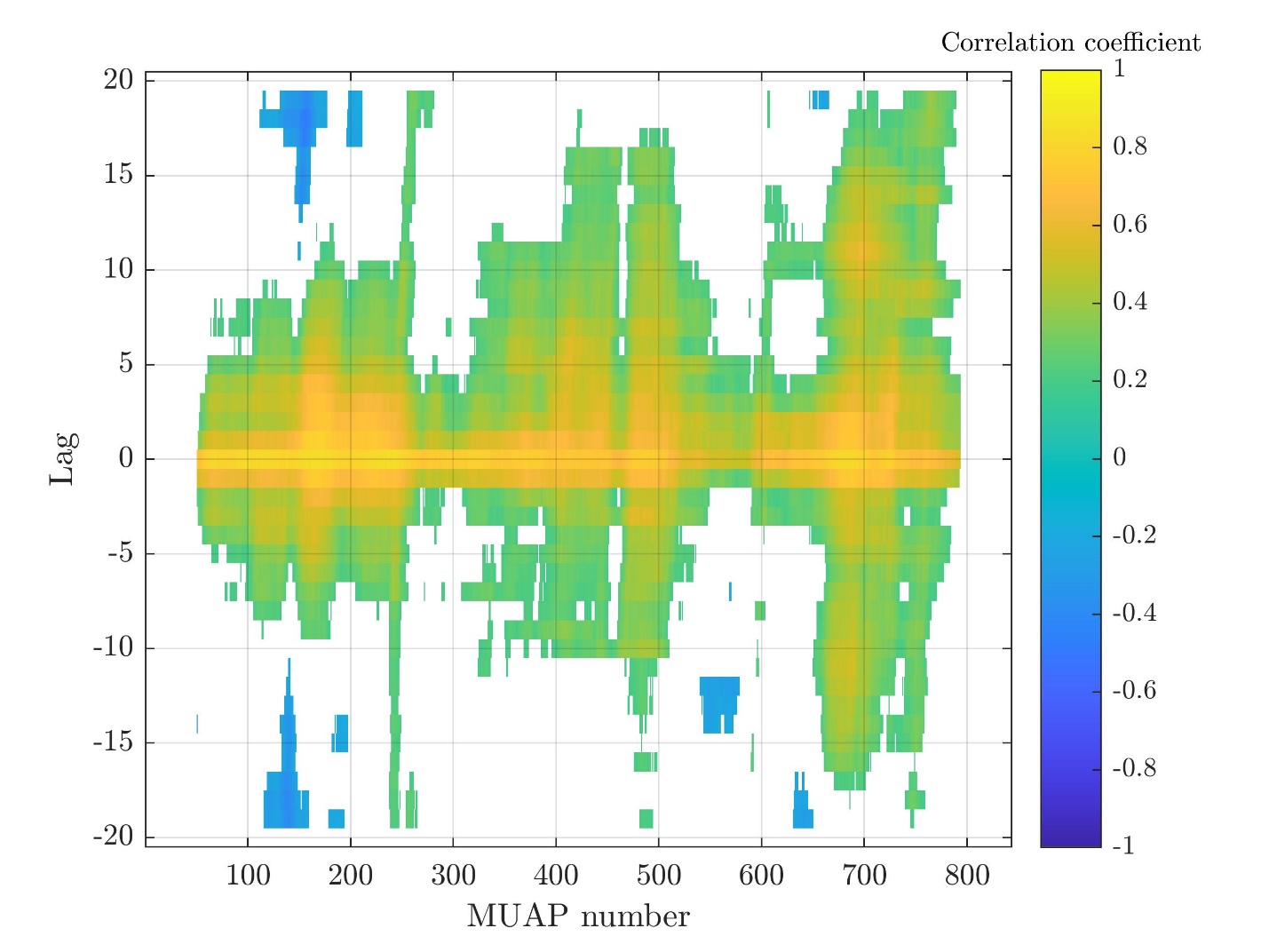

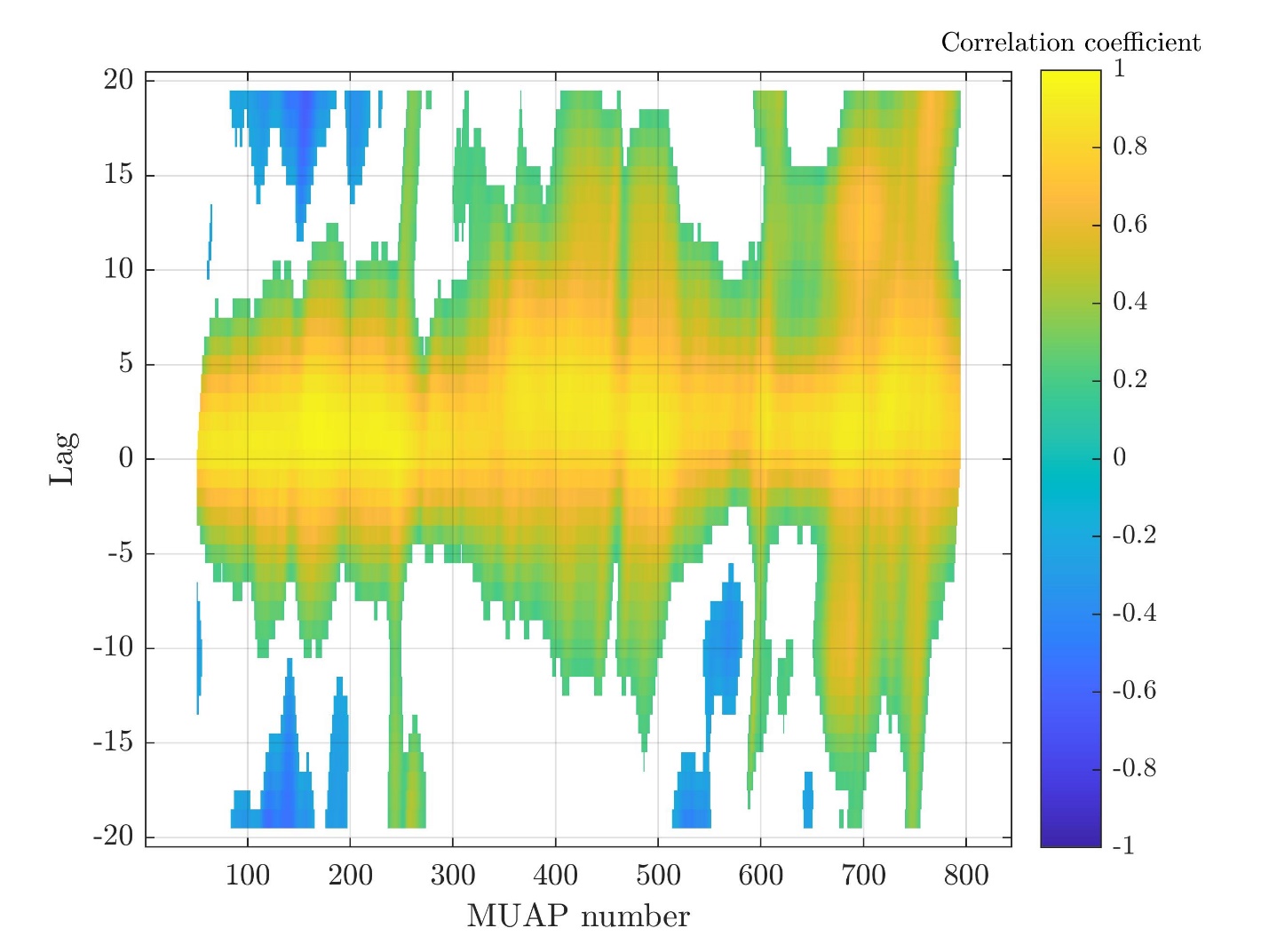

Subject 12

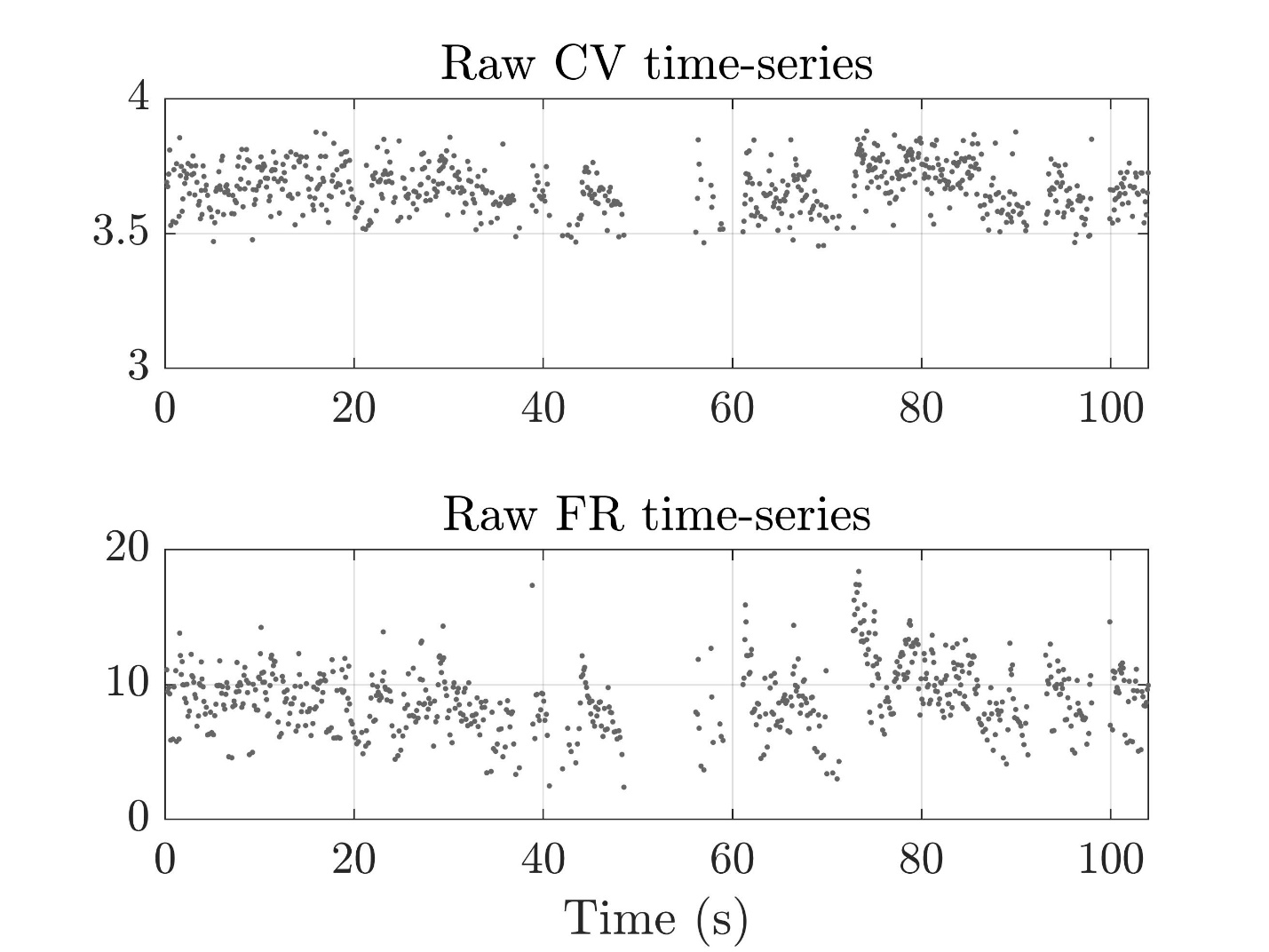

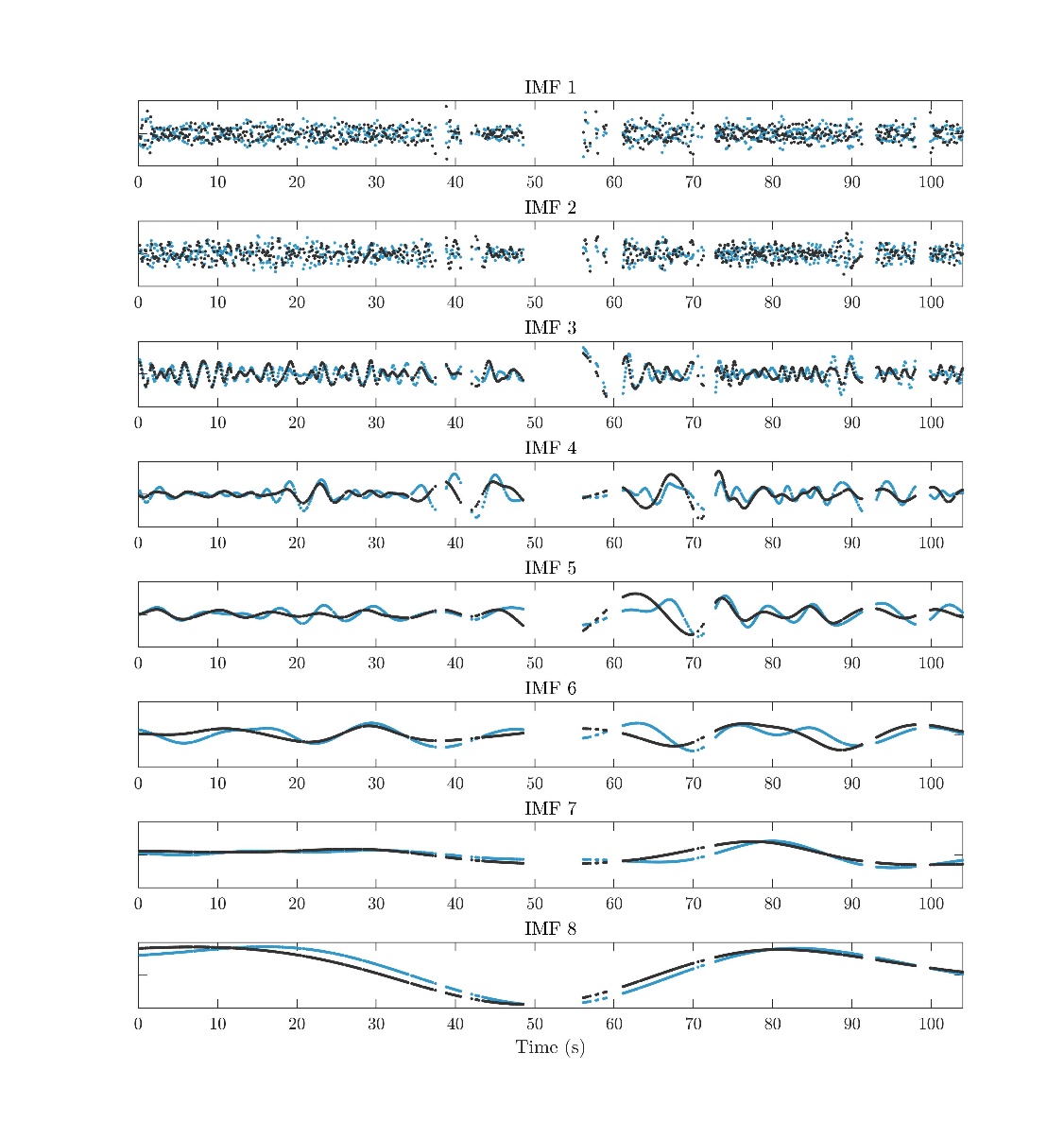

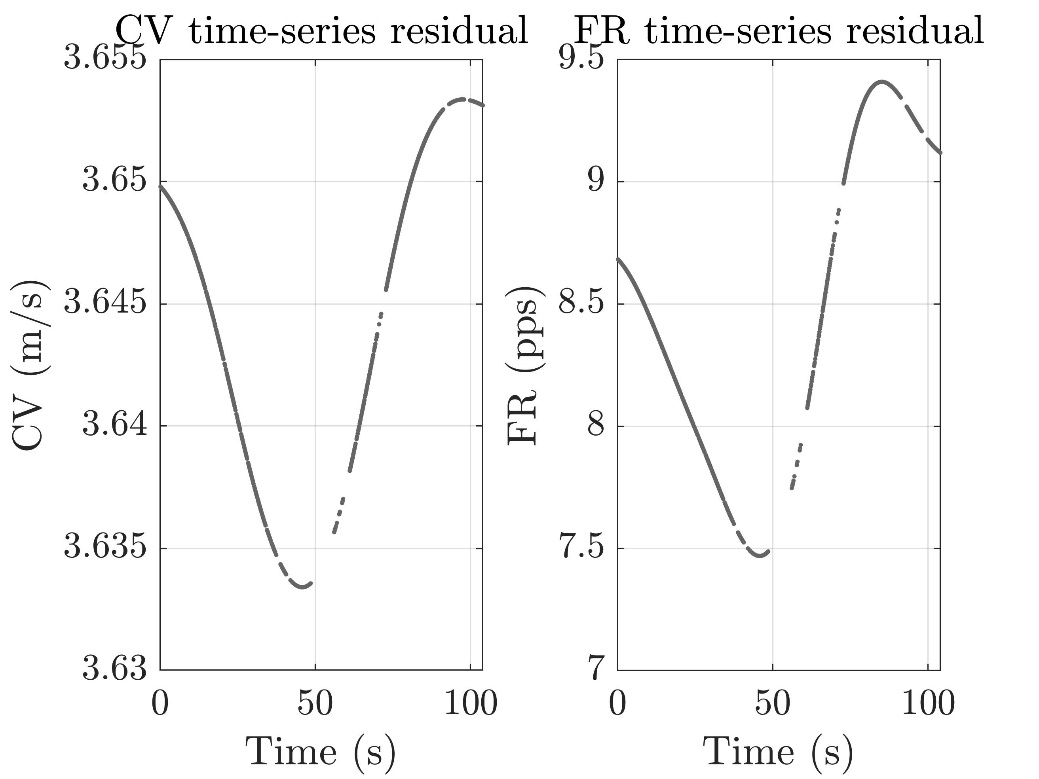

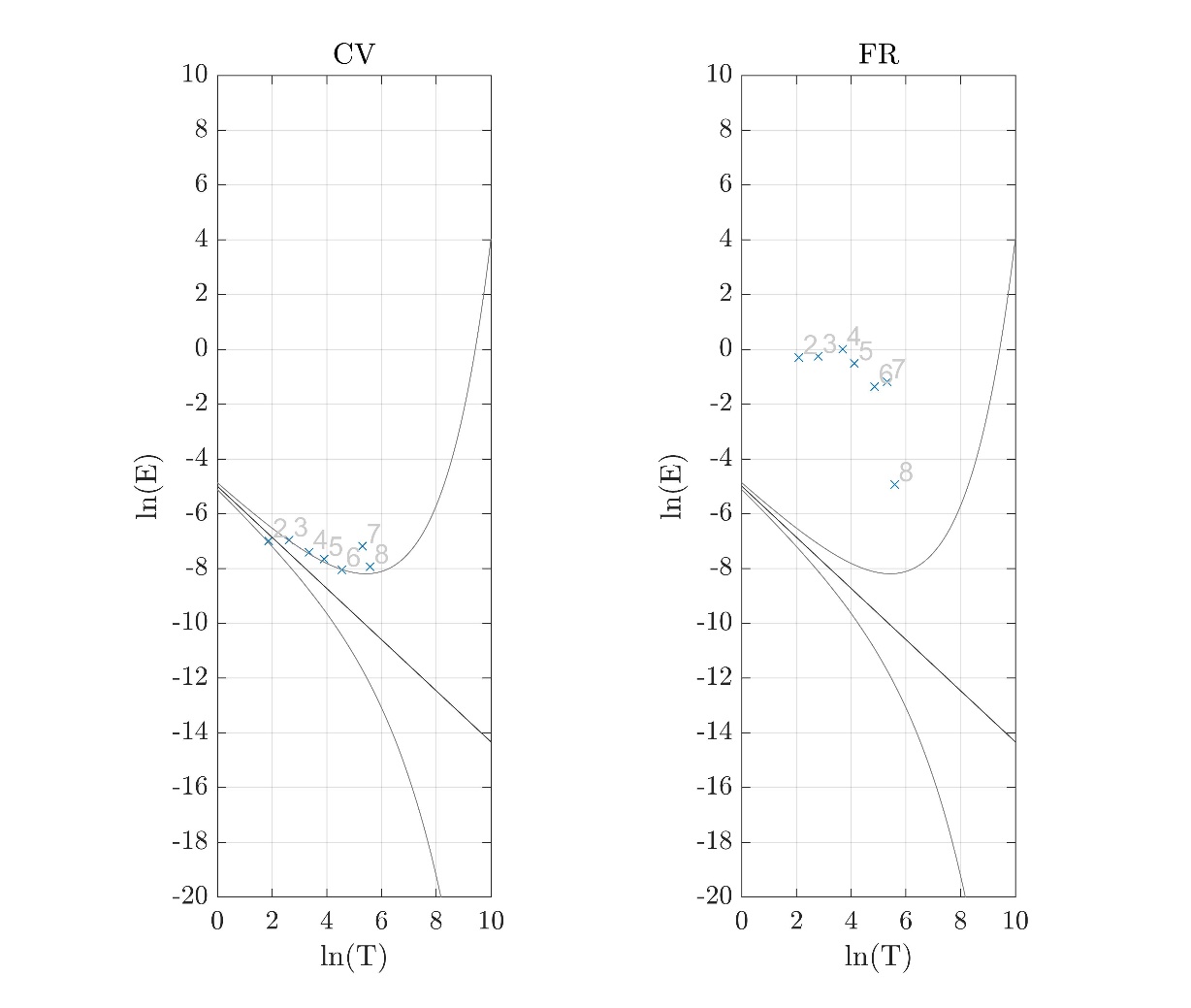

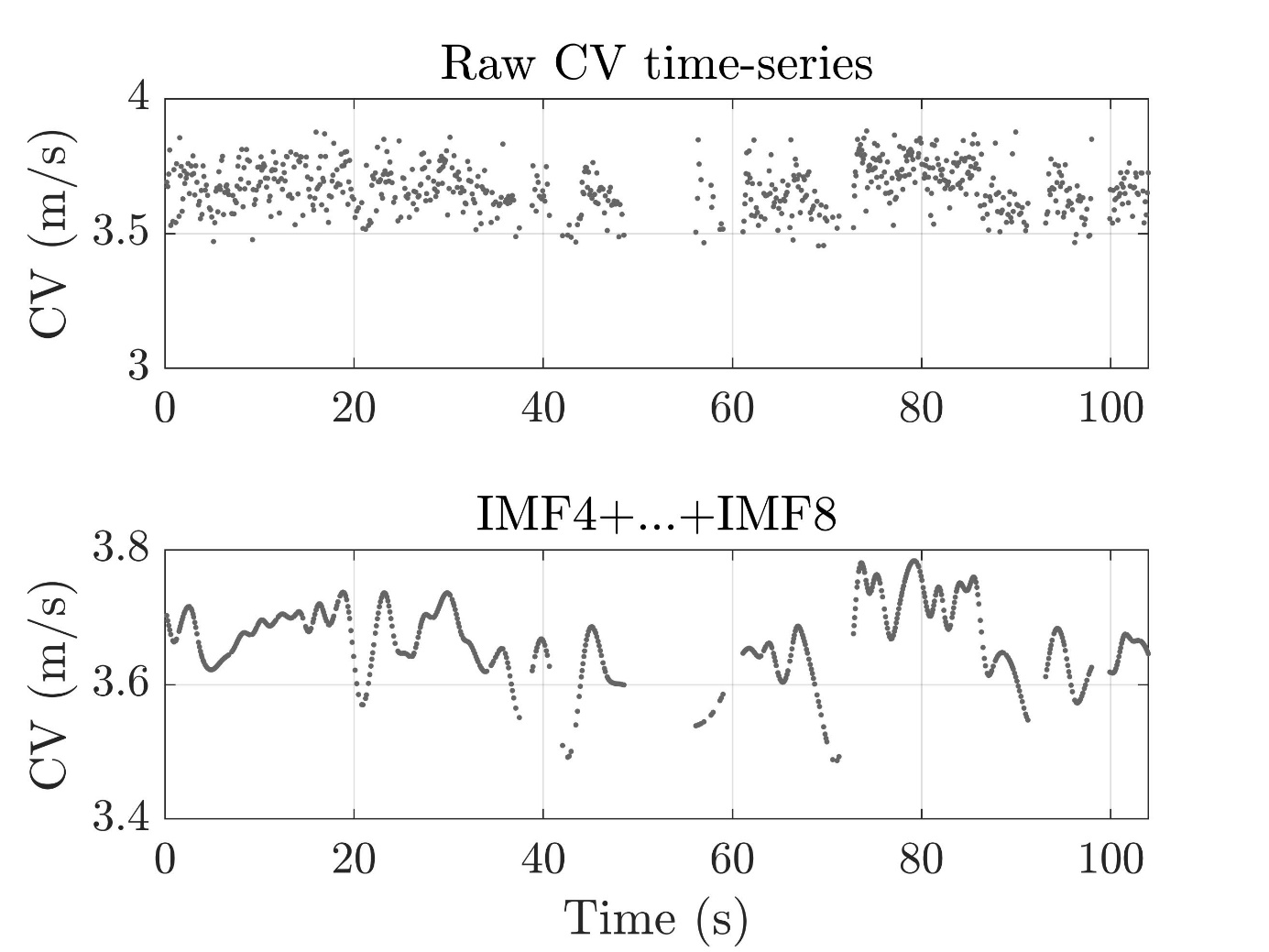

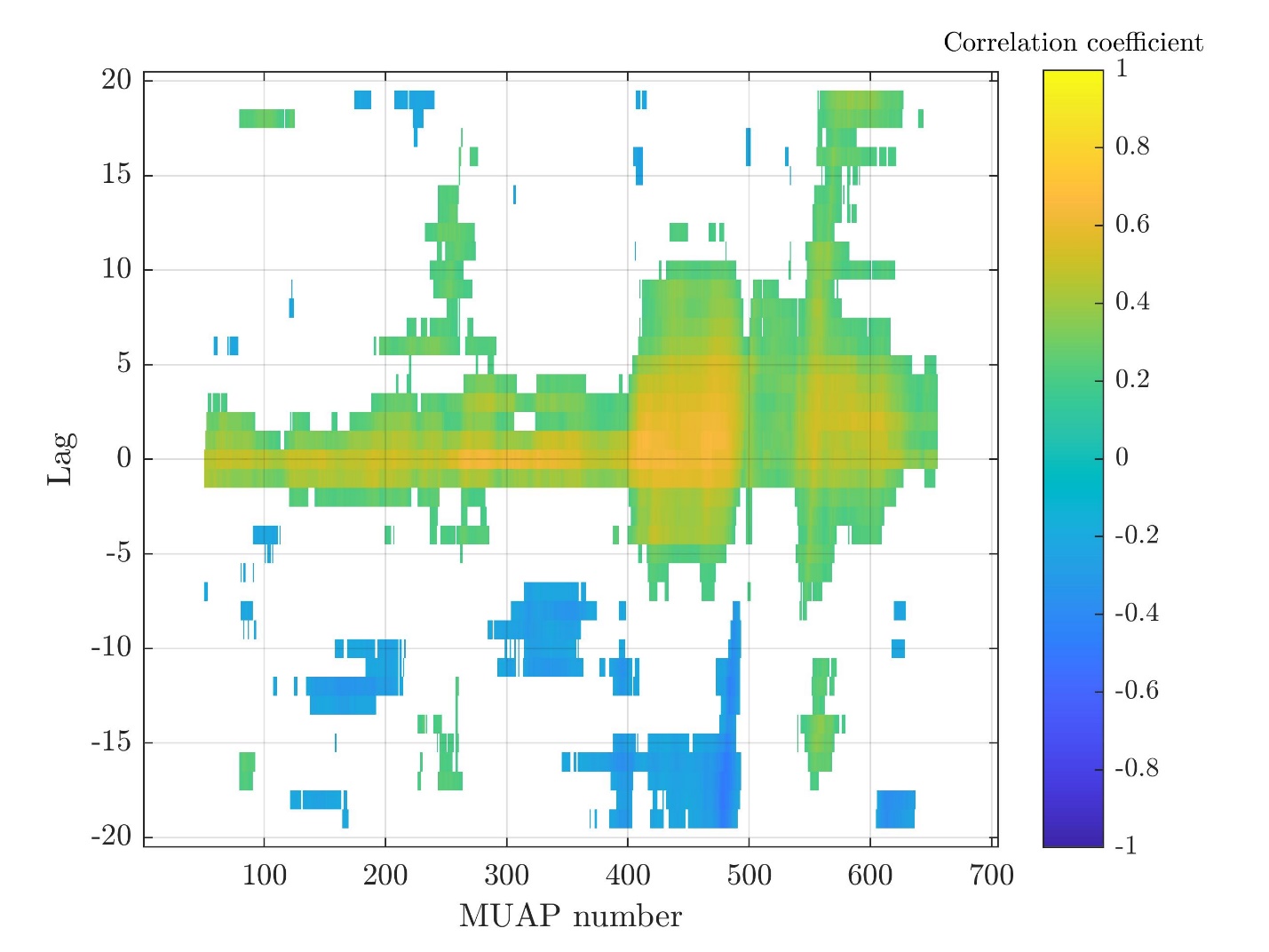

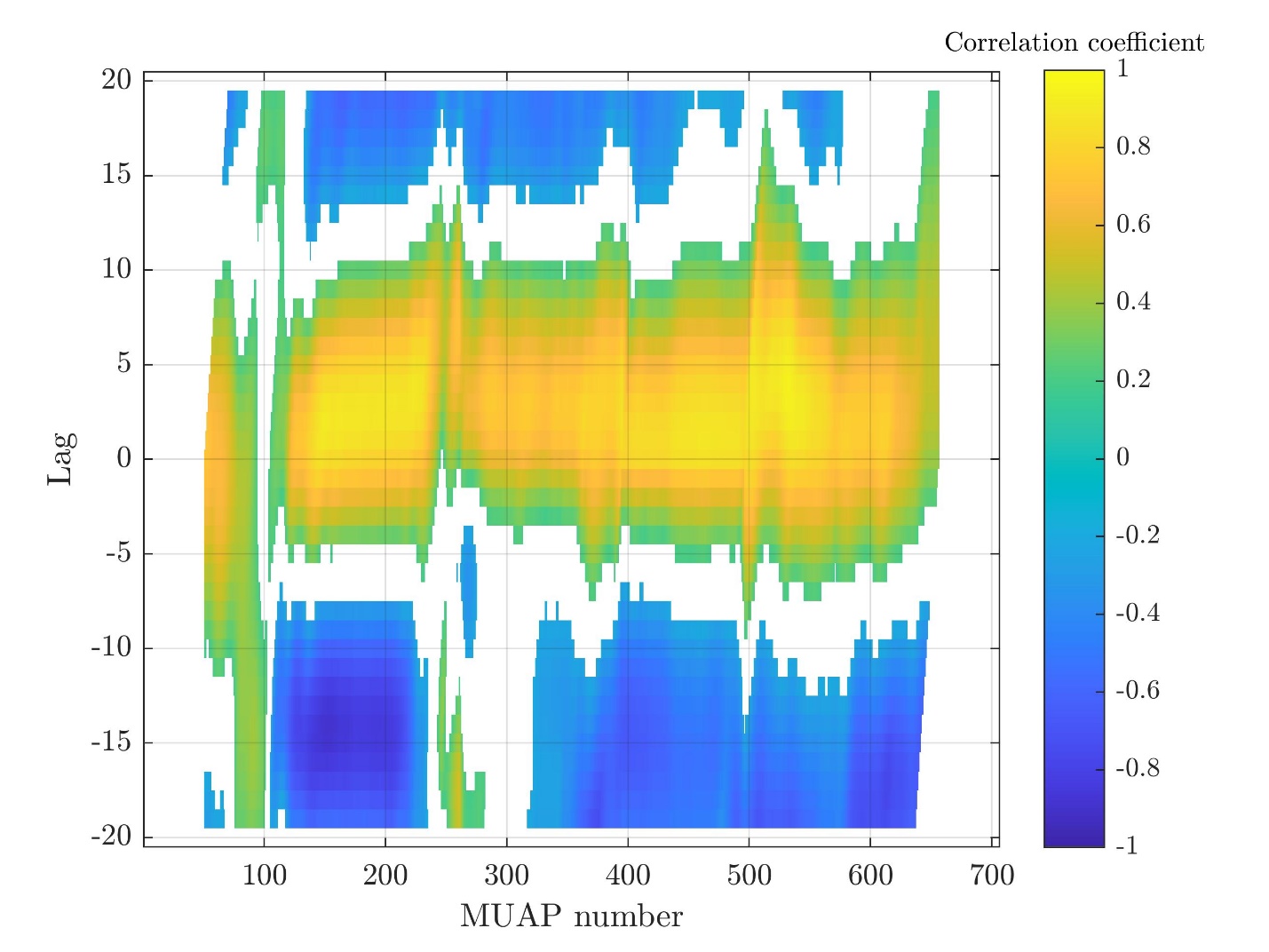

Subject 13

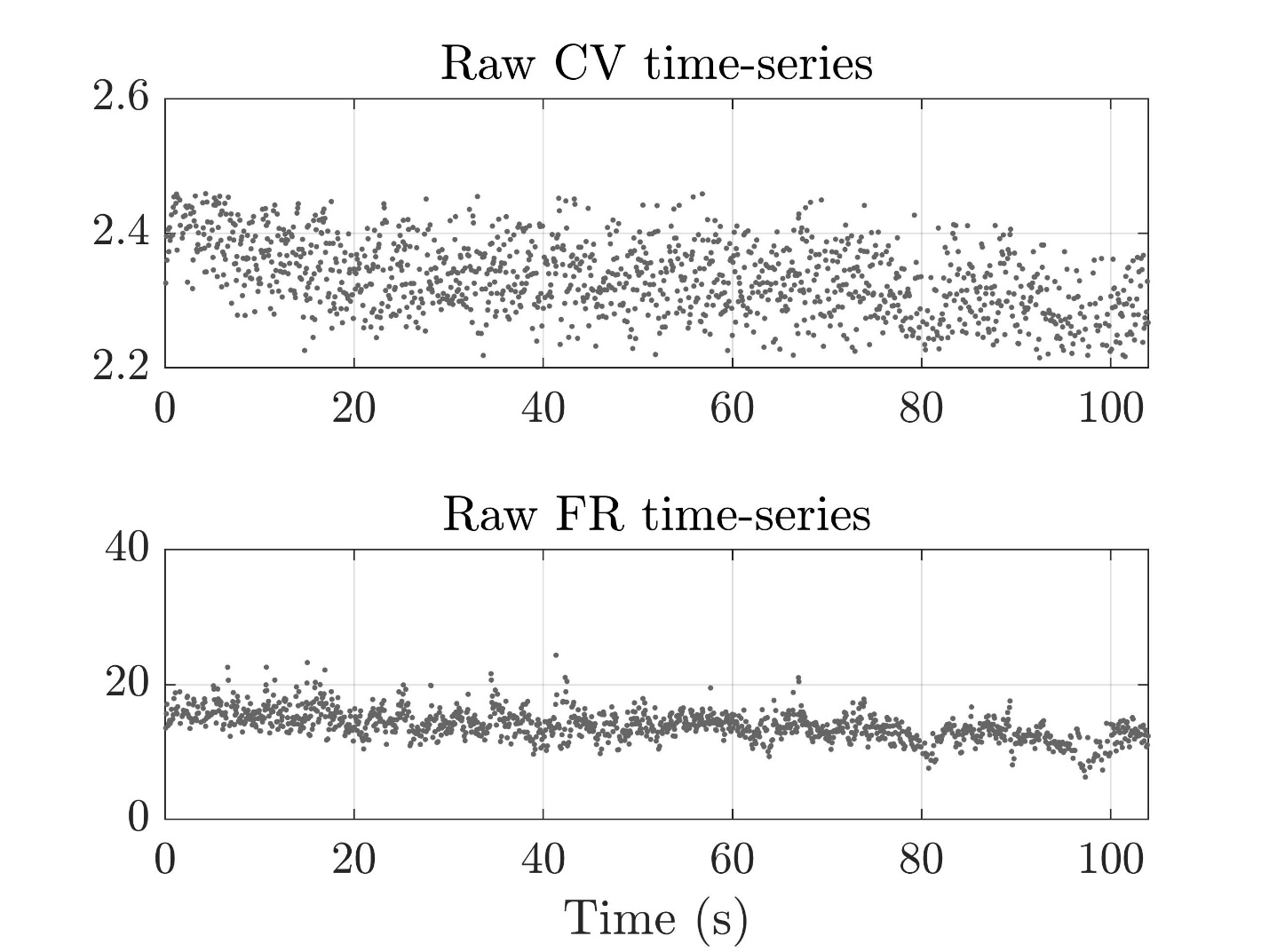

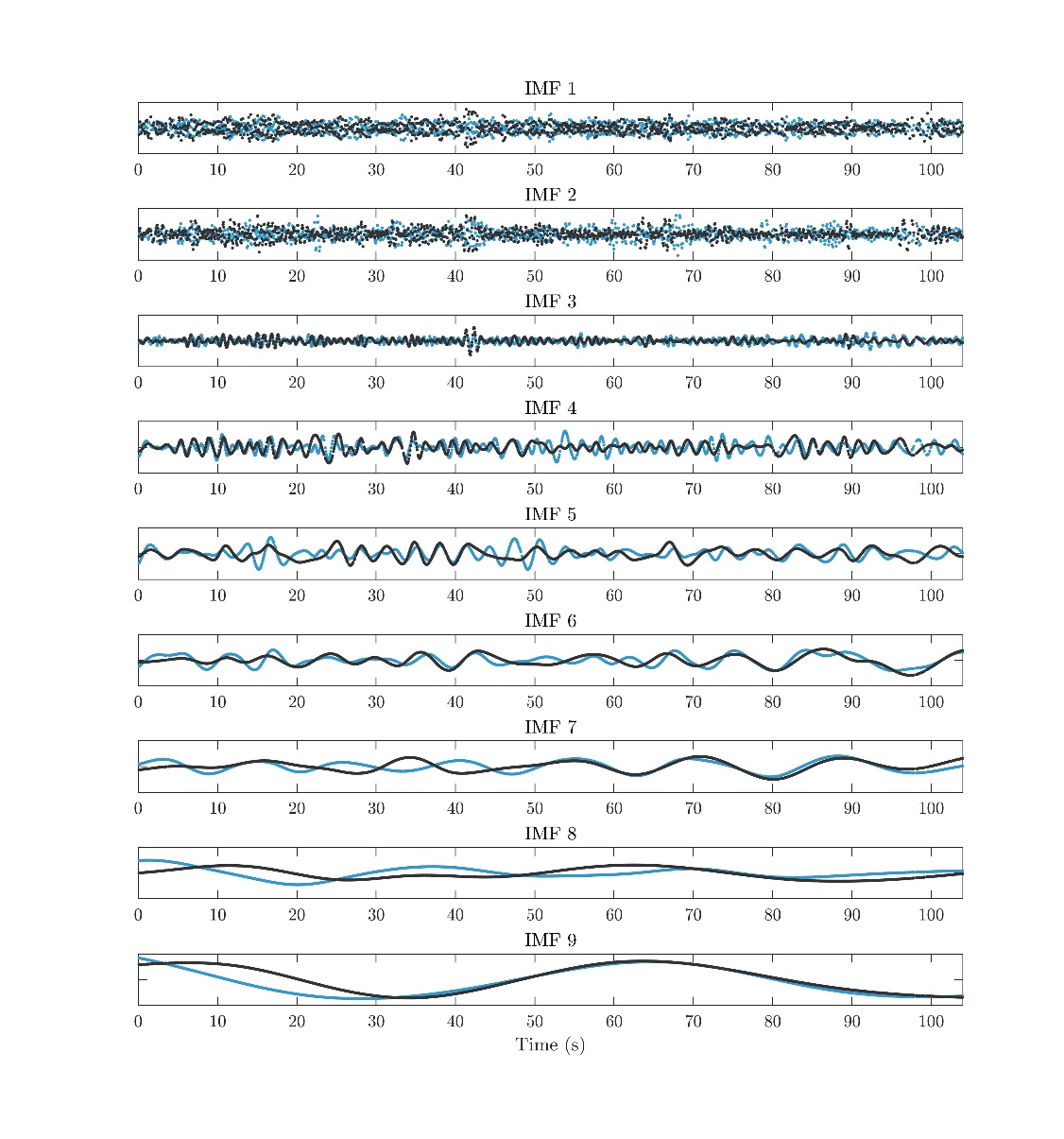

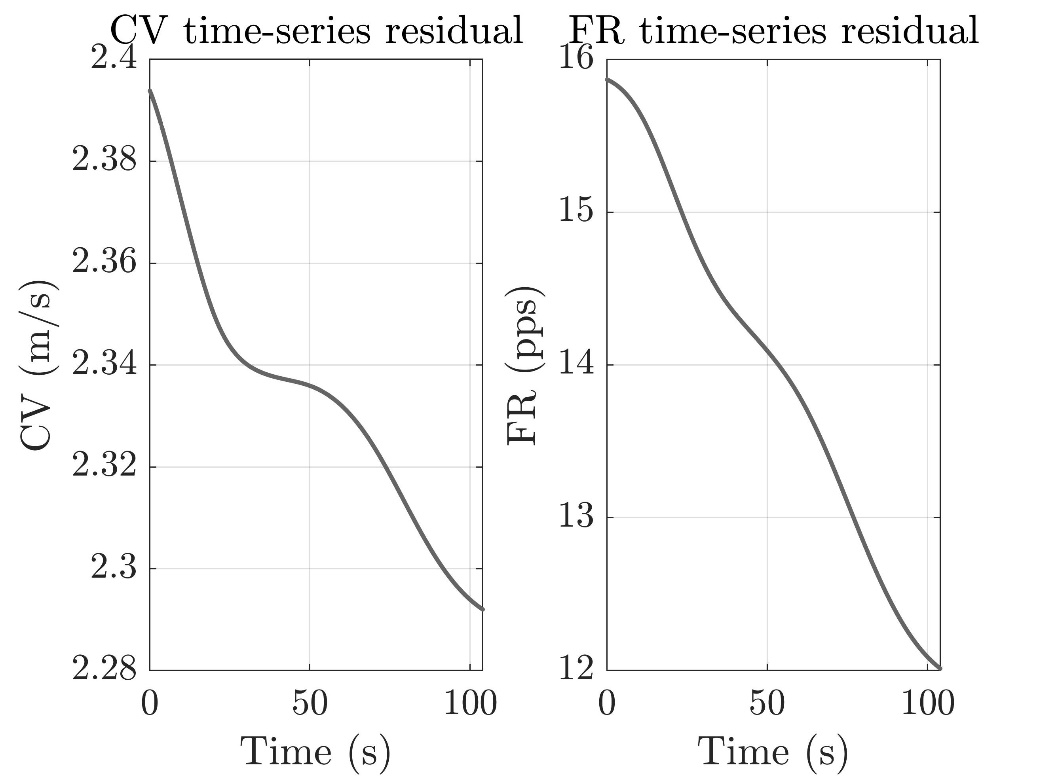

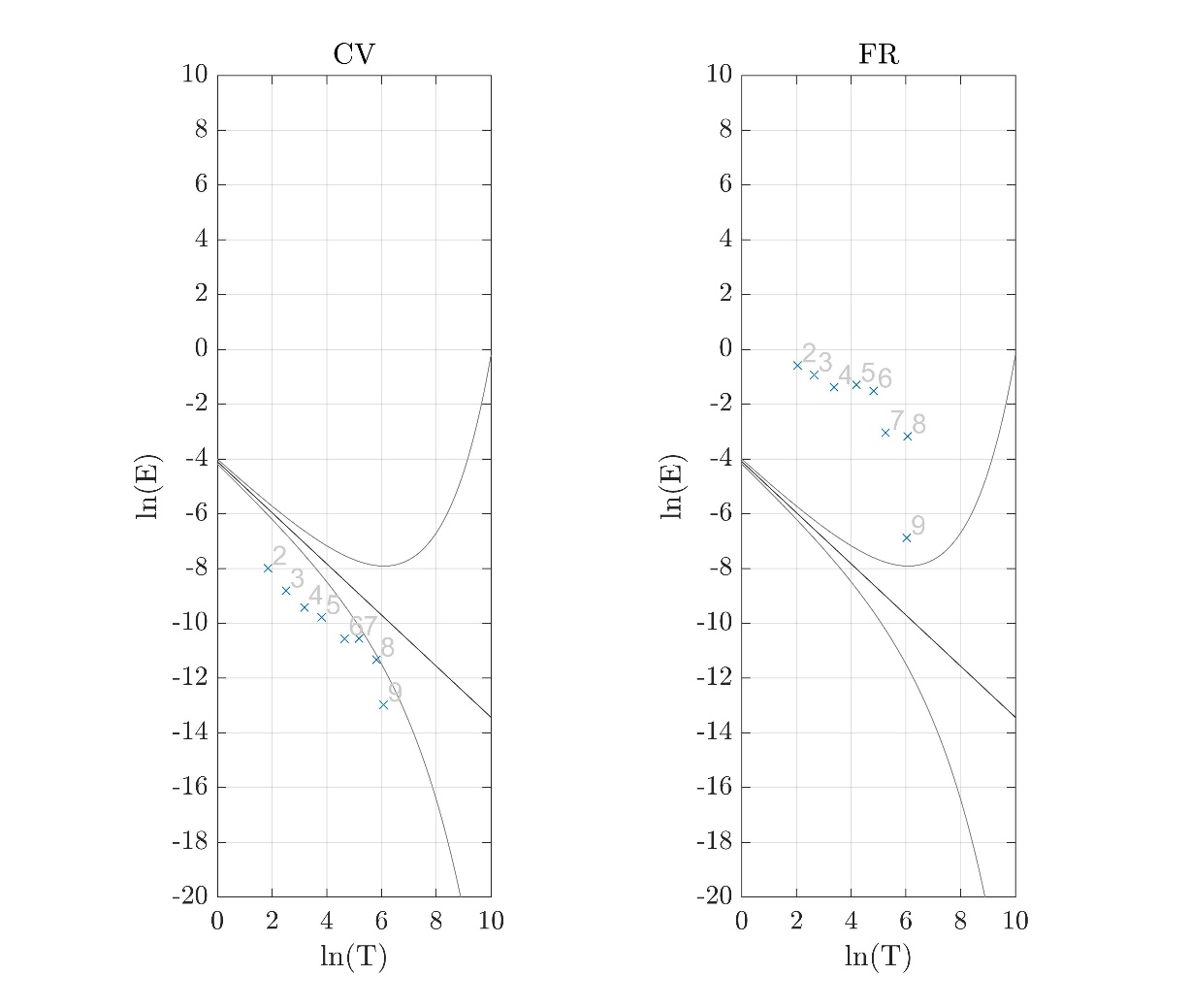

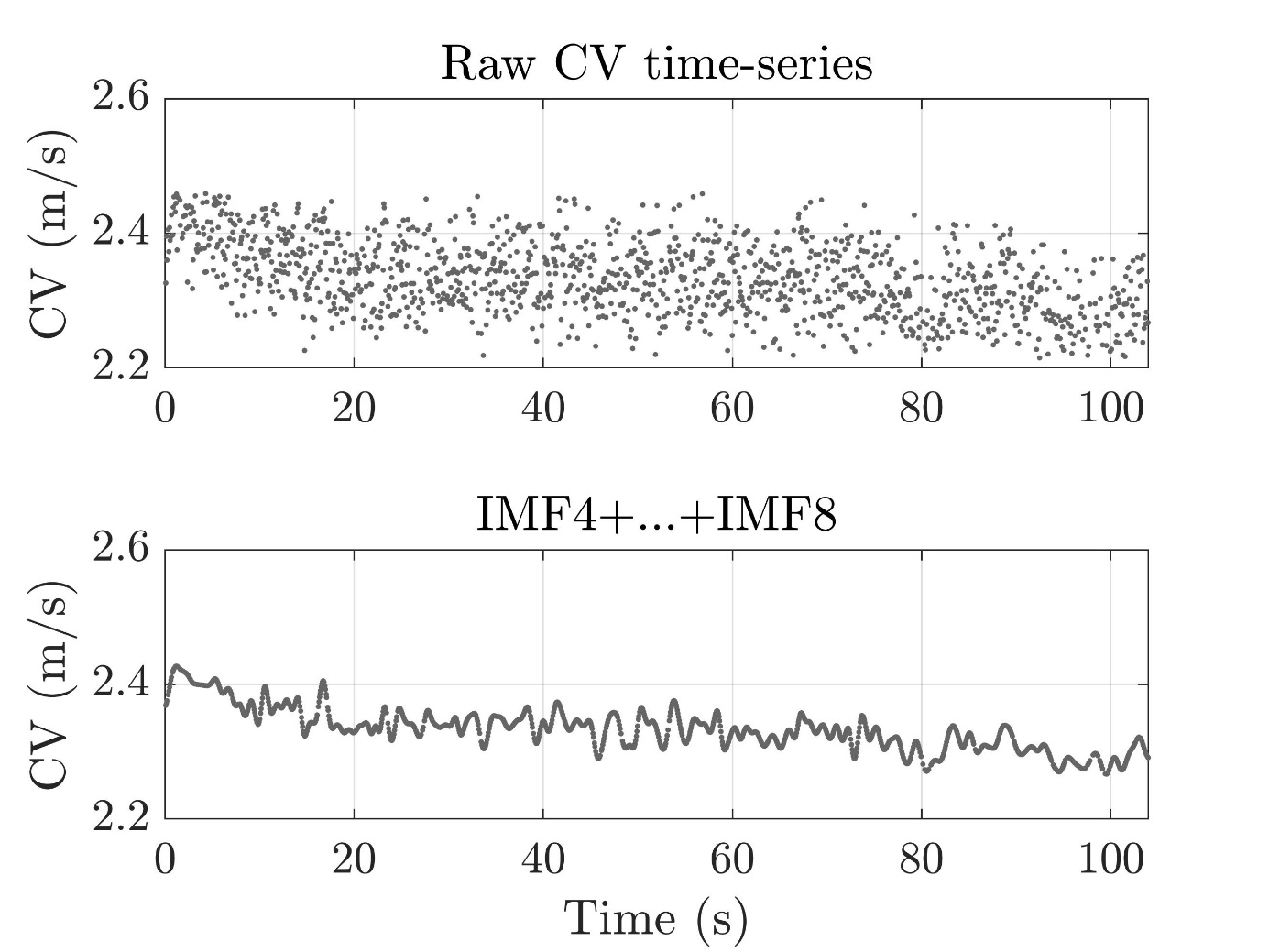

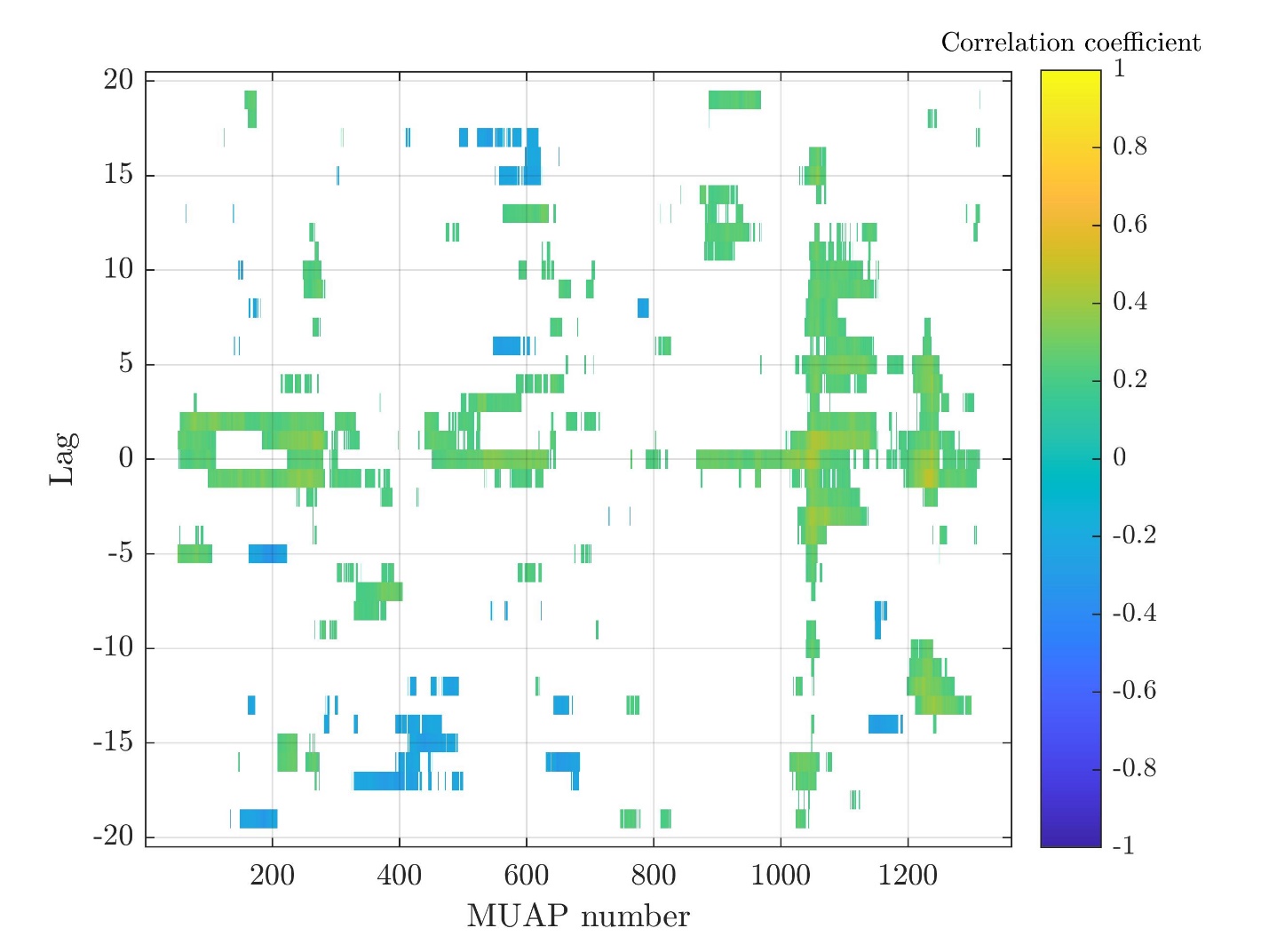

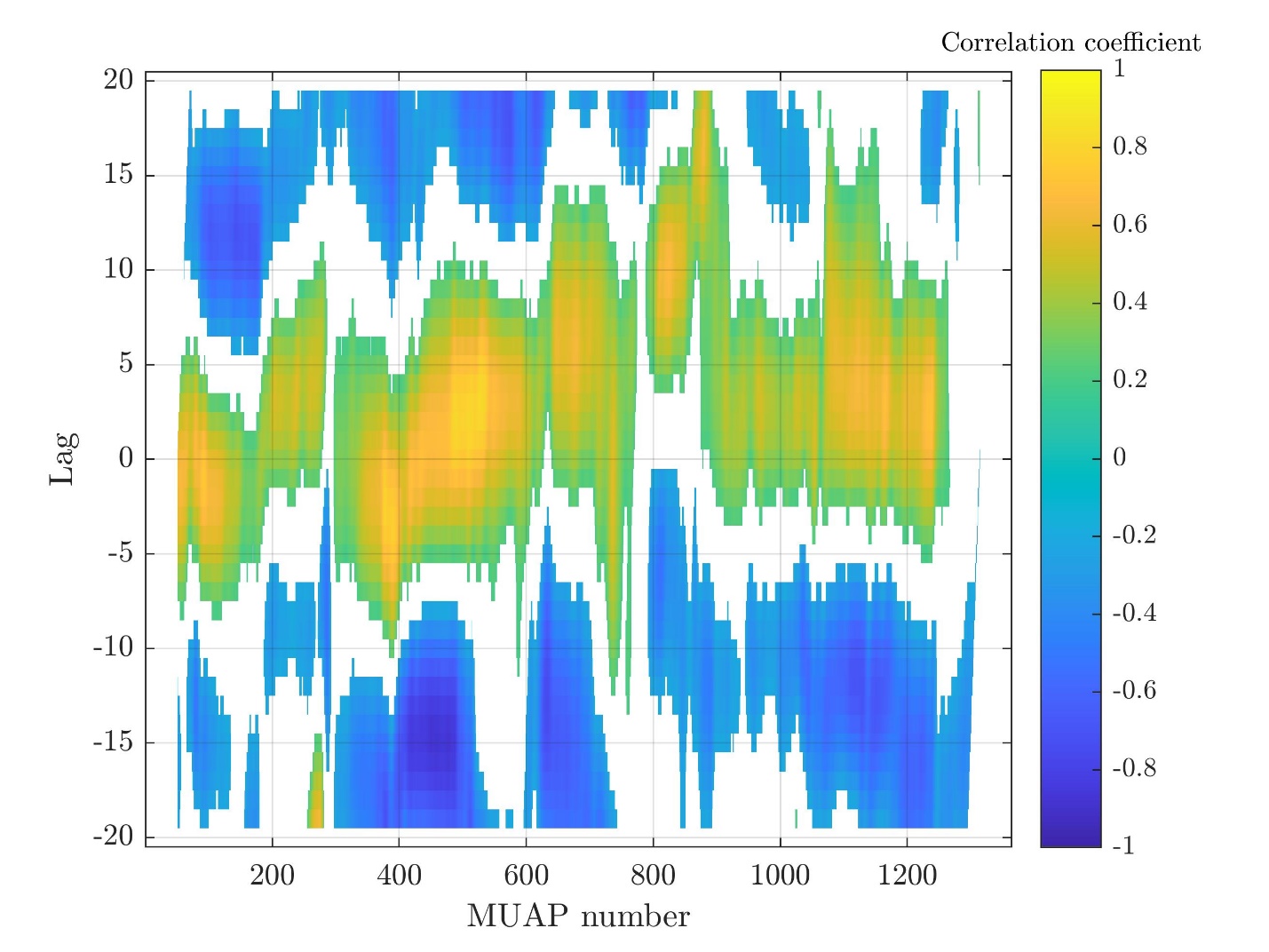

Subject 14

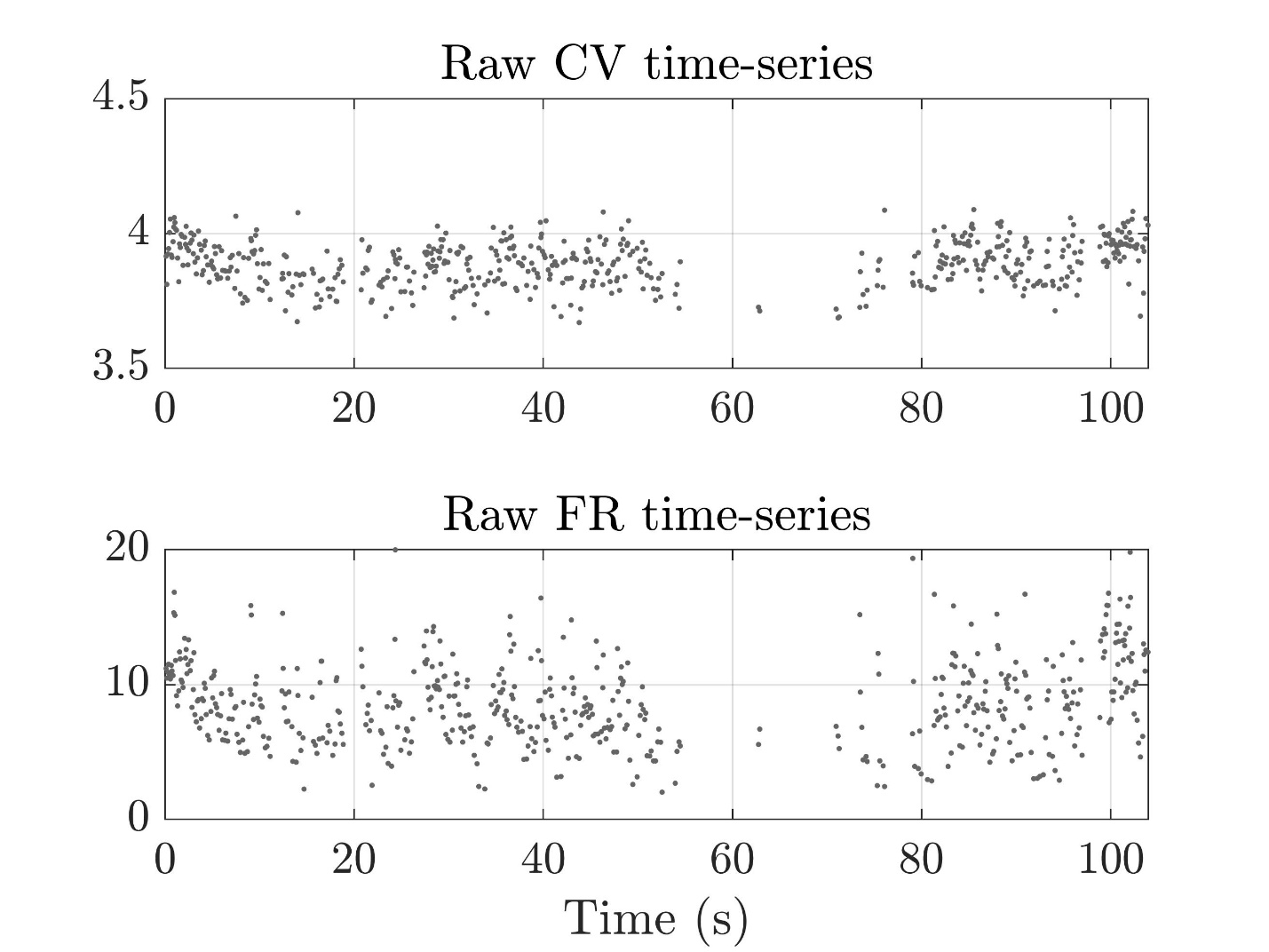

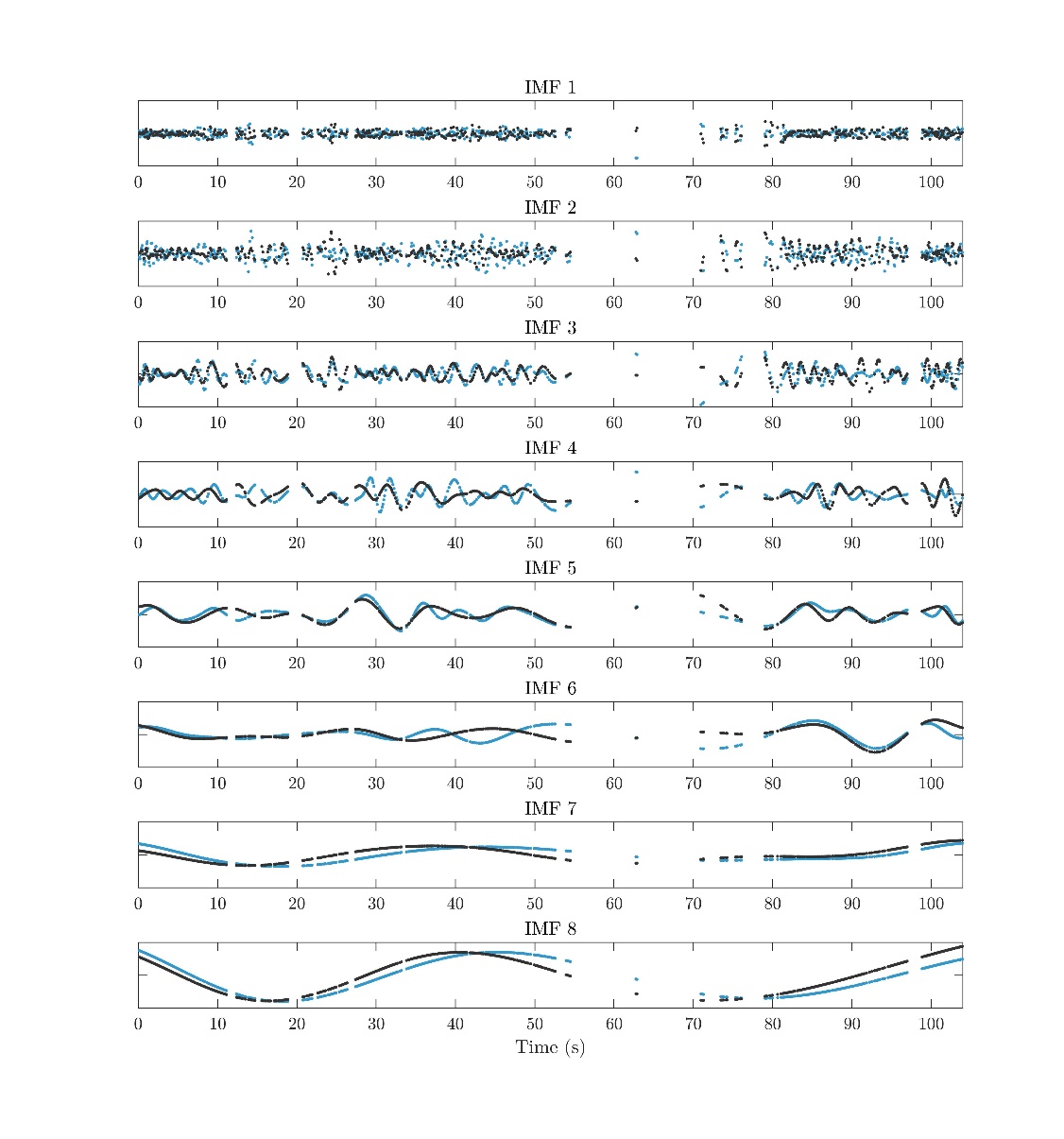

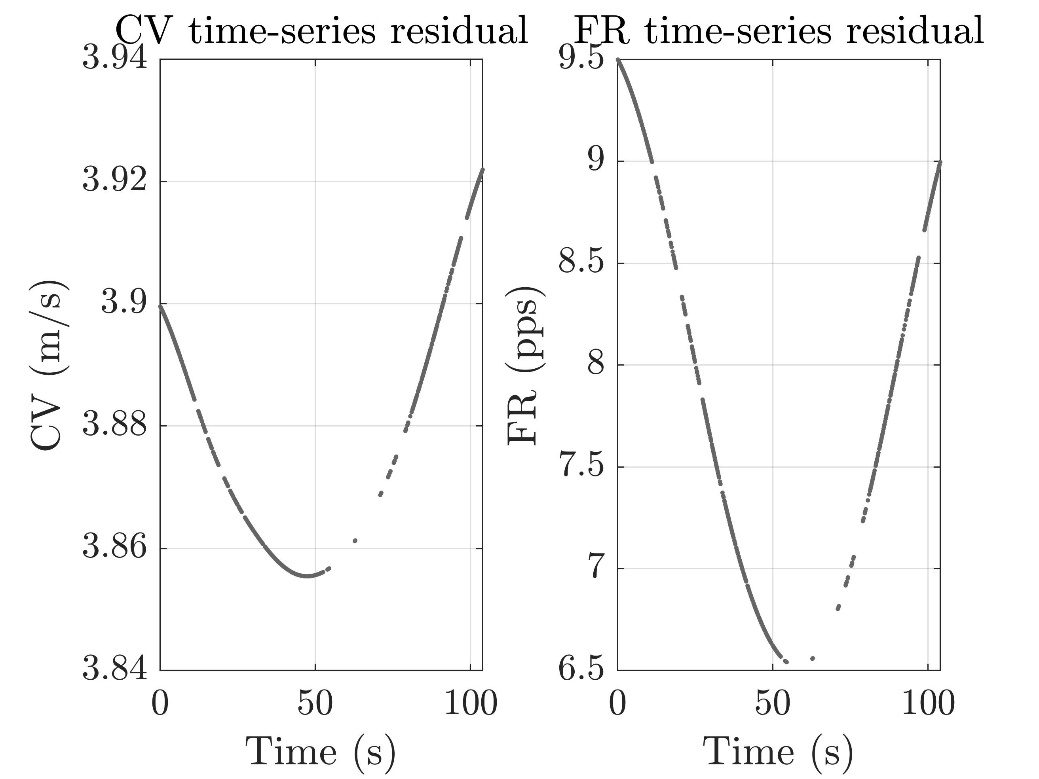

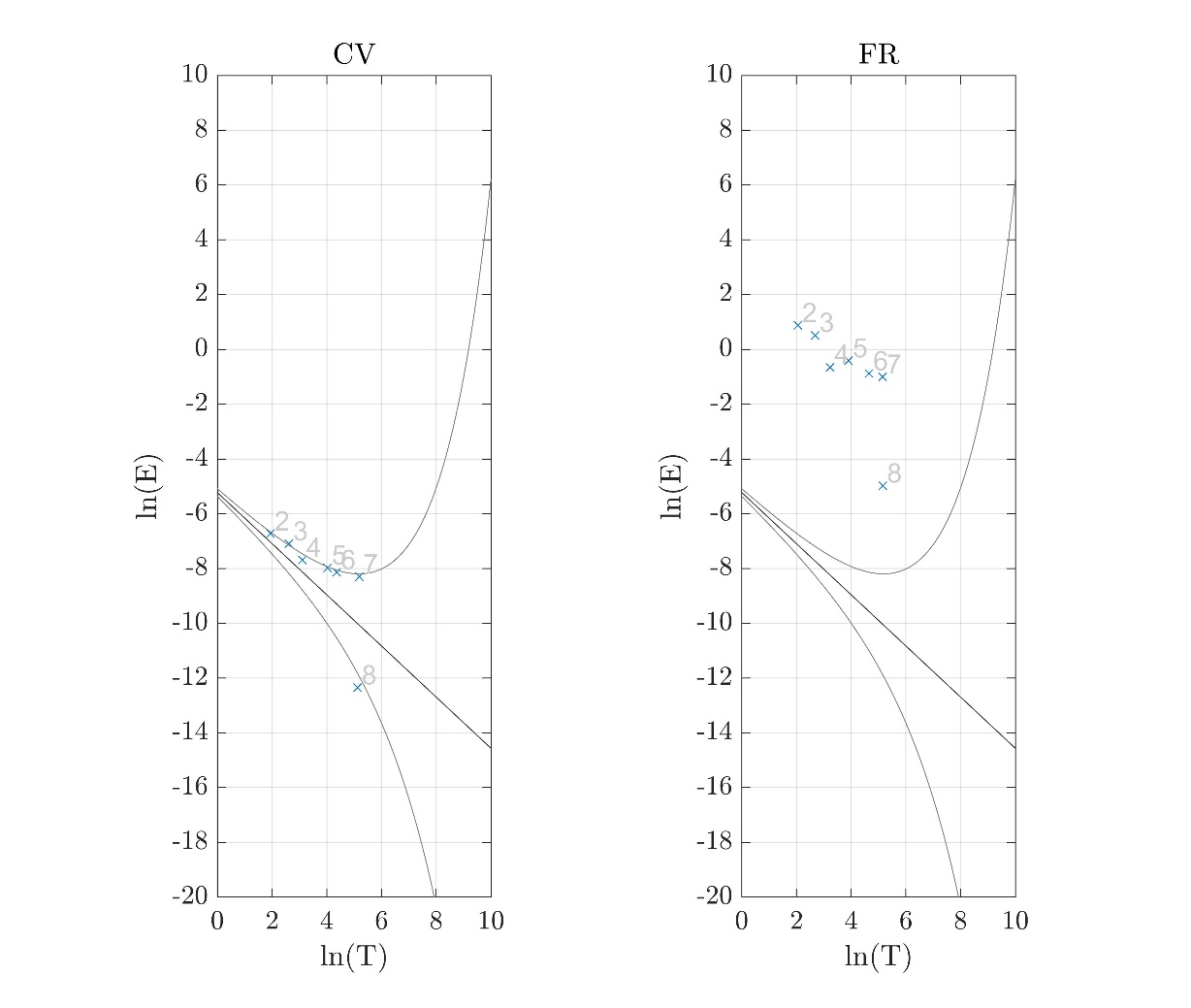

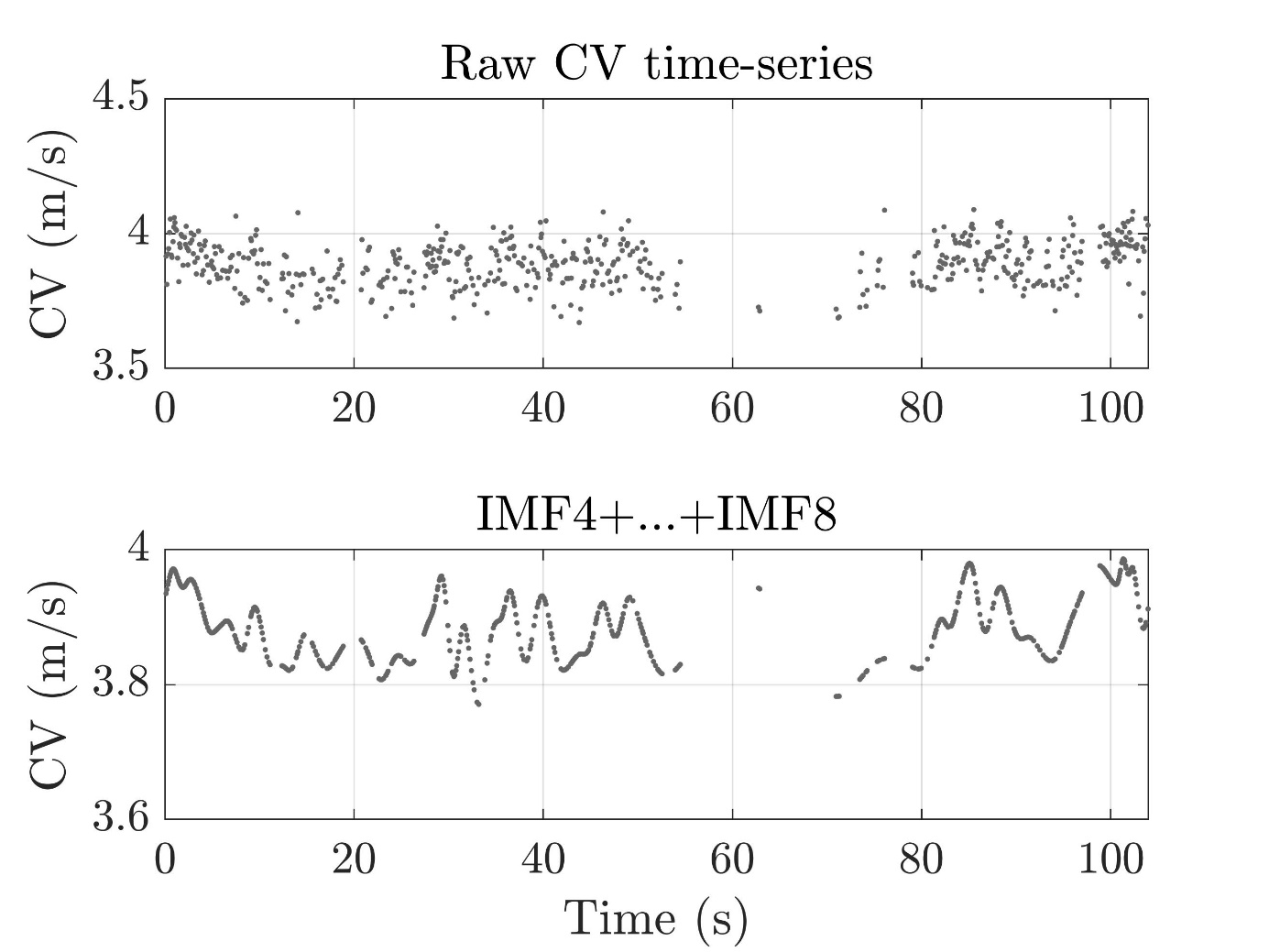

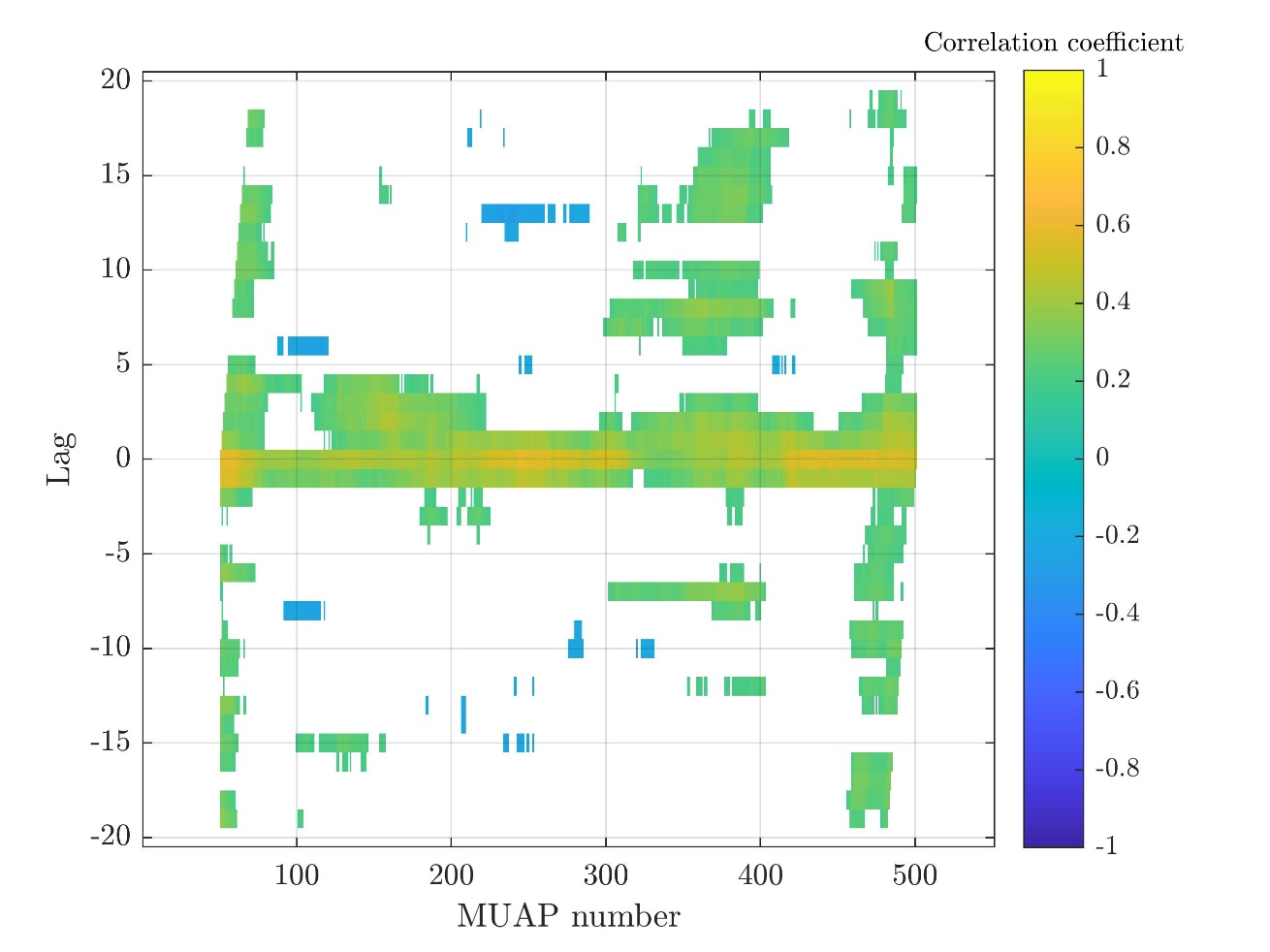

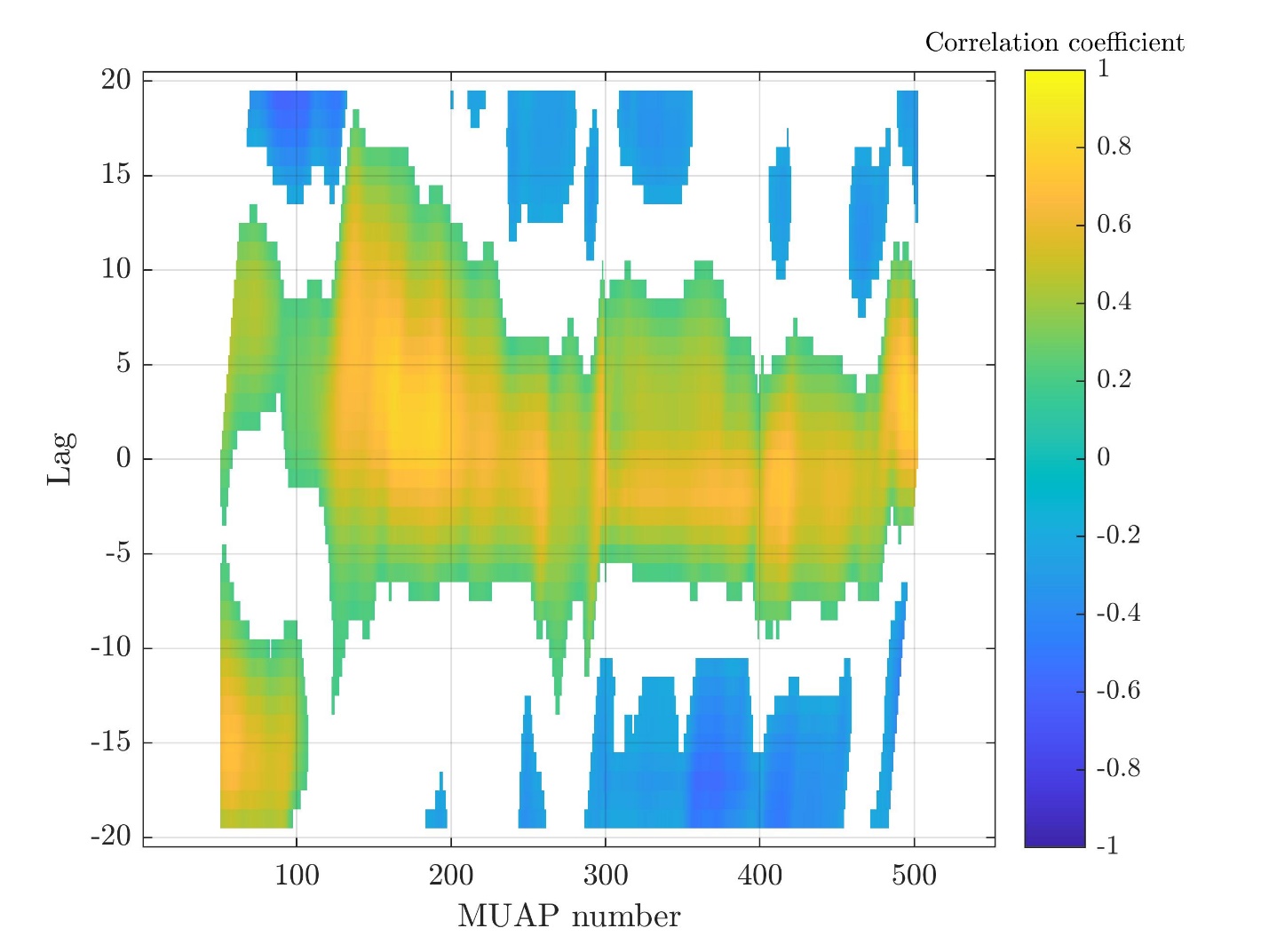

Subject 15

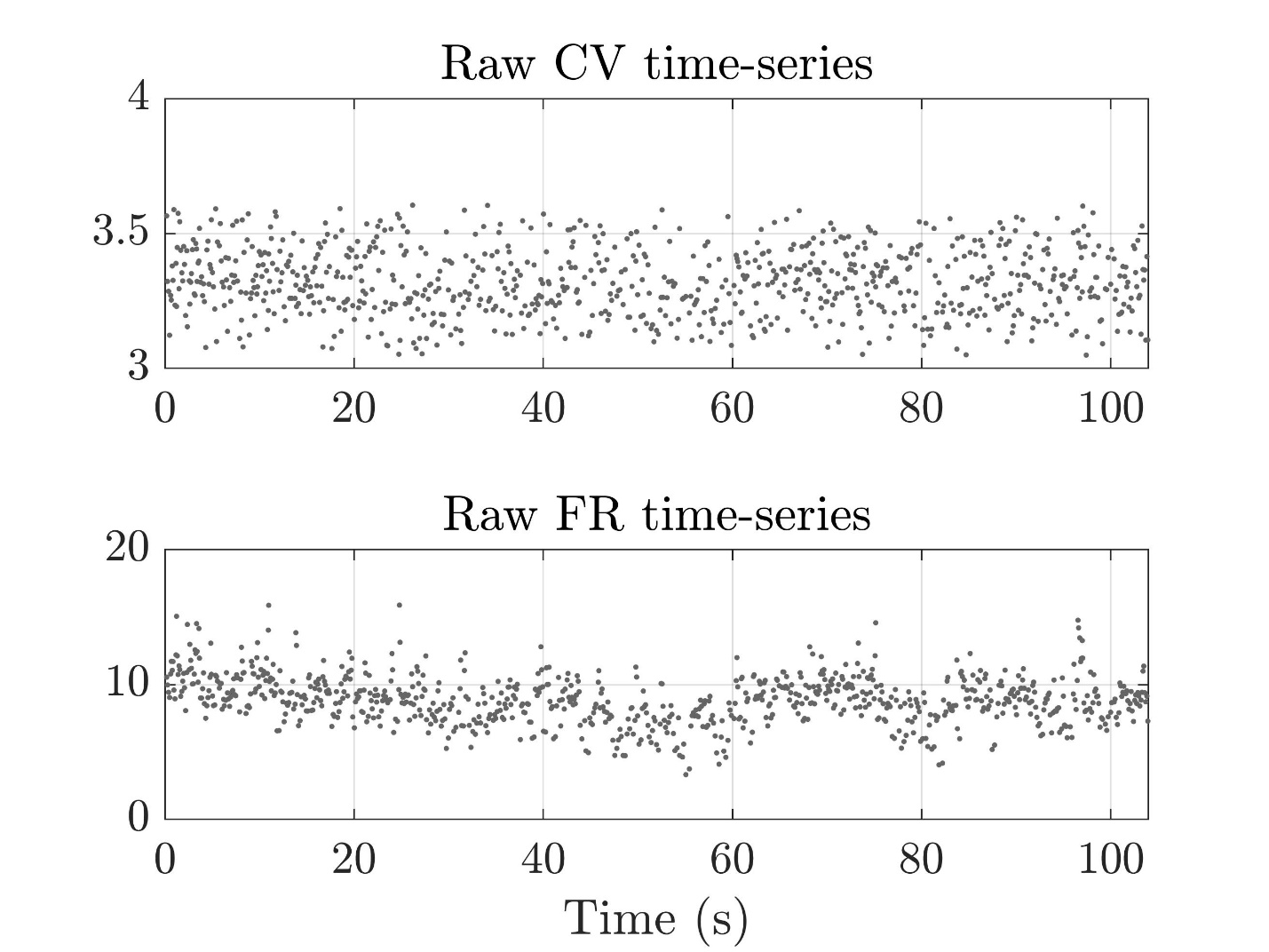

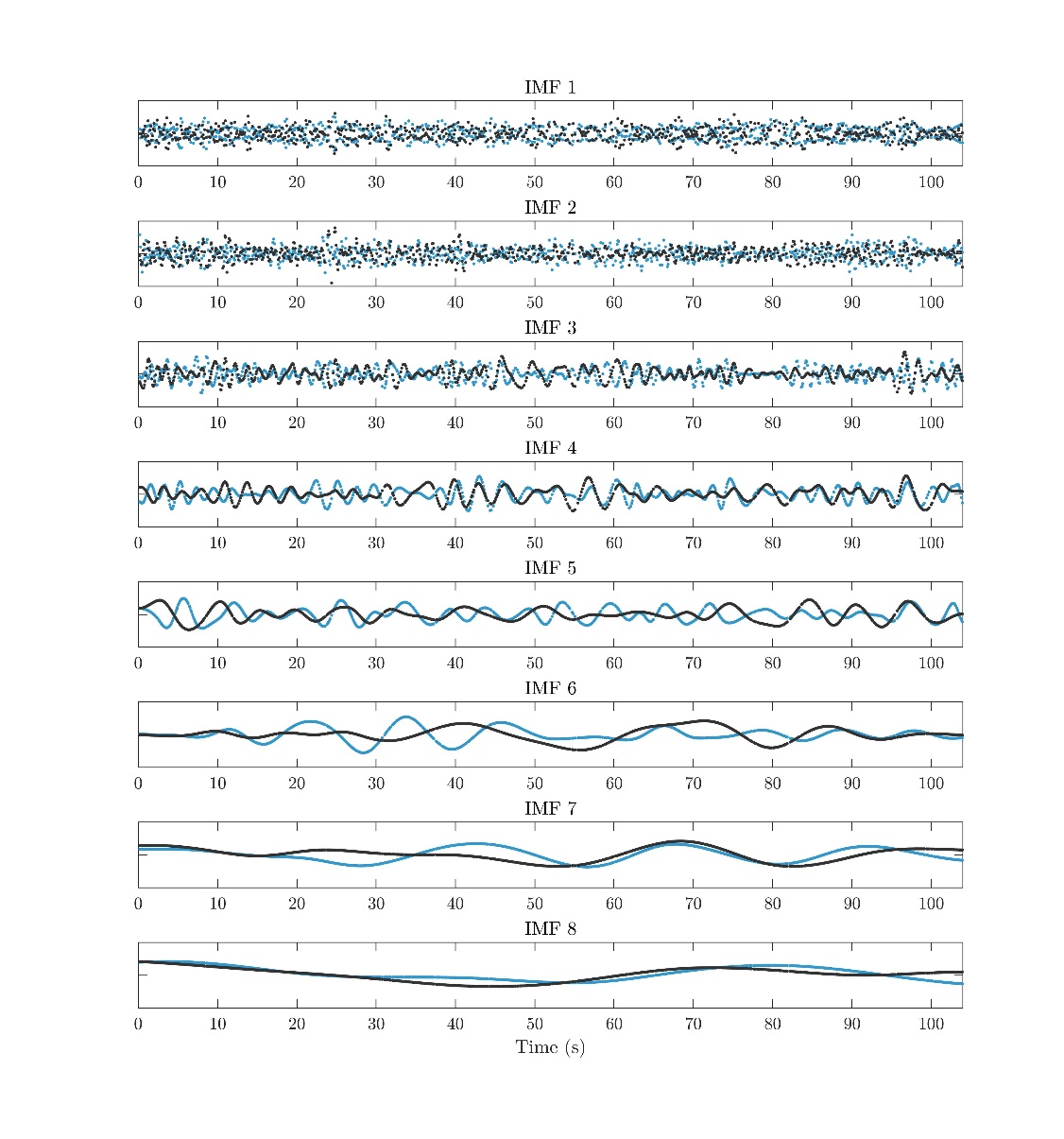

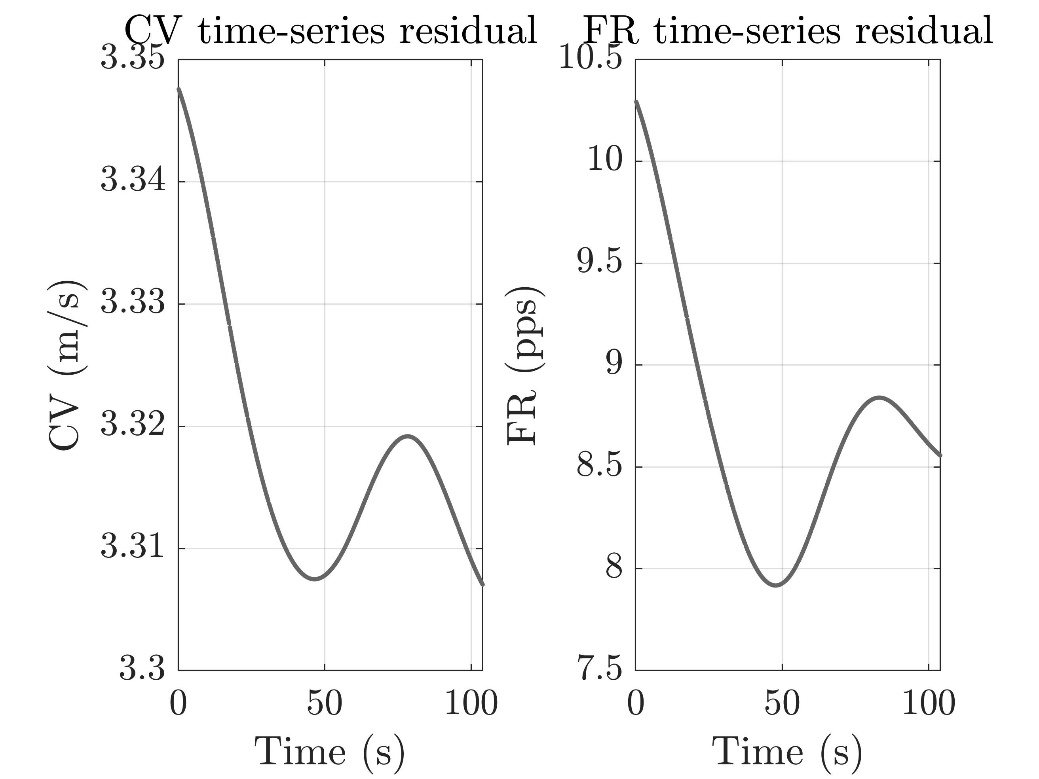

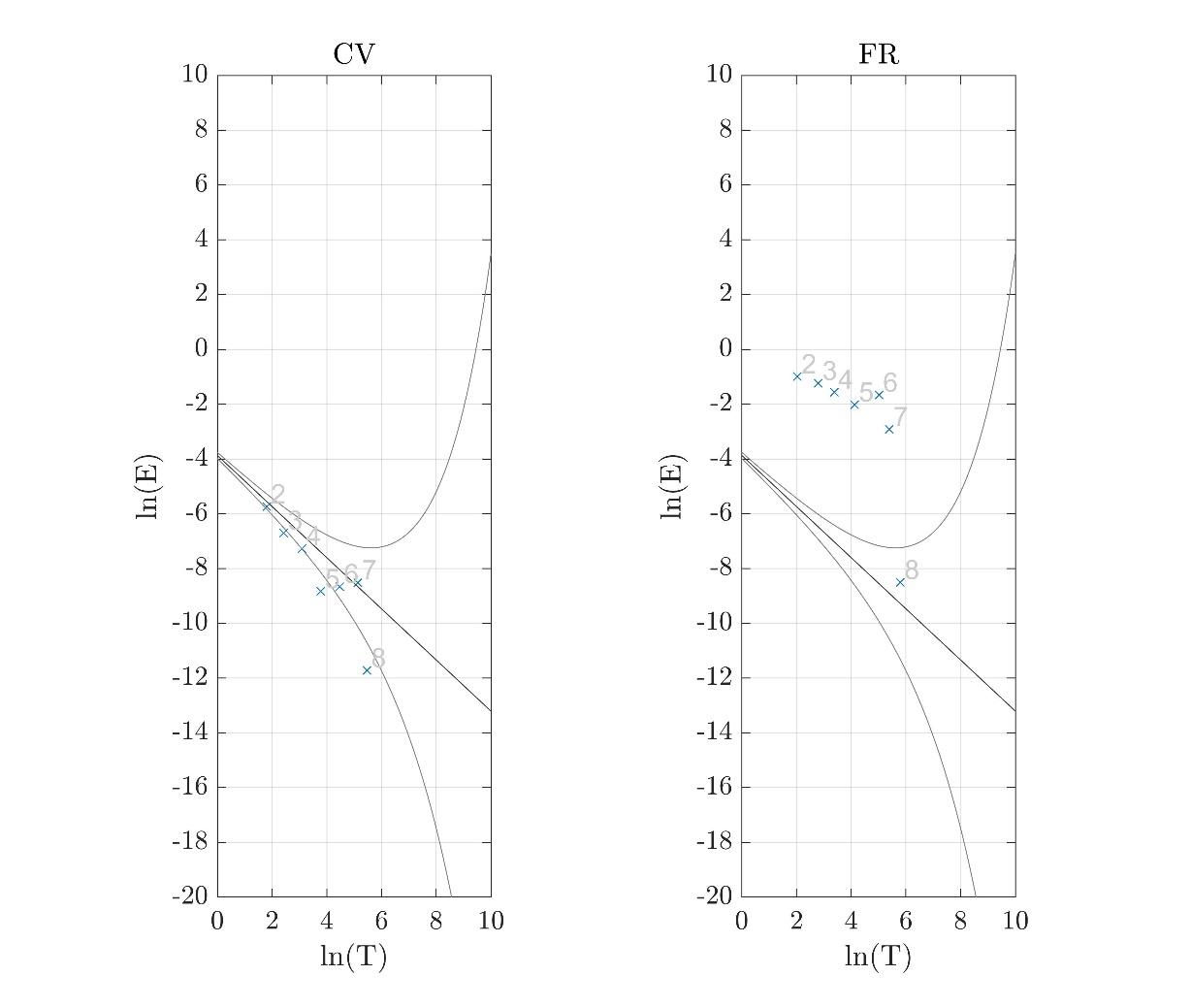

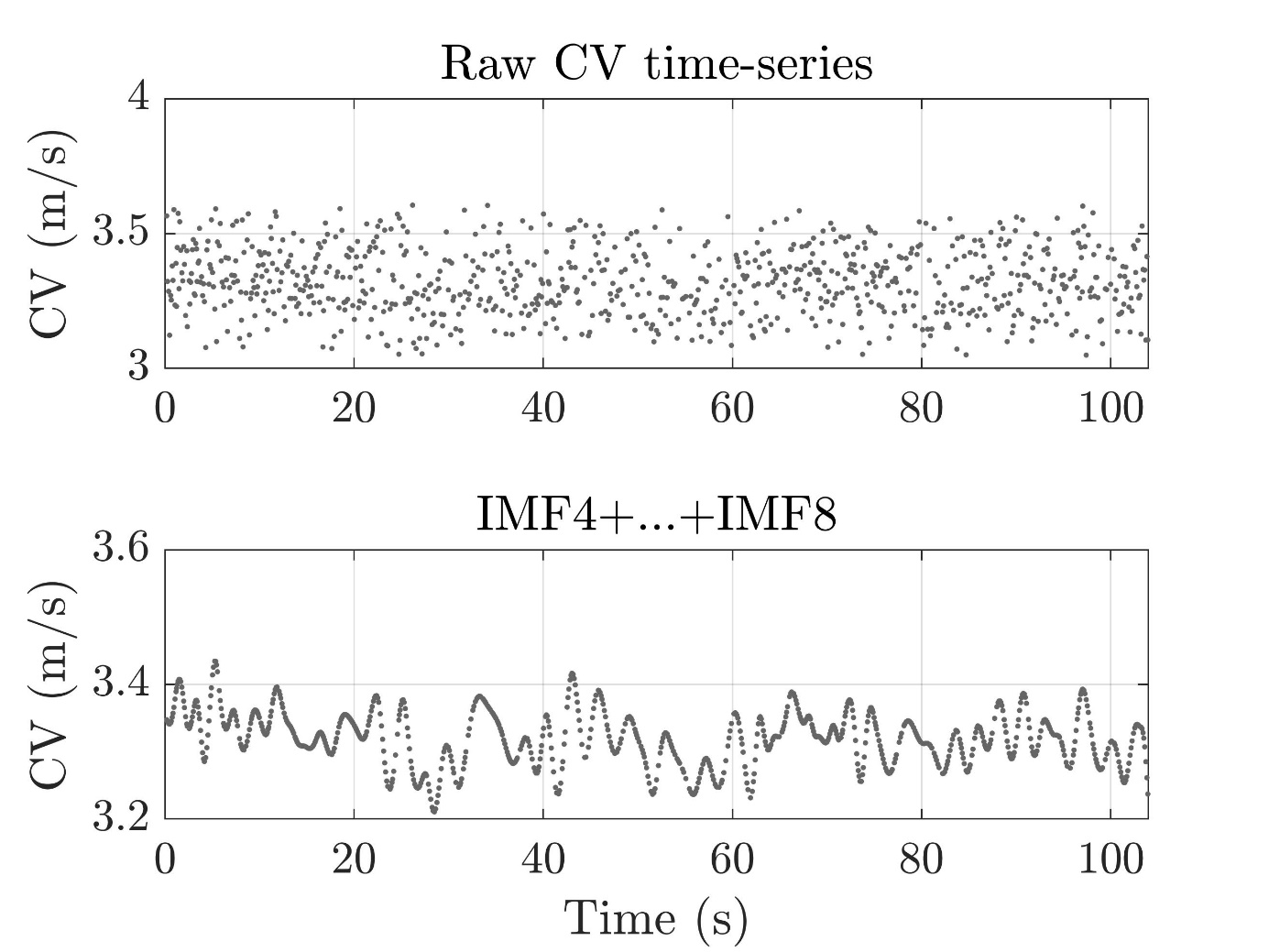

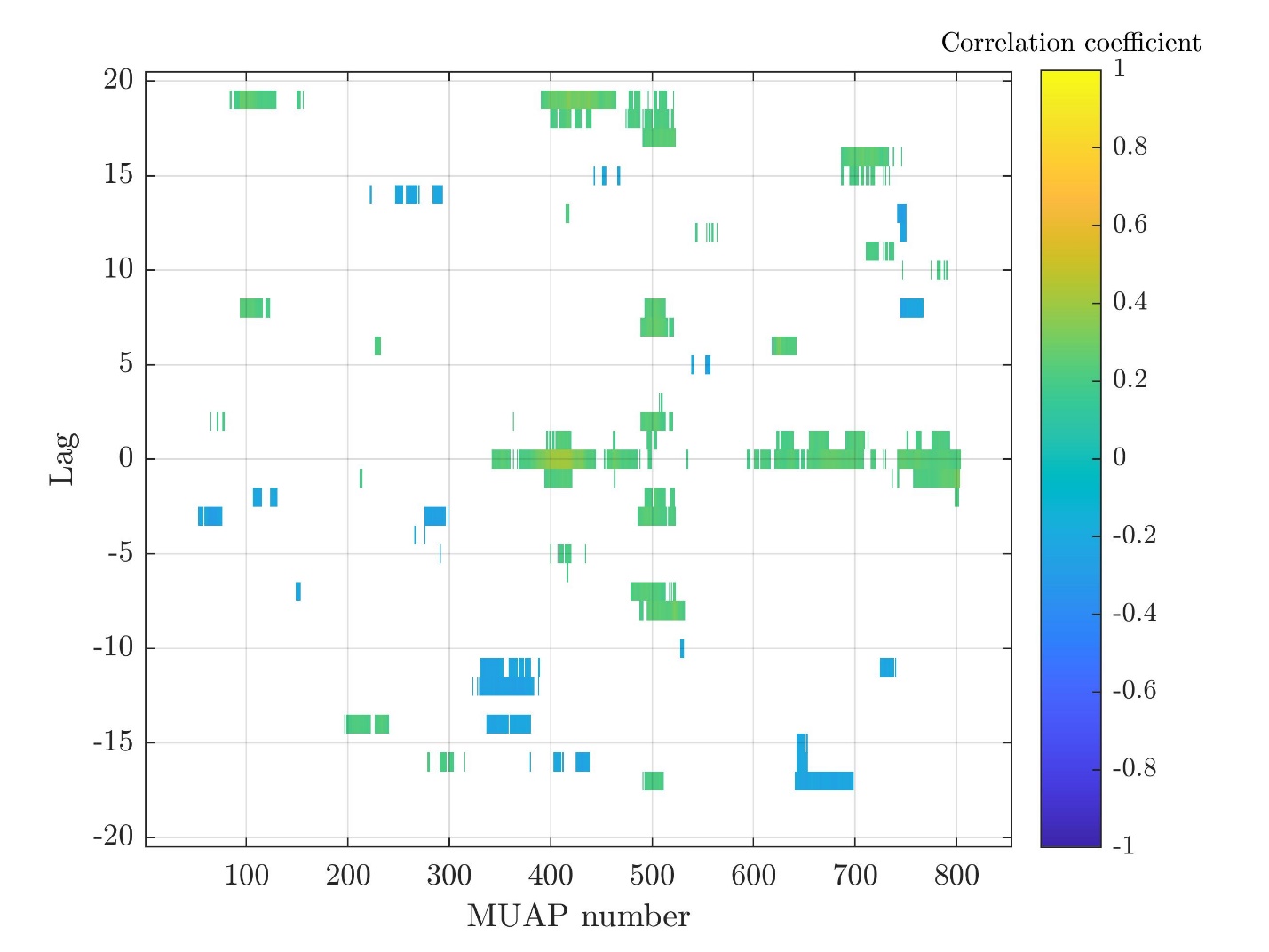

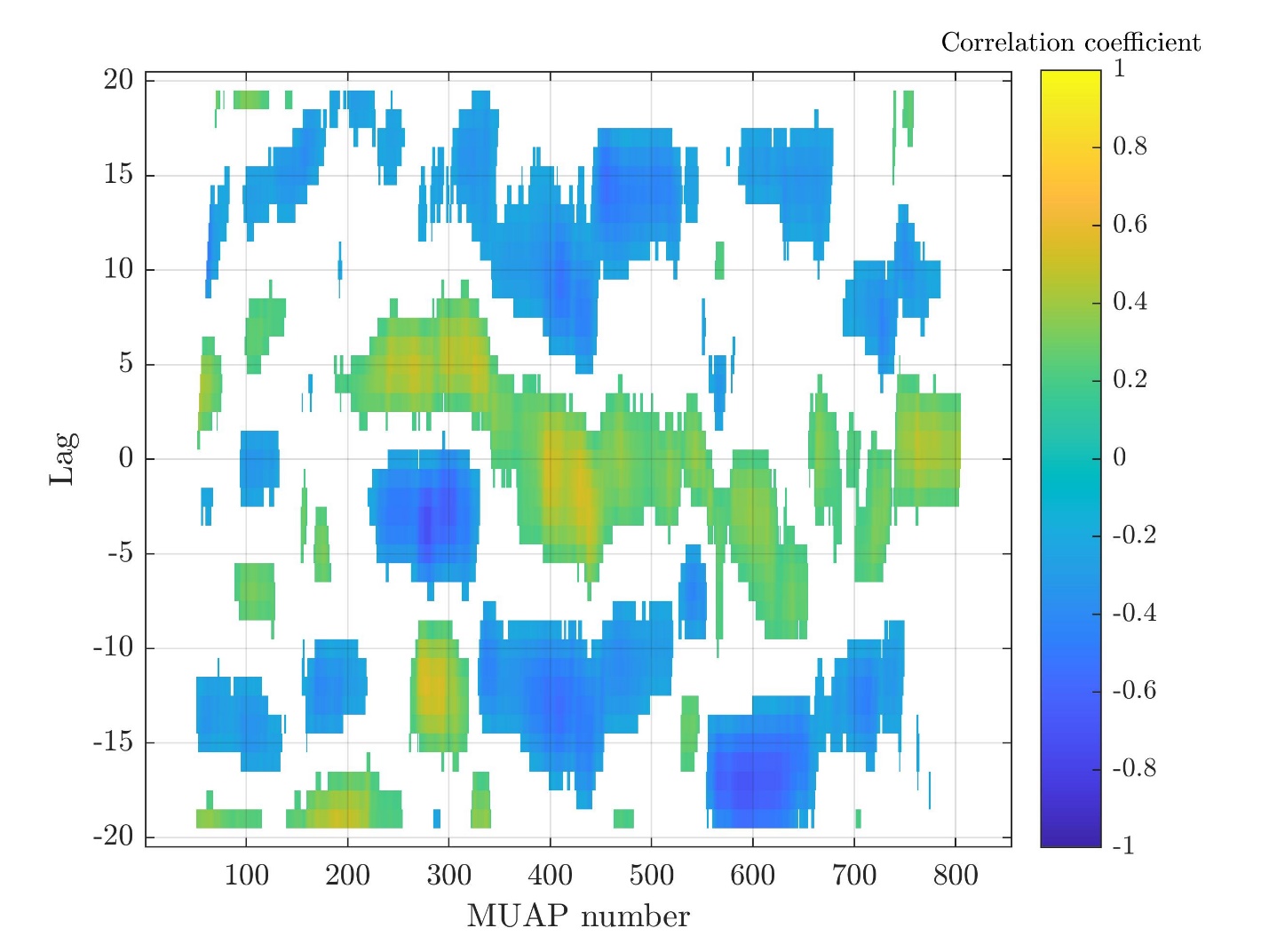
